# Iron-catalyzed antagonism between NO and H_2_O_2_ / HOCl causes mutual exclusivity of HOCl- and NO/peroxynitrite-mediated apoptosis-inducing signaling

**DOI:** 10.1101/2022.07.03.498603

**Authors:** Georg Bauer

## Abstract

Membrane-associated NADPH oxidase of malignant cells drives intercellular apoptosis-inducing HOCl- and NO/peroxynitrite signaling. Cells from late stages of oncogenesis, i. e. bona fide tumor cells, interfere with these signaling pathways through expression of membrane-associated catalase. Transformed cells and tumor cells with inhibited catalase show mutual exclusivity of apoptosis-inducing NO/peroxynitrite and HOCl signaling. H_2_O_2_ is the substrate for HOCl synthesis by DUOX-coded peroxidase. However, at high concentrations of H_2_O_2_, HOCl signaling is abrogated through the reaction between excess H_2_O_2_ and HOCl. Experimental increase in the NO concentration through addition of NO donors, induction of NO synthase (NOS), addition or the NOS substrate arginine, or inhibition of NO dioxygenase(NOD) leads to the inhibition of HOCl signaling, but also to abrogation of H_2_O_2_-mediated interference with HOCl signaling. Vice versa, an increase in extracellular H_2_O_2_ through increasing the cell density, gamma irradiation or addition of glucose oxidase inhibits NO/peroxynitrite signaling. These mutual reactions between NO and H_2_O_2_/HOCl are catalyzed by an iron-mediated reaction cycle, which allows consumption of NO by H_2_O_2_/HOCl and vice versa. These reactions explain the preferences of tumor cell lines for NO/peroxynitrite or HOCl signaling after inhibition of their protective catalase, as well as the differential expression of these pathways dependent on the degree of catalase inhibition. The understanding of these reactions allows to rationally modulate quality and strength of ROS/RNS-dependent apoptosis-inducing signaling pathways of malignant cells. The knowledge about these reactions should allow to optimize strategies for tumor therapy that are based on reactivation of intercellular ROS/RNS-dependent apoptosis-inducing signaling of tumor cells.

**Graphical abstract:** 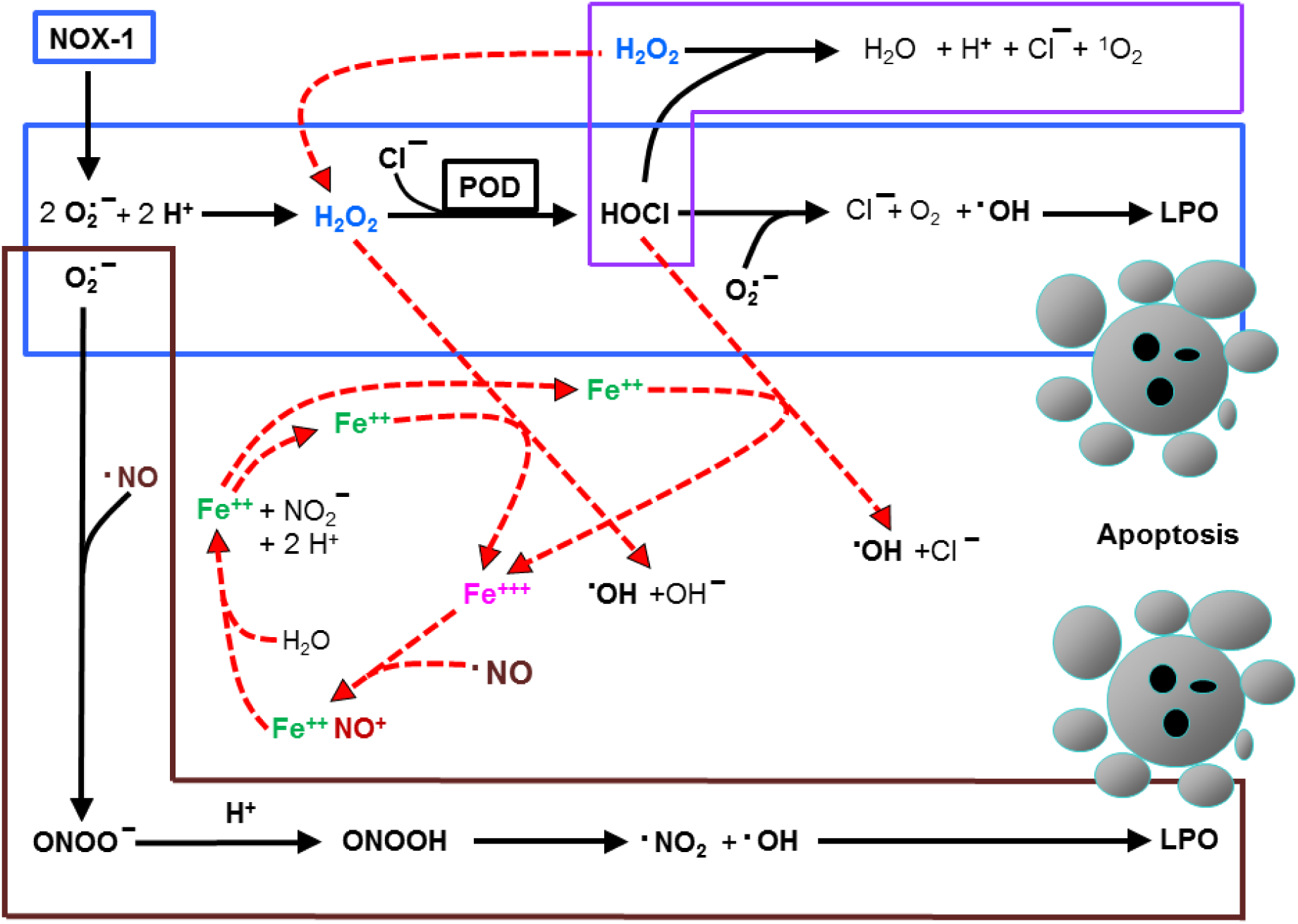

**Highlights:**

• NADPH oxidase and peroxidase drive apoptosis-inducing HOCl signaling
• NADPH oxidase and NO synthase drive apoptosis-inducing NO/peroxynitrite signaling
• H_2_O_2_ establishes and interferes with HOCl signaling, dependent on its concentration
• NO inhibits HOCl signaling and abrogates H_2_O_2_-dependent inhibition of HOCl signaling
• H_2_O_2_ interferes with NO/peroxnitrite signaling
• Mutual interactions of NO and H_2_O_2_ are catalysed by an iron-mediated reaction cycle

## Introduction

Malignant transformation depends on the expression of membrane-associated NADPH oxidase (NOX), leading to the generation of extracellular superoxide anions [1–7]. Their dismutation product H_2_O_2_ establishes an autocrine loop of proliferation stimulation and maintenance of the transformed state. However, extracellular superoxide anions also drive two intercellular ROS/RNS-dependent apoptosis-inducing signaling pathways, i. e. the HOCl and the NO/peroxynitrite signaling pathway [6–11]. Both pathways are based on multiple reactions between radical and nonradical reactive oxygen and nitrogen species (ROS/RNS). They induce apoptosis selectively in malignant cells [6, 8, 9, 10, 11, 12, 13] (Figure 1). Both pathways are driven by extracellular superoxide anions and are finalized by the generation of hydroxyl radicals. Selective apoptosis-inducing signaling in both pathways is established in close vicinity to the membrane of NADPH oxidase 1 (NOX1)- expressing malignant target cells, due to the relatively short free diffusions path length of superoxide anions and hydroxyl radicals [14, 15]. As a consequence, hydroxyl radicals cause lipid peroxidation, followed by execution of the mitochondrial pathway of apoptosis [16]. This leads to selective elimination of transformed cells [6, 17].

**Figure 1.**
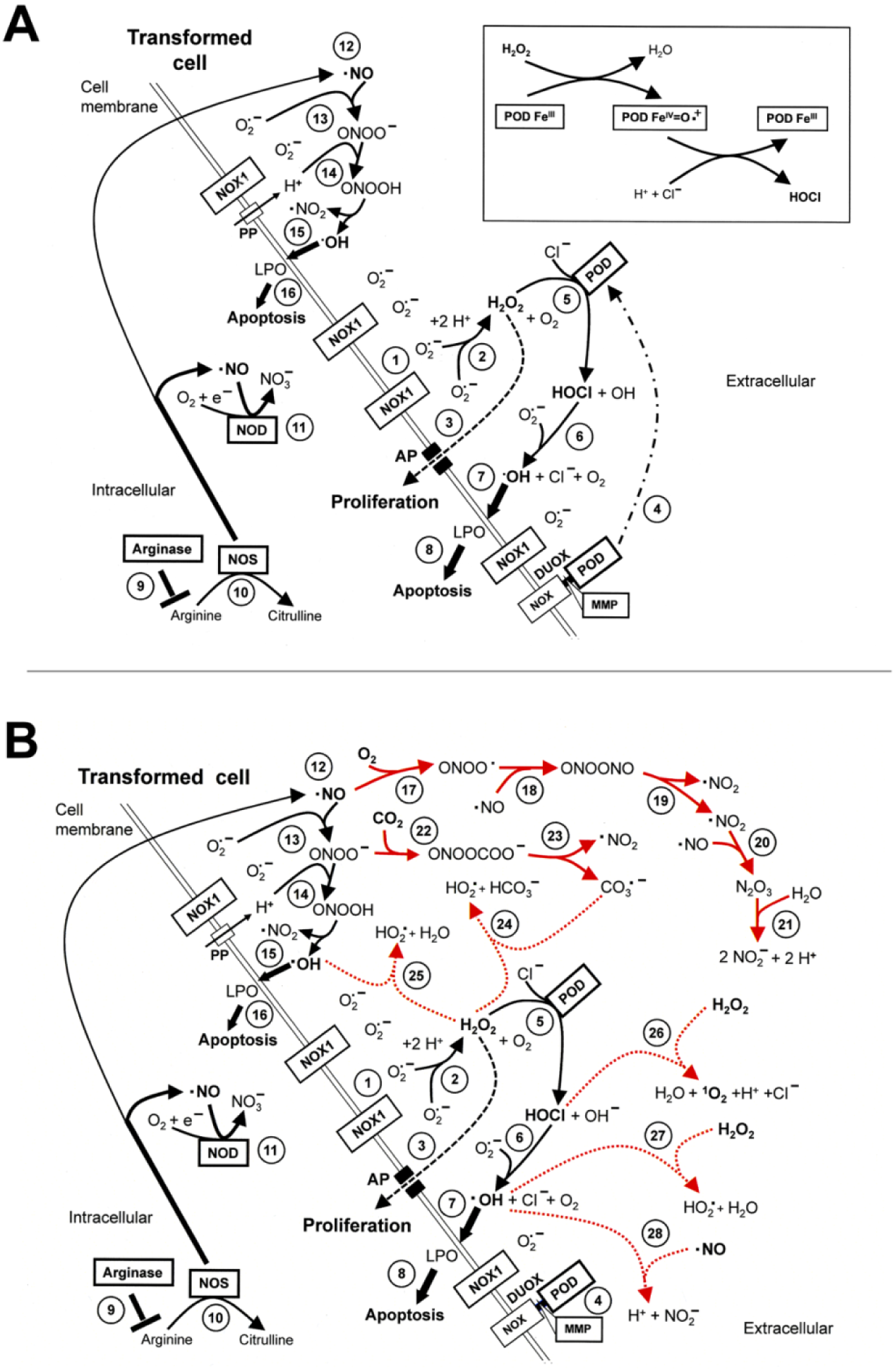
HOCl- and NO/peroxynitrite-mediated apoptosis-inducing signaling of transformed cells. A. Basic reactions. HOCl signaling: Membrane-associated NOX1 (#1) is a hallmark of transformed cells and generates extracellular superoxide anions (#2). These dismutate to form H_2_O_2_ which may either enter the cell through aquaporins (#3) or can be utilized by the peroxidase domain of DUOX 8POD) (#4) to generate HOCl (#5). The reaction of HOCl with superoxide anions (#6) leads to the generation of hydroxyl radicals (#7) in close vicinity to the cell membrane. Lipid peroxidation induced by hydroxyl radicals leads to lipid peroxidation (LPO) (#8) and triggers the mitochondrial pathway of apoptosis. NO/peroxynitrite signaling: Arginase (#9) controls the concentration of arginine, which is the substrate for NO synthase (#10). NO dioxygenase (NOD) consumes a significant part of free NO (#11). The remaining NO has a chance to pass the cell membrane (#12) and generate peroxynitrite after reaction with superoxide anions (#13). The protonation of peroxynitrite through protons derived from proton pumps in the membrane generates peroxynitrous acid (#14) which spontaneously decomposes into NO_2_ and apoptosis-inducing hydroxyl radicals (#15, 16). These two pathways do not act in parallel, but rather sequentially, as described in this manuscript. B. Side reactions. The HOCl- (#1-8) and the NO/peroxynitrite signaling pathway (#9-16) are subject to several side reactions of ROS/RNS that act towards their efficiency. The oxidation of NO by molecular oxygen (#17-21) decreases the concentration of NO available for peroxynitrite formation. The reaction between peroxynitrite and CO_2_ (#22) is competing with protonation of peroxynitrite (#14). Only in close vicinity to proton pumps the less efficient protonation reaction of peroxynitrite takes place. Nitrosoperoxycarboxylate (ONOOCOO**^─^**) resulting from reaction #22 is spontaneously decomposing into NO_2_ and carbonate radicals (#23). These may interact with H_2_O_2_ according to reaction # 24, with a potentially negative effect on HOCl signaling. The reaction of hydroxyl radicals with H_2_O_2_ (#25) might interfere with lipid peroxidation through the NO/peroxynitrite pathway. The reactions of H_2_O_2_ with either HOCl (#26) or hydroxyl radicals (#27) has the potential to interfere with HOCl signaling. A similar inhibitory effect is conceivable for free NO (#28).

Recent experimental work revealed that the effect of lipid peroxidation is initially counteracted by intracellular glutathione/glutathione peroxidase-4. Therefore, in addition to the action of hydroxyl radicals, an influx of H_2_O_2_ through aquaporins is required to cause glutathione depletion. This then allows the initiation of the mitochondrial pathway of apoptosis [18].

As shown in Figure 1 B, signaling through the HOCl and the NO/peroxynitrite pathway is counteracted by a multitude of interfering side reactions. Importantly, close to the membrane with its proton pumps, the protonation of peroxynitrite (ONOO^─^) to peroxynitrous acid (ONOOH) is favoured and allows the subsequent generation of hydroxyl radicals [11, 13]. Whereas these site-specific reactions are essential for NO/peroxynitrite signaling, the fast reaction between peroxynitrite and CO_2_ [19, 20] distant of the membrane, can be predicted to lower the efficiency of NO/peroxynitrite signaling through consumption of peroxynitrite.

Tumor progression requires establishment of resistance towards apoptosis-inducing intercellular ROS/RNS signaling [21–25]. This is achieved through expression of membrane-associated catalase [26–30] (Figure 2). Membrane-associated catalase prevents HOCl signaling through decomposition of H_2_O_2._ It also interferes with NO/peroxynitrite signaling through oxidation of NO and decomposition of peroxynitrite [9, 10, 28, 29, 31–33]. The activity of membrane-associated catalase is a regular phenotypic feature of *bona fide* tumor cells [28]. This finding explains the mechanism of resistance against oxidative stress, which has been found by Deichman and co-workers to represent a central step during tumor progression [21–25], Bona fide tumor cells express high local concentrations of catalase on their membrane, thus ensuring tight control of ROS/RNS signaling. Transformed cells, i. e. early stages of oncogenesis, only express a very low, but detectable concentration of membrane-associated catalase on their surface. This concentration of catalase is not sufficient to actually prevent ROS/RNS-mediated apoptosis signaling of transformed cells, but may have a modulatory effect [29]. Covalent attachment of catalase to the membrane of tumor cells is mediated by transglutaminase [29. 34]. Coexpression of SOD is necessary to prevent superoxide anion-dependent inhibition of catalase [10, 30, 35-38] (Figure 2 B). Inhibition or inactivation of protective catalase of tumor cells reactivates intercellular ROS/RNS-dependent apoptosis-inducing signaling and therefore has been suggested as a novel approach for specific tumor therapy [13, 28, 30, 34, 39-46]. Conceivable techniques are inhibition of catalase by neutralizing antibodies [41], or inactivation of catalase through extracellular singlet oxygen [39, 43–45]. Singlet oxygen can either be generated by photosensitizers [39], supplied by cold atmospheric plasma directly or through the reaction between long-lived species derived from cold atmospheric plasma or plasma activated medium [30, 42–46]. The generation of singlet oxygen can also be triggered through modulation of the endogenous NO concentration in tumor cells [13, 40]. This can be achieved by increasing the concentration of arginine, inhibition of arginase, induction of NOS by interferons or inhibition of NO dioxygenase (NOD) and leads to a complex signaling process that involves the redox biology of tumor cells, based on active NOX1, NOS and FAS receptor (Bauer, 2015). In addition, inhibition of membrane-associated SOD by high concentrations of exogenous H_2_O_2_ has been shown to lead to the same sequence of biochemical steps that culminate in the generation of singlet oxygen [47].

**Figure 2.**
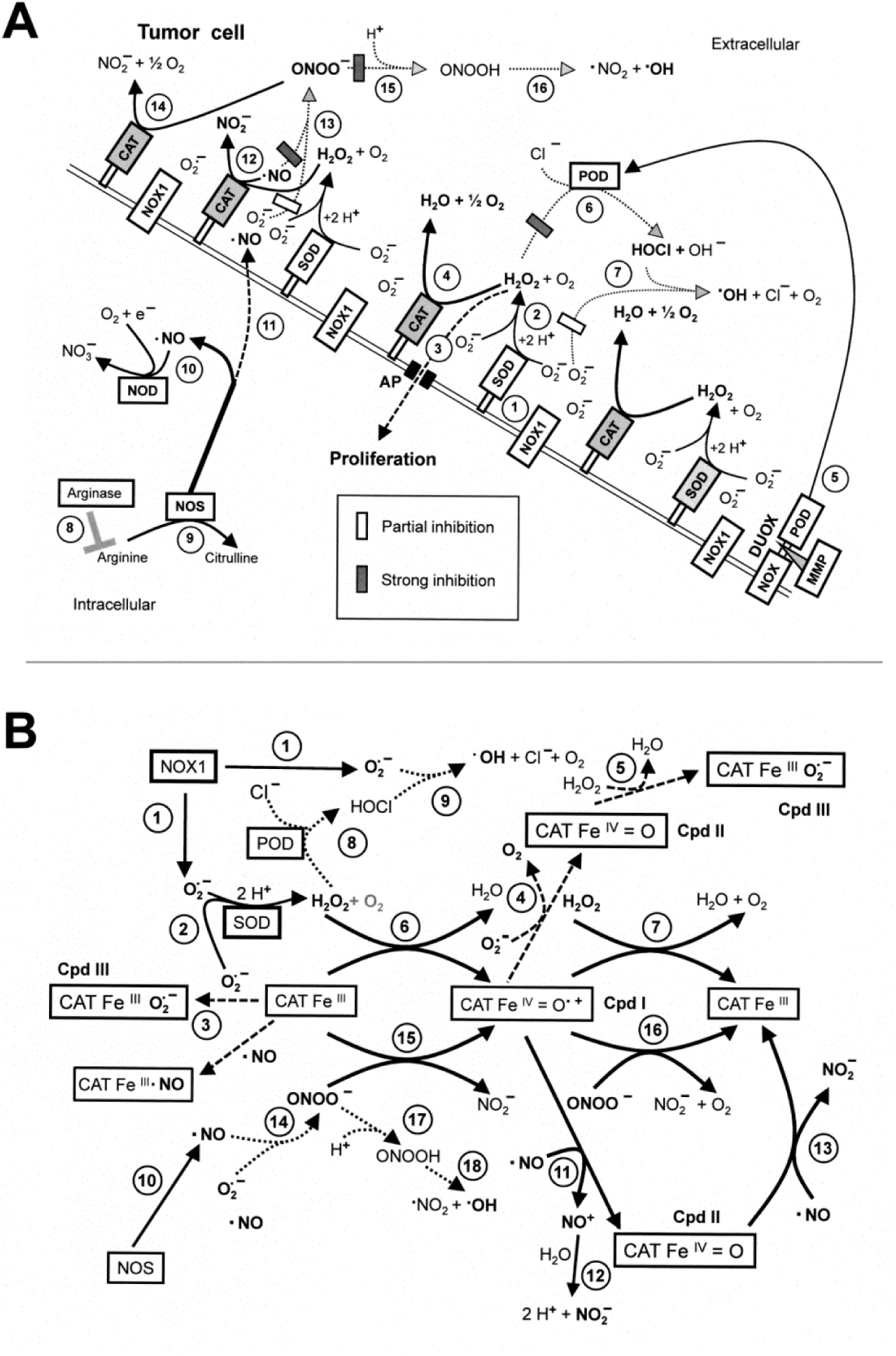
Membrane-associated catalase protects tumor cells towards exogenous ROS/RNS signaling. A. Redox-relevant elements on the membrane of tumor cells. Membrane-associated NADPH oxidase-1 (NOX-1) (#1) generates extracellular superoxide anions that may dismutate to H_2_O_2_ either spontaneously or catalysed by membrane-associated SOD (#2). A minimum of H_2_O_2_ may enter the cells through aquaporins (#3) and stimulate proliferation, whereas the majority of H_2_O_2_ is decomposed by catalase (#4). This prevents HOCl signaling (#5-7). The interaction between arginine, arginase, NOS and NOD (#8-10) determines the concentration of NO that passes through the membrane (#11). NO can be oxidized by compound I of catalase (#12). Compound I is formed through a one-step reaction between catalase and H_2_O_2_. The oxidation of NO interferes with peroxynitrite formation through NO/superoxide anion interaction (#13). Eventually formed peroxynitrite is decomposed by catalase in a two-step reaction (#14). Oxidation of NO and decomposition of peroxynitrite prevent the formation of peroxynitrous acid and hydroxyl radicals (#15, 16). This figure shows that catalase ensures tight control towards both signaling pathways. In contrast, the activity of SOD is not sufficient to remove all free superoxide anions, but sufficient to lower the superoxide anion concentration below a value where superoxide-mediated inhibition of catalase would interfere with catalase-mediated protetection. B. Multiple enzymatic activities of catalase. NOX1-derived superoxide anions (#1) dismutate to H_2_O_2_through the action of SOD (#2). This prevents superoxide anion-mediated inhibition of catalase through reactions #3-5. H_2_O_2_ interacts with native catalase (CAT Fe^+++^) (#6), resulting in the formation of compound I (CATFe^IV^=O**^●+^).** Compound I reacts with a second molecule of H_2_O_2_, resulting in H_2_O and O_2_ (#7). The decomposition of H_2_O_2_ by catalase prevents HOCl signaling (#8, 9). NO is generated by NOS (#10). NO can be oxidized by compound I of catalase in a two-step reaction (#11-13). This prevents NO-mediated inhibition of catalase through formation of a CATFe^III**●**^NO adduct. Alternatively, NO may interact with superoxide anions and generate peroxynitrite (#14), which is decomposed by catalase in a two-step reaction (#15, 16). This prevents the formation of peroxynitritous acid and apoptosis-inducing hydroxyl radicals (#17, 18).

Antagonistic as well as synergistic interactions between NO and H_2_O_2_ have been recognized by several groups. Particularly David Wink and his group have shown that NO can protect mammalian cells towards cell death induction by H_2_O_2_ [48], but that on the other hand NO can enhance H_2_O_2_-mediated antibacterial effects [49]. Prevention of toxic effects of H_2_O_2_ through reaction of NO with hydroxyl radicals derived from Fenton chemistry (^●^OH + ^●^NO → H^+^ + NO ^─^) is one potential mechanism of protection by NO [50–52], whereas the enhancing effect of NO on H_2_O_2_-mediated antibacterial effects involves several intercalated pathways.

Previous work from our group has shown that the NO/peroxynitrite and the HOCl pathway act sequentially, rather than additively [28]. Their players seem to have antagonistic potential, but also can act in a protagonistic mode under certain conditions. As H_2_O_2_ is the substrate for HOCl-generating peroxidase, it is one of the rate-limiting factors for HOCl signaling. Excess H_2_O_2_, however, may consume HOCl and therefore antagonize HOCl signaling [10, 27, 28, 53, 54]. In addition, H_2_O_2_ exerts a strong interfering effect on NO/peroxynitrite signaling [9, 28, 55]. On the other side, NO strongly interferes with HOCl signaling, but also may rescue HOCl signaling from inhibition by excess H_2_O_2_. NO/peroxynitrite signaling was enhanced by the increase in the NO concentration [9]. We therefore endeavoured to study the multiple NO/H_2_O_2_ interactions during intercellular signaling, to define the hierarchy of possible interactions, and to unravel the underlying mechanisms.

## Materials and Methods

### Materials

L-Arginine, the NADPH oxidase inhibitor 4-(2-aminoethyl-benzenesulfonyl fluoride (AEBSF), the fast decaying NO donor Diethylamino NONOate (DEA NONOate), the slowly decaying NO donor sodium nitroprusside (SNP), the catalase inhibitior 3-aminotriazole (3-AT), the iron chelator deferoxamine (DFO), human interferon gamma, the hydroxyl radical scavenger mannitol, manganese-containing superoxide dismutase (MnSOD) from *E. coli*, the NOS inhibitor N-omega-nitro-L-arginine methylester hydrochloride (L-NAME), the HOCl scavenger taurine, catalase from bovine liver, glucose oxidase (GOX), the matrix metalloprotein inhibitor galardin, the flavonoids quercetin, delphinidin, peonidin, pelargonin, xanthohumol, neutralizing monoclonal antibodies against human catalase (clone CAT-505, mouse, IgG1) and control antibodies directed against laminin were obtained from Sigma (Schnelldorf, Germany).

The mechanism-based peroxidase inhibitor 4-aminobenzoyl hydrazide (ABH) was purchased from Acros Organics, Geel, Belgium. Peroxynitrite the peroxynitrite decomposition catalyst 5-, 10-, 15-, 20-Tetrakis(4-sulfonatophenyl)porphyrinato iron(III) chloride [FeTPPS) and the catalase mimetic EUK-8 [manganese N,Ń-bis(salicylidiene)ethylenediamine chloride] were obtained from Calbiochem (Merck Biosciences GmbH, Schwalbach/Ts, Germany). The catalase mimetic EUK-134 (chloro[[2,2 -[1,2-ethanediylbis[(nitrilo-κN)methylidyne]]bis[6-methoxyphenolato-κO]]]-manganese) a product of Cayman, was obtained from Biomol (Hamburg, Germany).

TGF-beta1 was purified from human platelets as recently described [56]. Itraconazol (Sempera) was obtained from Janssen-Cilag, Neuss, Germany. The sites of action of the signaling inhibitors are illustrated in Figure 3 and in references [18, 57]. A list of references related to the specificity of these inhibitors can be found in references [18, 26].

**Figure 3.**
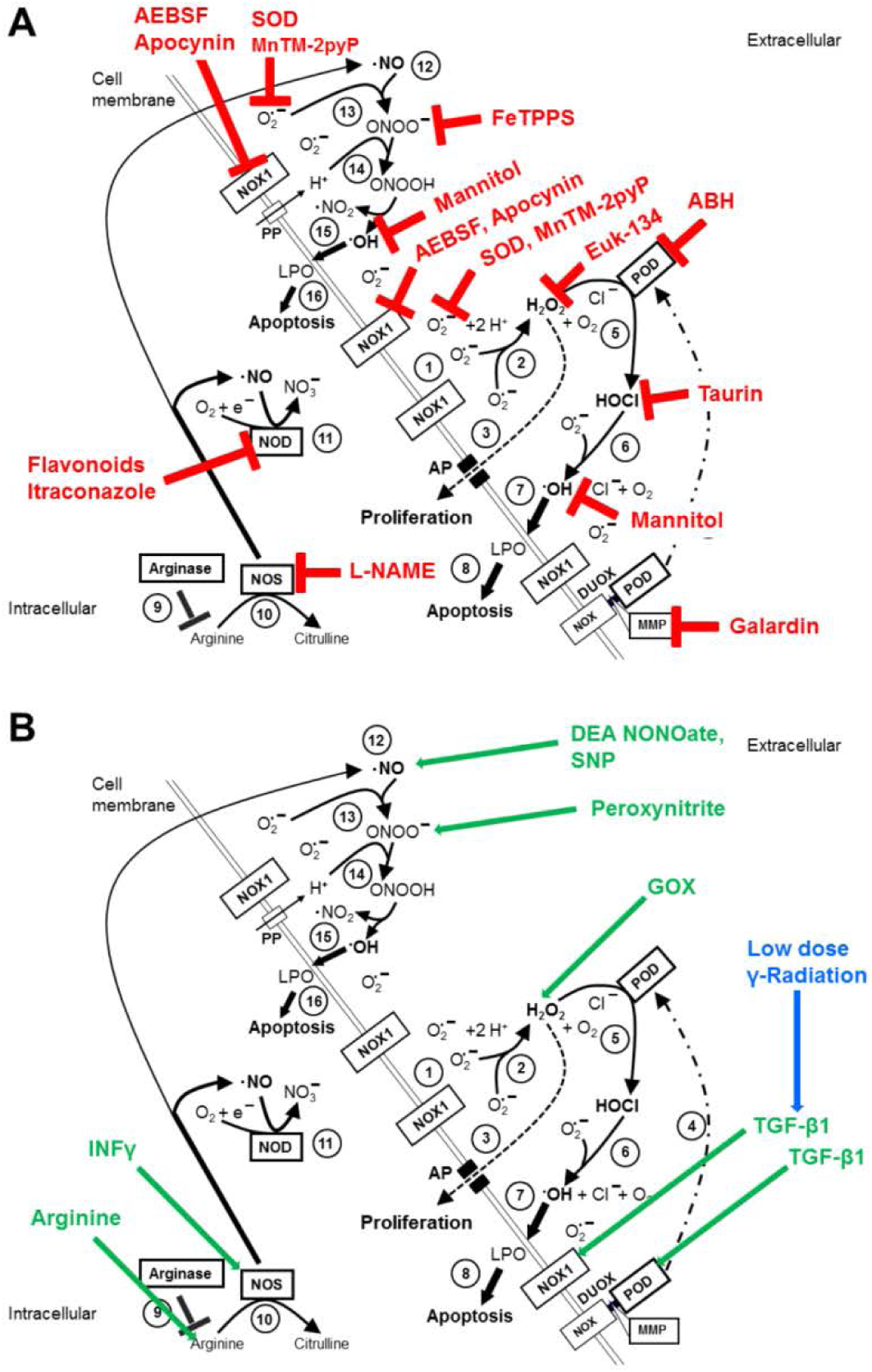
Site of action of inhibitors, scavengers and other redox-relevant compounds. This figure is based on the demonstration of the HOCl- and the NO/peroxynitrite signaling pathway as shown before in Figure 1A. HOCl- and NO/peroxynitrite signaling is active in the case of transformed cells as well as tumor cells with inactivated membrane-associated catalase (not shown in the figure). Note that these two pathways do not act simultaneously. A. Action of inhibitors/scavengers. Inhibition of NOX1 by AEBSF or apocynin prevents i) the generation of peroxynitrite, ii) the generation of H_2_O_2_ and iii) hydroxyl radical generation through HOCl/superoxide anion interaction. Scavenging of superoxide anions by SOD or the SOD mimetics MnTM-2pyP or MnTBAP (not shown in the figure) prevents i) peroxynitrite formation and ii) HOCl/superoxide anion interaction, but does not prevent the generation of H_2_O_2_ and subsequent HOCl synthesis. Mannitol scavenges hydroxyl radicals and therefore prevents lipid peroxidation/apoptosis induction by the HOCl and the NO/peroxynitrite signaling pathway. AEBSF, apocynin, SOD, SOD mimetics and mannitol therefore inhibit HOCl- as well as NO/peroxynitrite signaling. Inhibition of NO synthase (NOS) by L-NAME as well as decomposition of peroxynitrite by FeTPPS selectively inhibit NO/peroxynitrite signaling. Decomposition of H_2_O_2_ by the catalase mimetic EUK-134, inhibition of peroxidase by 4-aminobenzoyl hydrazide (ABH) or scavenging of HOCl through taurine interfere selectively with HOCl signaling. Galardin prevents the release of the DUOX-coded peroxidase domain, which leads to site-specific signaling effects [28]. Inhibition of NOD by flavonoids or itraconazole [13, 40, 87] leads to a relative increase in the concentration of NO which has modulatory effects on signaling. B. Effects of added compounds. Addition of exogenous NO by NO donors like DEA NONOate or SNP, peroxynitrite or H_2_O_2_-generating glucose oxidase (GOX) has been instrumental for our analysis. The concentration of endogenous NO can be enhanced by the addition of arginine (the substrate for NOS) or interferon gamma (an inducer of NOS expression). Low dose radiation causes an enhancement of TGF-beta activity, which is followed by increased NOX1 and DUOX1 activity [79]. This figure does not summarize the multiple ways for inhibition of membrane-associated catalase of tumor cells. Please have a look at references [6, 7, 13, 28, 30, 34, 39–41, 43, 45].

### Media for cell culture

Cells were either kept in Eagle’s Minimum Essential Medium (EMEM), containing 5% inactivated fetal bovine serum (Biochrom, Berlin, Germany) or in RPMI 1640 medium, containing 10% inactivated fetal bovine serum, as indicated for the respective cell lines. Both media were supplemented with penicillin (40 U/ml), streptomycin (50 µg/ml), neomycin (10 µg/ml), moronal (10 U/ml) and glutamine (280 µg/ml). Cells were grown in plastic tissue culture flasks and passaged once or twice weekly.

### Cells

208Fsrc3 cells, i. e. 208F rat fibroblasts transformed through constitutive expression of *v-src* were a generous gift by Drs C. Sers and R. Schäfer, Berlin, Germany. 208Fsrc3 cells have not been under the selection pressure of natural antitumour mechanisms in vivo. However, they possess the potential to induce tumors in immunocompromised animals. Transformed cells show criss cross morphology in monolayer, colony formation in soft agar and extracellular superoxide anion generation that drives both the efficiency and selectivity of intercellular ROS signaling. 208Fsrc3 cells were cultured in EMEM, 5% inactivated fetal bovine serum and supplemented as indicated above.

The human gastric carcinoma cell line MKN-45 was purchased from DSMZ, Braunschweig, Germany and was cultured in RPMI 1640, 10% inactivated bovine serum, and supplements. The human lymphoma cell line Gumbus was a gift from Dr. G. Doelken, Greifswald and was cultured in RPMI 1640, 10 % FBS. The murine fibrosarcoma cell line L929 was obtained from Dr. D. Adam, Kiel, Germany and was cultured in EMEM, 5% fetal bovine serum plus supplements. MKN-45, Gumbus and L929 cells show extracellular superoxide anion generation, but are resistant against intercellular ROS signaling through expression of membrane-associated catalase [26, 28, 30].

### Methods

#### 2.3.1 Autocrine apoptosis induction by intercellular ROS/RNS signaling

Cells in complete medium were seeded in 96-well tissue culture clusters at a density of 12 500 cells/100 µl. Modifications of the density are specified in the respective figure legends. Assays for the determination of autocrine apoptosis induction in transformed cells where supplemented with 20 ng/ml TGF-beta1 to compensate for the dilution of cell-derived TGF-beta. Assays for the determination of autocrine apoptosis in bona fide tumor cells required the inhibition of membrane-associated catalase. Further details are outlined in the respective figure legends. In all experiments, assays were performed in duplicate. After the indicated time of incubation at 37°C and 5% CO_2_ that allowed intercellular ROS-mediated apoptosis induction, the percentage of apoptotic cells was determined by inverted phase contrast microscopy based on classical criteria for apoptosis as described below.

#### 2.3.2 Determination of the percentage of apoptotic cells

After the indicated time of incubation at 37°C and 5% CO_2_, the percentage of apoptotic cells was determined by inverted phase contrast microscopy based on the classical criteria for apoptosis, i.e., nuclear condensation/fragmentation or membrane blebbing [9, 28, 57–59]. The characteristic morphological features of intact and apoptotic cells, as determined by inverted phase contrast microscopy have been published [18, 28, 40, 56, 57, 60]. At least 200 neighbouring cells from randomly selected areas were scored for the percentage of apoptotic cells at each point of measurement. Control assays ensured that the morphological features ‘nuclear condensation/fragmentation’ as determined by inverse phase contrast microscopy were correlated to intense staining with bisbenzimide and to DNA strand breaks, detectable by the TUNEL reaction [9, 40, 56, 60]. A recent systematic comparison of methods for the quantitation of apoptotic cells has shown that there is a perfect coherence between the pattern of cells with condensed/fragmented nuclei (stained with bisbenzimide) and TUNEL-positive cells in assays with substantial apoptosis induction, whereas there was no significant nuclear condensation/fragmentation in control assays [40, 57]. Further controls ensured that ROS-mediated apoptosis induction was mediated by the mitochondrial pathway of apoptosis, involving caspase-9 and caspase-3 [16, 40].

##### Determination of signaling pathways and ROS/RNS interactions

The readout in our experiments was the percentage of apoptotic cells. The role of defined ROS or RNS in apoptosis-inducing signaling or action towards this signaling was elucidated by inhibiting redox-relevant enzymes like NOX1, iNOS or nNOS, NO dioxygenase (NOD), peroxidase and catalase. In some experiments presented under Supplementary Materials, NOX1, iNOS, nNOS, NO dioxygenase peroxidase and catalase were knocked down by treatment of the cells with small interfering RNA (siRNA). In addition, the study of signaling chemistry was based on the determination whether scavenging specific compounds like superoxide anions, H_2_O_2_, HOCl, peroxynitrite, hydroxyl radicals had a significant impact on signaling-dependent apoptosis induction. Finally, our study involved the addition of defined signaling-relevant compounds like NO donors, H_2_O_2_-generating glucose oxidase, peroxynitrite, interferon (that induces NOS expression) or flavonoids that increase the endogenous NO concentration through inhibition of NOD. The site of action of inhibitors/scavengers, as well as additional compounds is summarized in Figure 3.

##### Statistical analysis

In all experiments, assays were performed in duplicate. The empirical standard deviation was calculated and is shown in the figures. Absence of standard deviation bars for certain points indicates that the standard deviation was too small to be reported by the graphic program. Empirical standard deviations were calculated merely to demonstrate how close the results were obtained in parallel assays within the same experiment and not with the intention of statistical analysis of variance, which would require larger numbers of parallel assays. The key experiments have been repeated more than three times, all others at least twice (with duplicate assays). The Yates continuity corrected chi-square test was used for the statistical determination of significances (p < 0.01 = significant; p < 0.001 = highly significant).

## Results

### NO antagonizes H_2_O_2_-dependent apoptosis-inducing HOCl signaling of transformed cells

Src oncogene-transformed fibroblasts were cultured under conditions that allowed autocrine HOCl pathway-mediated apoptosis induction selectively in transformed cells. Parallel controls with nontransformed cells showed no apoptosis induction (data not shown), in line with preceding studies [61]. The cell cultures of 208F src3 cells were confronted with increasing concentrations of the NO donor SNP (Figure 4).

**Figure 4.**
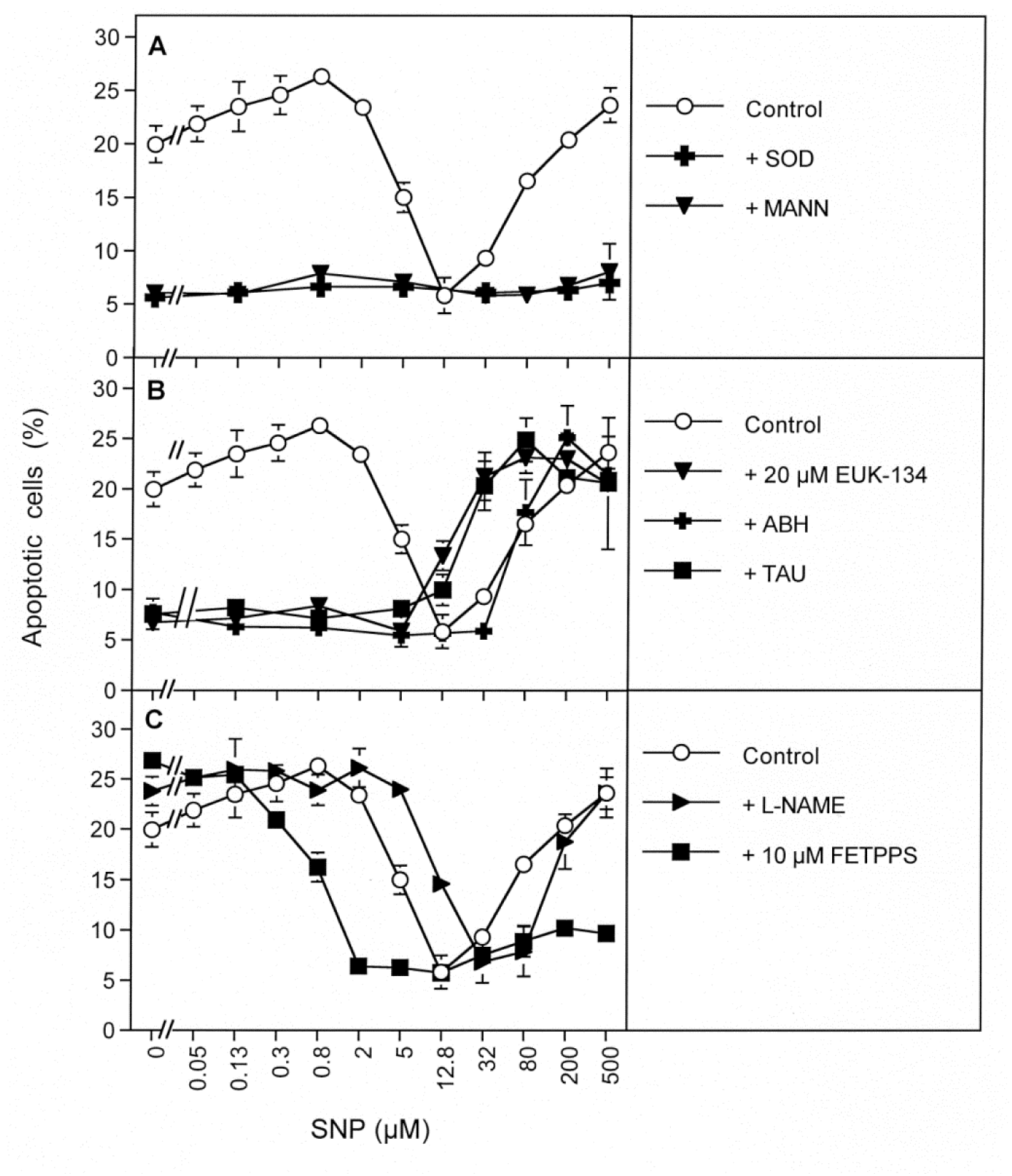
Mechanistic aspects of the antagonism between NO and H_2_O_2_-dependent HOCl signaling. 208Fsrc3 cells were seeded at a density of 12 500 cells/assay in the presence of 20 ng/ml TGF-beta. These conditions allowed autocrine apoptosis induction through HOCl signaling. Where indicated, assays received 100 U/ml MnSOD, 10 mM of the hydroxyl radical scavenger mannitol, 2 µM of the catalase mimetic EUK-134, 150 µM of the peroxidase inhibitor ABH, 50 mM of the HOCl scavenger taurine, 2.4 mM of the NOS inhibitorL-NAME, or 10 µM of the peroxynitrite decomposition catalyst FeTPPS. The NO donor SNP was added at the indicated concentrations and apoptosis induction was determined after 19 h. A. The NO donor caused an initial increase in apoptosis induction, followed by inhibition that was complete at 12.8 µM SNP. Further increase in SNP caused resumption of apoptosis induction. In all assays apoptosis was dependent on superoxide anions and mediated by hydroxyl radicals. B. Apoptosis induction in the absence of SNP and up to 12.8 µM SNP was dependent on HOCl signaling, as it required H_2_O_2_, peroxidase and taurine. EUK-134 and taurine cause a leftward shift of the right flank of the bell-shaped SNP-dependent curve. C. Apoptosis induction at concentrations higher than 12.8 µM SNP was dependent on NO-dependent peroxynitrite formation, with exogenous NO as major source, but contribution by endogenous NO as well, as seen by the inhibition curve by L-NAME. 10 µM FeTPPS also causes a strong enhancement of SNP-mediated inhibition of HOCl signaling, seen by the leftward shift of the inhibition curve. *Statistical analysis: Autocrine apoptosis induction, its inhibition by 5 and 12.8 µM SNP, as well as resumption of apoptosis induction at concentrations higher than 32 µM SNP were highly significant (p < 0.001), whereas the slight increase of apoptosis induction by 0.05-0.8 µM SNP was not significant. High significance (p < 0.001) was obtained for inhibition by SOD and mannitol on both flanks of the SNP-dependent curve, the inhibition by EUK-134, ABH and taurine on the left flank of the inhibition curve, the leftward shift of the right flank of the curve through EUK-134 and taurine, inhibition of the right flank of the SNP-determined curve by 10 µM FeTPPS and for the rightward shift of the inhibition curve by L-NAME. 10 µM FeTPPS caused a highly significant leftward shift of the left flank of the SNP-determined curve and highly significant inhibition at the right flank of this curve*.

Addition of SNP up to 0.8 µM caused a slight increase in HOCl-mediated apoptosis induction. In the presence of 2 – 12.8 µM SNP, HOCl-mediated apoptosis induction was gradually inhibited. Inhibition was dependent on the concentration of the NO donor. At 12.8 µM SNP, inhibition of apoptosis induction was complete. Further increase in the concentration of SNP caused resumption of apoptosis induction, which reached a maximal value at 500 µM SNP. The use of specific inhibitors revealed that apoptosis induction in control assays without SNP was exclusively depending on HOCl signaling, as it was completely inhibited by the HOCl scavenger taurine, the peroxidase inhibitor ABH and the catalase mimetic EUK-134 (Figure 4 B), as well as by SOD and the hydroxyl radical scavenger mannitol (Figure 4 A). In the absence of SNP, no inhibition by the NOS inhibitor L-NAME or the peroxynitrite decomposition catalyst FeTPPS was observed (Figure 4 C). In contrast, apoptosis induction in the presence of SNP concentrations higher than 12.8 µM seemed to exclusively depend on NO/peroxynitrite signaling, as there was strong inhibition by 10 µM FeTPPS (Figure 4 C), as well as by SOD and mannitol (Figure 4 A), but not by taurine or ABH (Figure 4 B). These findings confirm that SNP has released NO under the culture conditions. At lower concentrations of SNP, the released NO seemed to modulate HOCl signaling, whereas at higher concentrations it was interacting with superoxide anions to generate apoptosis-inducing peroxynitrite. This then caused apoptosis induction through generation of hydroxyl radicals. As the NOS inhibitor L-NAME only caused a distinct rightward shift of the curve, but did not block apoptosis induction at higher concentrations of SNP, SNP-derived NO rather than NO generated by NOS seemed to play the major part during inhibition of HOCl signaling and establishment of NO/peroxynitrite signaling. The FeTPPS-mediated leftward shift of the inhibition curve was due to a more complex signaling phenomenon which will be analyzed in Supplementary Figure S1. The result seems to be indicative of an involvement of singlet oxygen generation, catalase inactivation and subsequent increase in free H_2_O_2_ (Supplementary Figure S1). In summary, the experiment shown in Figure 4 can be dissected into three subsequent NO-controlled steps: i) initial increasing of HOCl signaling by SNP up to 0.8 µM SNP, ii) complete inhibition of HOCl signaling by 0.8-12.8 µM SNP, and iii) establishment of NO/peroxynitrite signaling at higher SNP concentrations (12.8-500 µM SNP). Importantly, in this experiment, NO/peroxynitrite-dependent apoptosis signaling was not established in parallel to HOCl signaling. *Rather, it required inhibition of HOCl signaling by NO before NO/peroxynitrite signaling became prominent at higher concentrations of the NO donor.* Furthermore, the catalase mimetic EUK-134 as well as the HOCl scavenger taurine, caused a left-ward shift of the curve that described the dependence of NO/peroxynitrite signaling on the concentration of exogenous SNP (Figure 4 B). These findings are indicative of consuming effects of H_2_O_2_ as well as HOCl directed towards NO/peroxynitrite signaling. These consuming effects seem to be abrogated by the catalase mimetic EUK-134 and the HOCl scavenger taurine. Therefore, under conditions of removal of either H_2_O_2_ or HOCl, less NO is required for establishment of NO/PON signaling as in the presence of a parallel consuming effect and the concentration-response curve is shifted leftward. These consuming reactions are one of the clues for the understanding of NO/H_2_O_2_ interactions.

### NO has the potential to either antagonize or to restore H_2_O_2_-dependent HOCl signalling of tumor cells, dependent on the concentration of H_2_O_2_

Increasing concentrations of the catalase inhibitor 3-AT abrogated the catalase-mediated protection of the human lymphoma cell line Gumbus towards intercellular ROS-mediated signaling. The presence of 3-AT allowed apoptosis induction in the mode of an optimum curve (Figure 5 A). In this cell line, apoptosis induction at suboptimal and optimal concentrations of 3-AT (25 and 75 mM) seemed to be due to signaling through the HOCl signaling pathway, as it was strongly inhibited by the NOX1 inhibitor AEBSF, the superoxide anion scavenger SOD, the peroxidase inhibitor ABH, the HOCl scavenger taurine and the hydroxl radical scavenger mannitol (Figure 5 B, C). Addition of the catalase mimetic EUK-134 to assays containing 50 or 75 mM 3-AT caused inhibition of apoptosis induction dependent on the concentration of EUK-134 (Figure 5 D). When EUK-134 was added to assays containing a supraoptimal concentration of 3-AT (200 mM), the relatively low apoptosis induction was initially enhanced with increasing low concentrations of EUK-134. Further increase in the concentration of the catalase mimetic then caused concentration-dependent inhibition of apoptosis. These findings show that H_2_O_2_ is establishing HOCl signaling after appropriate inhibition of catalase. However, when high concentrations of the catalase inhibitor were leading to an apparent relative excess of H_2_O_2_, H_2_O_2_ seemed to inhibit HOCl signaling. This is explained by the reaction between excess H_2_O_2_ with HOCl [27], following the reaction described by Held et al. and Connick [53, 54]:

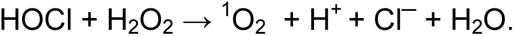

**Figure 5.**
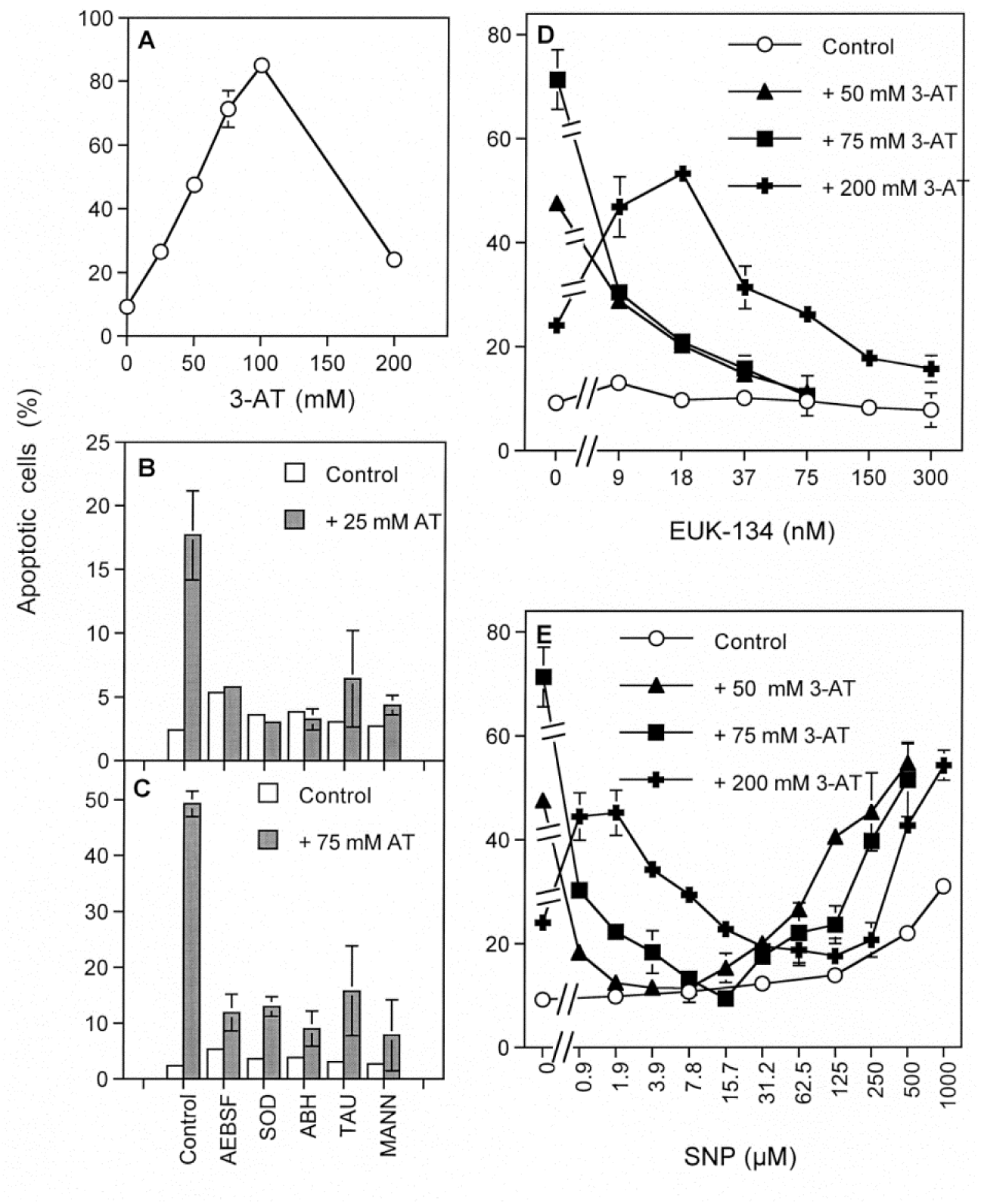
The catalase mimetic EUK-134 and the NO donor SNP antagonize H_2_O_2_-dependent HOCl signaling of the human lymphoma cell line Gumbus. In all assays, Gumbus cells were applied at a density of 25 000 cells/100 µl. A. 3-AT-mediated apoptosis induction in Gumbus cells at 5 h showed the mode of an optimum curve. B, C: Apoptosis induction by 25 mM and 75 mM 3-AT at 4 h was inhibited by 100 µM of the NOX inhibitor AEBSF, 100 U/ml MnSOD, 150 µM of the mechanism-based peroxidase inhibitor ABH, 50 mM of the HOCl scavenger taurine and 10 mM of the hydroxyl radical scavenger mannitol. Untreated cells were protected towards apoptosis induction (“control”). D: Increasing concentrations of the catalase mimetic EUK-134 inhibited apoptosis induction by 50 and 75 mM 3-AT in a concentration-dependent mode. Supraoptimal apoptosis induction at 200 mM 3-AT was enhanced by low concentrations of EUK-134, followed by inhibition at higher concentrations of EUK-134. E: The NO donor SNP inhibited apoptosis induction mediated by 50 and 75 mM 3-AT. Supraoptimal apoptosis induction at 200 mM 3-AT was enhanced by low concentrations of SNP, followed by inhibition at higher concentrations of SNP. After maximal inhibition of 3-AT-mediated apoptosis SNP induced apoptosis. Apoptosis induction was dependent on the concentrations of SNP and 3-AT. *Statistical analysis: Induction of apoptosis mediated by 3-AT (A), inhibition of 3-AT-mediated apoptosis by AEBSF, SOD, ABH, taurine and mannitol, inhibition of apoptosis induced by 50 and 75 mM 3-AT through EUK-134 as well as SNP, increase of apoptosis induction by 200 mM 3-AT through EUK-134 and SNP were highly significant (p < 0.001)*.

Removal of the inhibitory excess of H_2_O_2_ through well-balanced concentrations of the catalase mimetic EUK-134 restored optimal HOCl signaling. Further increase in the concentration of the catalase mimetic then abrogated HOCl signaling, conceivably through lowering the concentration of H_2_O_2_ below the level that is needed for HOCl synthesis.

When increasing concentrations of the NO donor sodium nitroprusside (SNP) were added rather than the catalase mimetic (Figure 5 E), an analogous effect to that of the catalase mimetic was initially observed. SNP inhibited apoptosis induction mediated by 50 and 75 mM 3-AT. At higher concentrations of SNP (in the presence of 50 and 75 mM 3-AT), SNP-mediated apoptosis induction resumed. In the presence of 200 mM 3-AT, i. e. at 3-AT concentrations that favoured H_2_O_2_-dependent inhibition of HOCl signaling, lower concentrations of SNP initially enhanced apoptosis induction. Further increase of the SNP concentration then inhibited apoptosis induction. This was followed by SNP-mediated apoptosis at higher SNP concentrations. These findings demonstrate strong and concentration-dependent interactions between NO and H_2_O_2_-dependent signaling processes, resulting either in inhibition or stimulation of HOCl signaling. These reactions were analogous to those of the catalase mimetic EUK-134 and therefore might be explained by controlled decomposition or removal of H_2_O_2_. At high concentrations of SNP, these reactions were different from those of EUK-134, as NO/peroxynitrite signaling resumed in the presence of high concentrations of SNP, whereas EUK-134, as expected, had no own signaling potential.

The effects of NO towards HOCl signaling as shown in Figure 5 were fully confirmed for the murine fibrosarcoma cell line L 929 (Supplementary Figure S2).

In line with a previous report [28], inhibition of membrane-associated catalase of the human gastric carcinoma cell line MKN-45 caused a more complex signaling pattern than that of the cell lines Gumbus and L929. Figure 6 A shows that 3 mM and 6 mM 3-AT caused apoptosis induction of MKN-45 cells exclusively through the NO/peroxynitrite signaling pathway, as demonstrated by the inhibition through L-NAME and FeTPPS, and the lack of inhibition by taurine. In contrast, higher concentrations of 3-AT caused apoptosis signaling through the HOCl pathway, as demonstrated by taurine-mediated inhibition, but not by L-NAME- or FeTPPS-dependent inhibition. HOCl signaling reached its optimum at 75 mM 3-AT and then turned into supraoptimal inhibition at higher concentrations of the catalase inhibitor. The latter has been shown to be caused by the interfering effect of excess H_2_O_2_ on HOCl signaling [27], based on the direct reaction between H_2_O_2_ and HOCl [27, 28, 35]. At low concentrations of 3-AT, addition of 50 µM of the NO donor SNP or 0.75 mM of the NOS substrate arginine caused a distinct enhancement of NO/peroxynitrite signaling. This was followed by nearly complete inhibition of HOCl signaling by SNP or arginine, at 3-AT concentrations up to 75 mM 3-AT (Figure 6B). At higher concentrations of 3-AT, SNP as well as arginine counteracted the supraoptimal decline of apoptosis induction. Thereby, HOCl signaling was maintained and NO/peroxynitrite signaling was not established.

**Figure 6.**
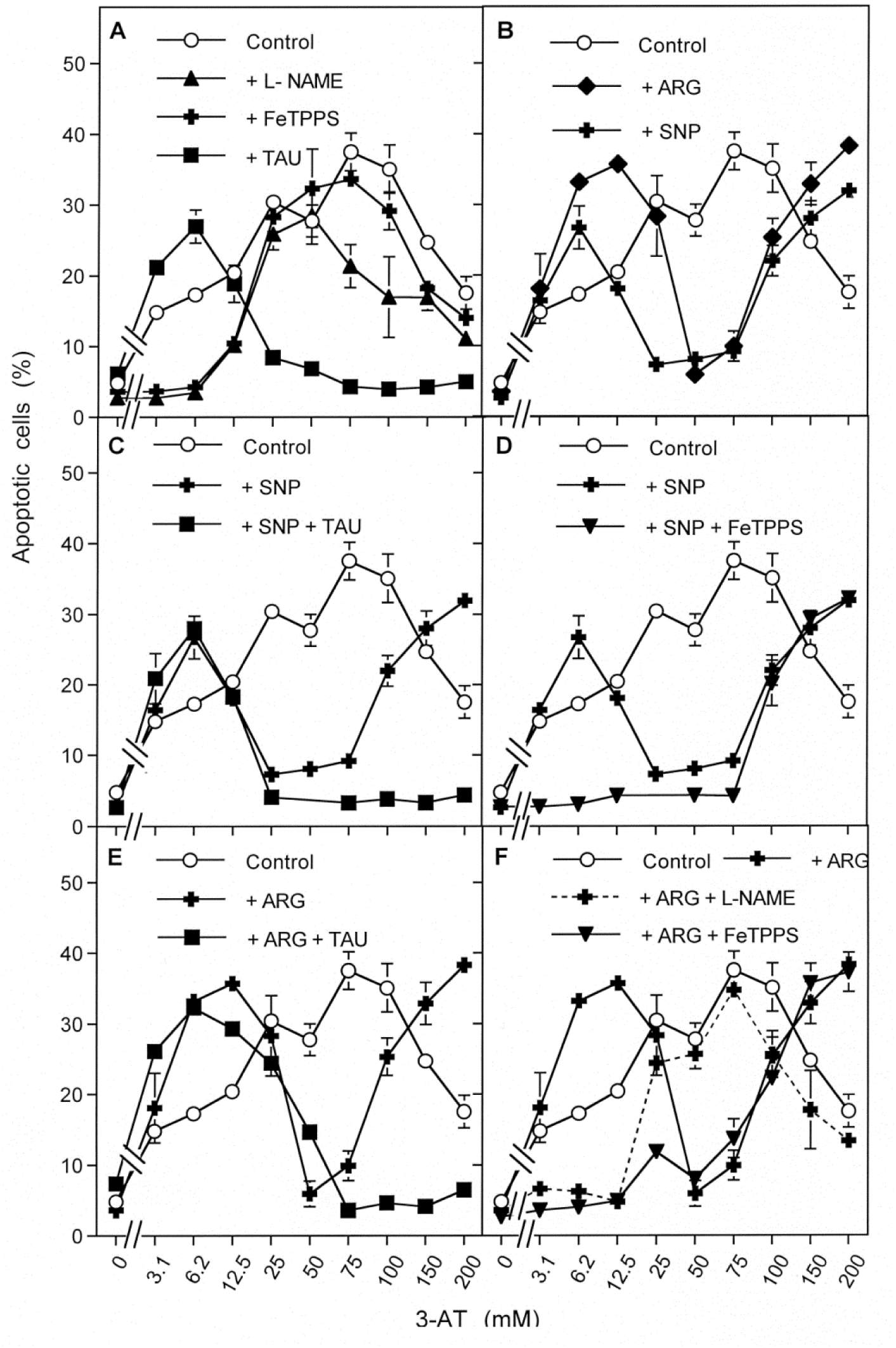
Interaction between exogenous and endogenous NO with NO/peroxynitrite and HOCl signaling after catalase inhibition in the human gastric carcinoma cell line MKN-45. A. MKN-45 cells were treated for 5 hs with increasing concentrations of 3-AT in control cells (no further addition) and in the presence of 2.4 mM of the NOS inhibitor L-NAME, 25 µM of the peroxynitrite decomposition catalyst FeTPPS or 50 mM of the HOCl scavenger taurine. B. As indicated, cells received 50 µM of the NO donor SNP, 50 µM SNP plus 25 µM FeTPPS or 50 µM SNP plus 50 mM taurine and were cultivated in the presence of increasing concentrations of 3-AT. C,D: As indicated, cells received 0.75 mM of the NOS substrate arginine, 0.75 mM arginine plus 50 mM taurine, 0.75 mM arginine plus 2.4 mM L-NAME or 0.75 mM arginine plus 25 µM FeTPPS and were cultivated in the presence of increasing concentrations of 3-AT. The result demonstrates the differential induction of NO/peroxynitrite and HOCl signaling with increasing concentrations of the catalase inhibitor. Exogenous as well as endogenous NO enhances NO/peroxynitrite signaling, inhibits suboptimal and optimal HOCl signaling and brings supraoptimal HOCl signaling back to its optimum. *Statistical analysis: Apoptosis induction by 3-AT was highly significant (p < 0.001). The inhibitory effect of L-NAME and FeTPPS at 6.2 mM 3-AT (*Figure 6A*), the stimulatory effect of taurine at 6.2 mM 3-AT and the inhibitory effect of taurine at 3-AT concentrations higher than 25 mM were highly significant (p < 0.001). SNP and arginine dependent i) enhancement of NO/peroxynitrite-mediated apoptosis induction, ii) inhibition of HOCl-mediated apoptosis induction (25-100 mM 3-AT) and abrogation of supraoptimal inhibition of HOCl signaling (200 mM 3-AT) were highly significant (p < 0.001). The FeTPPS-dependent inhibition of NO/peroxynitrite signaling in the presence of SNP and arginine was highly significant (p < 0.001), whereas there was no significant effect of FeTPPS on NO-mediated effects directed towards HOCl signaling. In contrast, the arginine-dependent NO-mediated effects towards HOCl signaling were inhibited by L-NAME with high significance (p < 0.001)*. product peroxynitrite, as FeTPPS did not counteract this inhibition.

The control of the SNP-mediated effects, using the HOCl scavenger taurine and the peroxynitrite decomposition catalyst FeTPPS (Figure 6 C, D) demonstrated that the SNP-mediated effect at low concentrations of 3-AT was due to enhancement of NO/peroxynitrite signaling, partially also on the expense of HOCl signaling. This was then followed by SNP-mediated inhibition of HOCl signaling between 25 and 75 mM 3-AT. This inhibition seemed to be directly mediated by NO, and not by its reaction The reactivation of apoptosis-inducing signaling as seen at 3-AT concentrations higher than 75 mM was characterized as NO-dependent reactivation of HOCl signaling. This particular NO-mediated step was independent of the action of peroxynitrite.

The inhibitor-based analysis of the effects of arginine (Figure 6 E, F) showed that the NOS substrate arginine caused an effect analogous to that of SNP, i. e. it enhanced NO/peroxynitrite signaling at low concentrations of 3-AT, inhibited HOCl signaling at intermediate concentrations of 3-AT and counter-acted supraoptimal inhibition of HOCl signaling at high concentrations of 3-AT. Inhibition of HOCl signaling as well as restoration of HOCl signaling at supraoptimal concentrations of 3-AT were dependent on NOS-derived NO, as seen by the abrogatory effect of the NOS inhibitor L-NAME on these effects. These NO-mediated effects were independent of the formation of peroxynitrite, as seen by the lack of effect of the peroxynitrite decomposition catalyst FeTPPS. In contrast, the stimulatory effect of arginine at low concentrations of 3-AT was inhibited by L-NAME and FeTPPS, indicating that this was due to NO-dependent, peroxynitrite-mediated apoptosis-inducing signaling. This analysis therefore allows to differentiate between the effects of NO on NO/peroxnitrite-dependent apoptosis-inducing signaling and two NO-mediated, peroxynitrite-independent effects on H_2_O_2_-dependent processes related to HOCl signaling. These discrete NO-dependent effects on H_2_O_2_/HOCl signaling are adverse, as NO interferes with optimal H_2_O_2_-dependent HOCl signaling, but also restores HOCl signaling that is counteracted by excess H_2_O_2_ due to strong inhibition of catalase. The inhibitory effect of NO on HOCl signaling can be explained both by NO- dependent decrease in H_2_O_2_ or HOCl, or both. In contrast, the NO-dependent restoration of HOCl signaling at very high H_2_O_2_ concentration can only be explained by NO-dependent decrease in the concentration of H_2_O_2_.

The enhancing effect of SNP on NO/peroxynitrite signaling and its inhibitory as well as enhancing effect on HOCl signaling were dependent on the concentration of the NO donor, as shown in Supplementary Figure S3.

NO dioxygenase (NOD), converts NO into nitrate (2 .NO + 2 O_2_ + NADPH → 2 NO ^−^ + NADP^+^ + H^+^) and thus leads to constant intracellular consumption of NO [62–65]. Its inhibition has been shown to represent an effective way for the modulation of the endogenous and extracellular NO concentration of tumor cells [11, 13]. Quercetin as well as azoles have been reported to inhibit NOD [63]. In addition, an inhibitory potential of anthocyanidins for NOD has been shown [40]. Addition of NOD inhibitors to malignant cells should therefore have an analogous effect on HOCl signaling as addition of an NO donor or of the NOS substrate arginine. As shown in Figure 7, increasing concentrations of the anthocyanidins delphinidin, peonidin, pelargonin, the flavonoids quercetin and xanthohumol, or the azole itraconazole caused concentration-dependent inhibition of HOCl-mediated apoptosis induction in 208Fsrc3 cells. At higher concentrations of the compounds, apoptosis induction resumed. Controls containing the NOS inhibitor L-NAME confirmed that inhibition of HOCl-mediated apoptosis signaling, as well as resumption of apoptosis induction, were dependent on NOS-mediated NO synthesis. As shown for quercetin, this modulatory potential was equally seen for the aglycon quercetin and for distinct glycosylated quercetin derivatives like quercitrin, isoquercitrin and rutin (Supplementary Figure S4).

**Figure 7.**
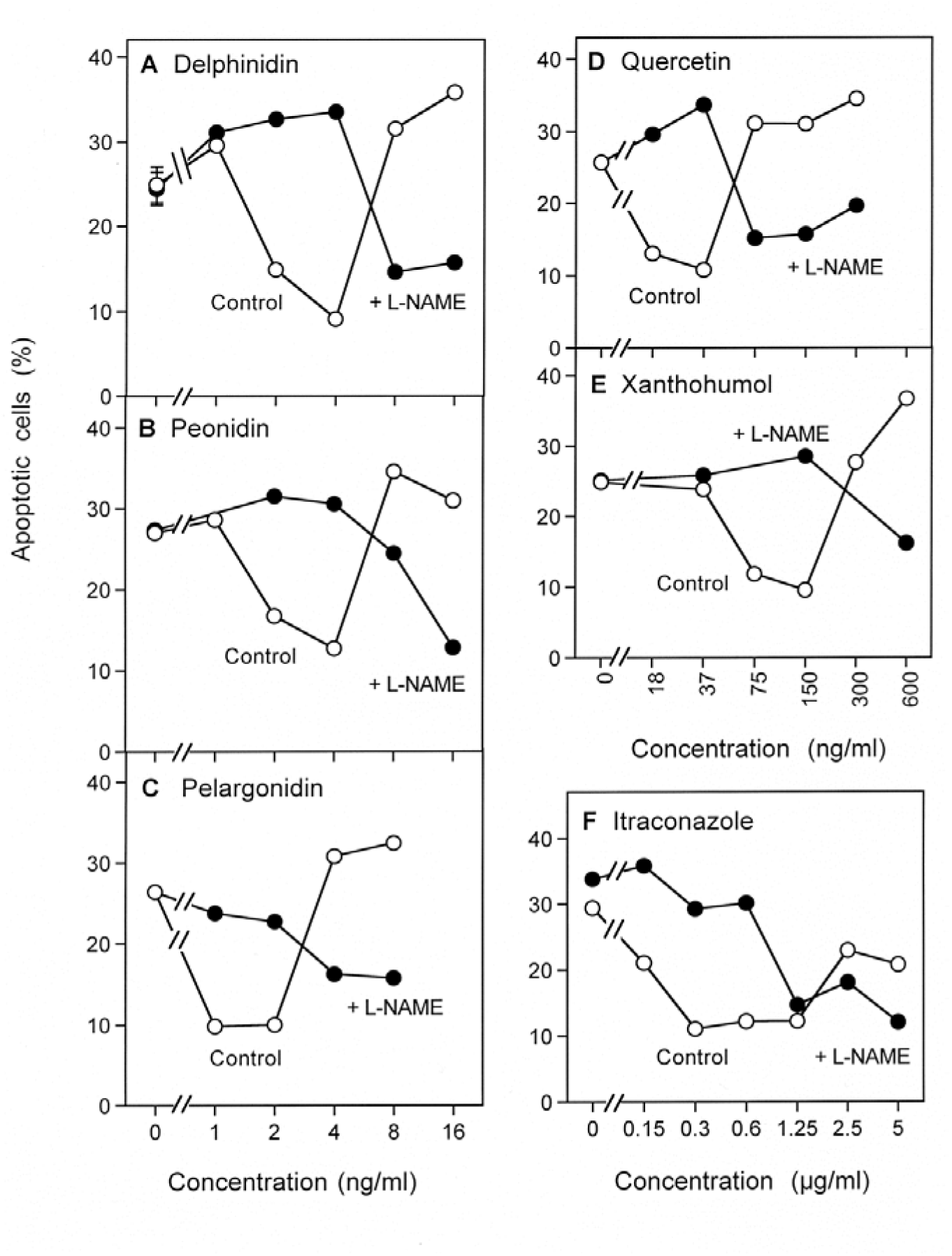
The effect of increasing cell-derived NO concentrations through application of NOD inhibitors. Transformed 208 Fsrc3 cells were seeded at a density of 12 500 cells/assay in the presence of 20 ng/ml TGF-beta1. The assays received either 2.4 mM of the NOS inhibitor L-NAME or remained free of inhibitor (“control”). The indicated concentrations of the anthocyanidins delphinidin, peonidin, pelargonin, the flavonoids quercetin and xanthohumol, or itraconazole were added. All these compounds have been defined as inhibitors of NO dioxygenase [13, 40, 87]. The percentages of apoptotic cells were determined after 16 h. All compounds used cause an initial inhibition of apoptosis induction, followed by an increase. Both effects are counteracted by the NOS inhibitor L-NAME, indicating that an increased NO concentration after inhibition of NOD through the compounds was responsible of the observed effects. *Statistical analysis: The effects of all compounds, as well as the interfering effect of L-NAME, are highly significant (p<0.001)*.

The concentration of cell-derived NO can also be efficiently modulated by enhancement of NOS expression (see reference [11] for review). Interferon gamma is known as an efficient inducer of NOS [66]. Therefore, the impact of interferon gamma treatment on autocrine ROS-driven apoptosis induction in rat fibroblasts transformed with the src oncogene was studied. 208 F src3 cells showed autocrine apoptosis induction that was mediated by the HOCl signaling pathway, i. e. it was dependent on HOCl synthesis, followed by HOCl / superoxide anion reaction and subsequent generation of apoptosis-inducing hydroxyl radicals [57]. As shown in Figure 8 A, the kinetics of autocrine apoptosis induction in 208 F src3 cells started after a lag phase of 15 hrs and was dependent on HOCl signalling. NO seemed to play no role for apoptosis induction, as the NOS inhibitor L-NAME had no inhibitory effect. In the presence of interferon gamma, the kinetics of apoptosis induction was accelerated and the signaling profile was shifted from HOCl signaling completely to NO/peroxynitrite signaling, as it was inhibited by L-NAME and SOD, whereas taurine caused no inhibitory effect (Figure 8 B). These findings demonstrate that interferon treatment had caused the generation of NO in a concentration range that allowed complete antagonism to HOCl signaling and establishment of alternative NO/peroxynitrite signaling.

**Figure 8.**
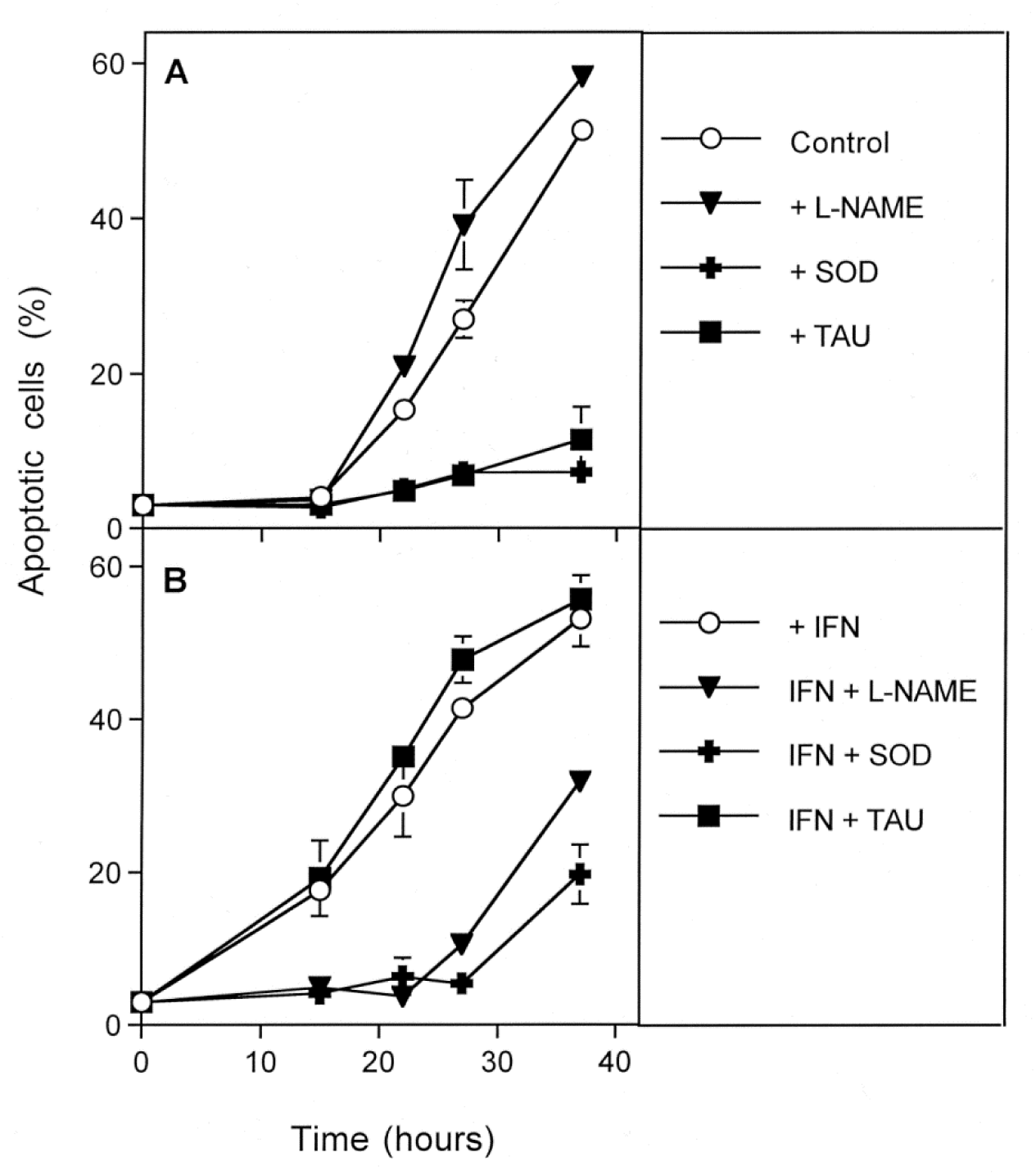
The effect of increasing cell-derived NO concentrations after application of NOS-inducing interferon gamma. Kinetic analysis. 12 500 208 Fsrc3 cells were seeded per assay and received 20 ng/ml TGF-beta1. The assays received either no addition or the indicated compounds (L-NAME 2.4 mM; Mn-SOD 100 U/ml; Taurine 50 mM, interferon gamma (IFN) 30 U/ml. The percentages of apoptotic cells were determined kinetically. A. Apoptosis induction in the absence of interferon was characterized by a lag phase and subsequent action of the HOCl signaling pathway, without contribution of NO/peroxynitrite signaling, based on the inhibition profile. B. In the presence of interferon gamma, the lag phase was shorter and apoptosis induction was completely dependent on NO/peroxynitrite signaling, without contribution by HOCl signaling. This shift in signaling quality can be explained by the NOS-inducing potential of interferon. Statistical analysis: The inhibitory action of taurine and SOD under A, and the inhibitory effect of L-NAME and SOD under B are highly significant (p<0.001).

The NO enhancing effect of interferon treatment was dependent on the concentration of interferon (Figure 9). At 15 hrs of culture, untreated 208 Fsrc3 cells were still in the lag phase before apoptosis induction. In the presence of interferon, apoptosis was enhanced and was dependent on NO/peroxynitrite signaling exclusively, as seen by the strong inhibiton by SOD and L-NAME and the lack of inhibition by taurine (Figure 9 A). Taurine even caused further enhancement of apoptosis induction in the medium concentration range of interferon. Addition of catalase caused a further enhancement of the interferon-dependent NO/peroxynitrite-dependent apoptosis induction (Figure 9 C). When apoptosis induction was analysed at 27 hours of culture, untreated 208F src3 cells had undergone apoptosis induction up to nearly 30 percent, dependent on HOCl signaling. L-NAME even caused enhancement of HOCl signaling. Increasing concentrations of interferon initially inhibited HOCl signaling and caused resumption of NO/peroxynitrite signaling at higher concentrations of interferon (Figure 9 B). Addition of catalase interfered with HOCl signaling and enhanced NO/peroxynitrite signaling in the presence of interferon (Figure 9D).

**Figure 9.**
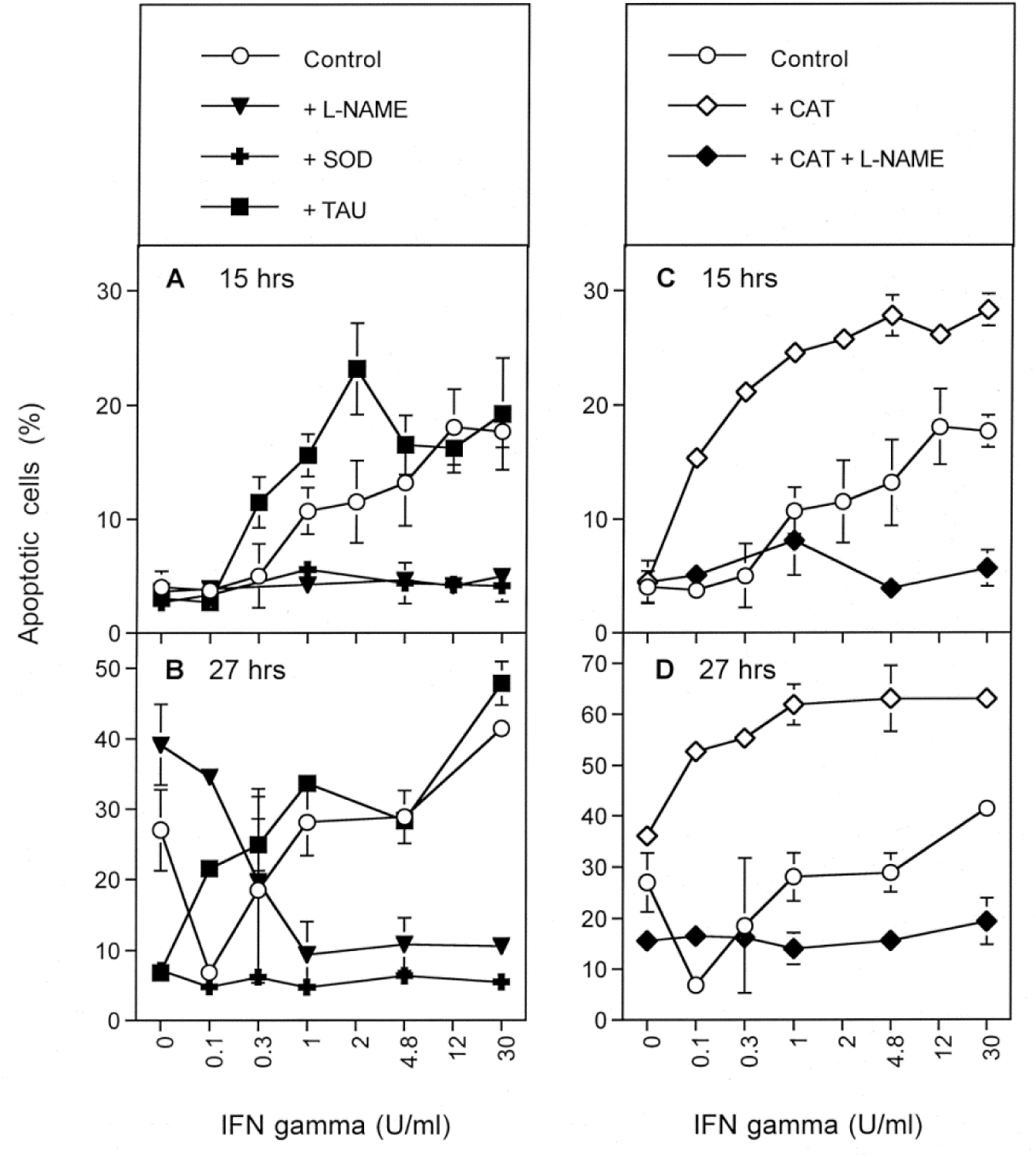
The effect of increasing cell-derived NO concentrations after application of NOS-inducing interferon gamma. Dependency on the concentration of interferon gamma. 12 500 208 Fsrc3 cells were seeded per assay and received 20 ng/ml TGF-beta1. The assays received either no addition (“Control”) or the indicated compounds (L-NAME 2.4 mM; Mn-SOD 100 U/ml; Taurine 50 mM; catalase (CAT) 100 U/ml or a combination of CAT and L-NAME. Interferon gamma was added at the indicated concentrations and the percentages of apoptotic cells were determined after 15 and 27 h. Please find further details and results in the text. *Statistical analysis: the concentration-dependent effects of interferon and the interference of L-NAME with these effects are highly significant*.

### H_2_O_2_ antagonizes NO/peroxynitrite signaling

If the concentration-dependent effects of exogenous and cell-derived NO towards H_2_O_2_-dependent HOCl signaling (Figures 4-9) was dependent on an antagonistic action between NO and H_2_O_2_, the reverse setting, i. e. the addition of H_2_O_2_ to a NO-driven process should lead to an antagonistic effect as well.

In order to analyze the modulation of NO-dependent, peroxynitrite-mediated apoptosis induction by H_2_O_2_, we induced apoptosis in transformed 208Fsc3 cells through application of the fast decaying NO donor DEA NONOate and studied the effects of H_2_O_2_ on this process. As shown in Figure 10 A, apoptosis induction by DEA NONOate was dependent on the concentration of the NO donor. It was inhibited by SOD and by the peroxynitrite decomposition catalyst FeTPPS. It therefore seems to depend on the reaction between transformed cell-derived superoxide anions and NO, resulting in the formation of apoptosis-mediating peroxynitrite. When the cell-derived H_2_O_2_ concentration was increased through doubling the cell density, apoptosis induction at the lower concentrations of the NO donor was strongly inhibited, whereas there was no inhibitory effect when high concentrations of the NO donor had been applied (Figure 10 B). The effect of the increased cell density at low concentrations of the NO donor was mimicked by the H_2_O_2_-generating enzyme GOX (Figure 10 C). The inhibitory effect of GOX was completely abrogated by 2 µM of the catalase mimetic EUK-134, indicating that it was due to H_2_O_2_. EUK-134 added to the cells in the absence of GOX also caused a distinct enhancement of apoptosis induction mediated by lower concentrations of the NO donor, pointing to an inhibitory effect on NO-mediated apoptosis induction by cell-derived H_2_O_2._

**Figure 10.**
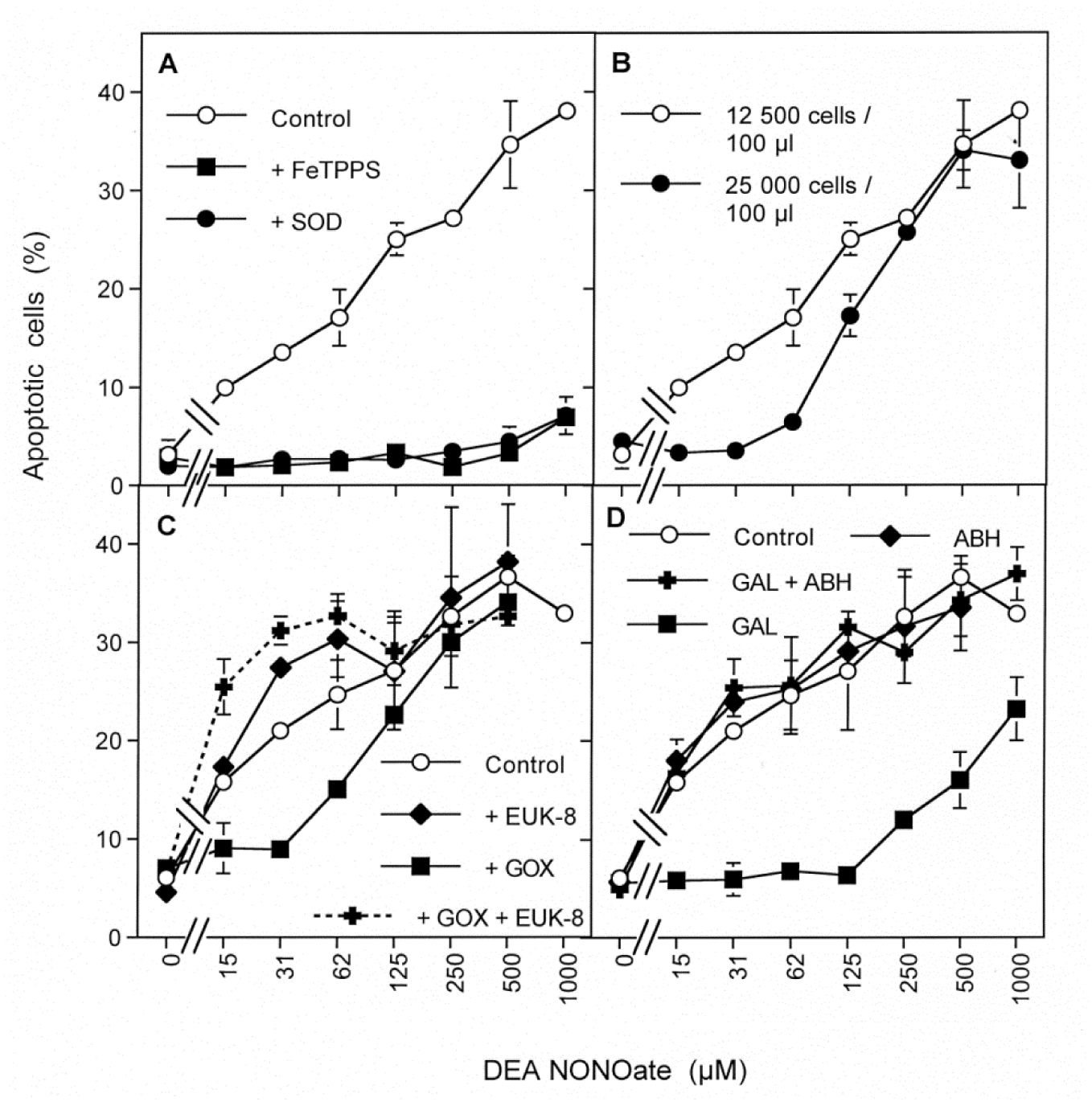
H_2_O_2_ antagonizes the NO/peroxynitrite signaling pathway. 12 500 src oncogene-transformed murine 208Fsrc3 cells per assay were treated with 20 ng/ml TGF-beta1 for 1 hr and then received the indicated additions and increasing concentrations of the NO donor DEA NONOate. Control cells were only treated with TGF-beta and DEA NONOate. Apoptosis induction was determined 3 h after addition of DEA NONOate. A. The strong inhibitory effect of 25 µM FeTPPS and 100 U/ml MnSOD on DEA NONOate-mediated apoptosis induction demonstrates that peroxnitrite generation from NO/superoxide anion interaction was the basis for apoptosis induction. B. Doubling of the cell density caused a strong inhibition of NO/peroxynitrite signaling in the low concentration range of the NO donor. C. Addition of 2 mU/ml of the H_2_O_2_-generating enzyme glucose oxidase (GOX) caused strong inhibition of apoptosis induction in the low concentration range of the NO donor. GOX-mediated inhibition was abrogated by 2 µM of the catalase mimetic EUK-8. Application of 2 µM EUK-8 alone caused a slight increase of apoptosis induction in the lower concentration range of DEA NONOate. D. Treatment of the cells with 10 µM of the matrix metalloprotein inhibitor galardin 15 minutes before the addition of the NO donor caused a strong inhibition of DEA NONOate-dependent apoptosis induction. This inhibition was abrogated by 150 µM of the peroxidase inhibitor ABH. ABH alone had no effect. *Statistical analysis: High significance (p < 0.001) was seen for i) DEA NONOate-dependent apoptosis induction and its inhibition by SOD and FeTPPS (Figure 10A), ii) the inhibitory effect of the higher cell density in the concentration range between 15 and 62 µM DEA NONOate (Figure 10B), iii) the inhibitory effect of GOX on DEA NONOate-dependent apoptosis induction in the concentration range between 15 and 62 µM DEA NONOate and the abrogation of this inhibitory effect by EUK-8 (Figure 10C), iv) inhibition of DEA NONOate-dependent apoptosis induction in galardin-pretreated cells as well as abrogation of galardin-dependent inhibition through ABH (Figure 10D). There was no significant effect of ABH on DEA NONOate-mediated apoptosis induction*.

Inhibition of the release of peroxidase from the cells by the matrix metalloprotein inhibitor galardin [67] allows to localize POD at high local concentration to the cell membrane [61]. Under these conditions, NO-mediated apoptosis induction was completed inhibited in the concentration range up to 125 µM DEA NONOate (Figure 10 D). At higher concentrations of the NO donor, inhibition became gradually weaker. The inhibitory effect in the presence of galardin was completely abrogated by the peroxidase inhibitor ABH, indicating that it was based on an enzymatic reaction of peroxidase. ABH, in the absence of galardin, caused no detectable effect on NO/peroxynitrite signaling (Figure 10 D). This result confirms that a high local concentration of POD on the membrane of malignant cells has the potential to interfere with NO/peroxynitrite-mediated apoptosis induction. However, cell-derived peroxidase has no negative effect on NO/peroxynitrite signaling under the conditions of our test system, as long as it release from the cells is not prevented.

For a further quantitative analysis of the inhibitory role of cell-derived H_2_O_2_ for NO/peroxynitrite-mediated apoptosis induction, transformed cells where treated with increasing concentrations of DEA NONOate and cell-derived extracellular H_2_O_2_ was gradually removed by increasing concentrations of soluble catalase (Figure 11 A). Catalase concentrations up to 222 U/ml enhanced NO-mediated apoptosis induction preferentially in the low concentration range of DEA NONOate. When higher concentrations of catalase were used, overall apoptosis induction was gradually inhibited. 2000 U/ml of catalase caused a nearly complete block of NO-mediated apoptosis induction. When the results derived from this experiment were plotted in relationship to the catalase concentration (Figure 11 B), the stimulatory effect of low catalase concentrations on apoptosis induction by low concentrations of DEA NONOate was clearly visualized. In the presence of 1 mM DEA NONOate, no more stimulation of apoptosis by catalase was seen. For all concentrations of DEA NONOate, a strong inhibition of apoptosis induction beyond 222 U/ml catalase was overt. In contrast, much lower concentrations of soluble catalase were needed to inhibit apoptosis induction by direct addition of peroxynitrite.

**Figure 11.**
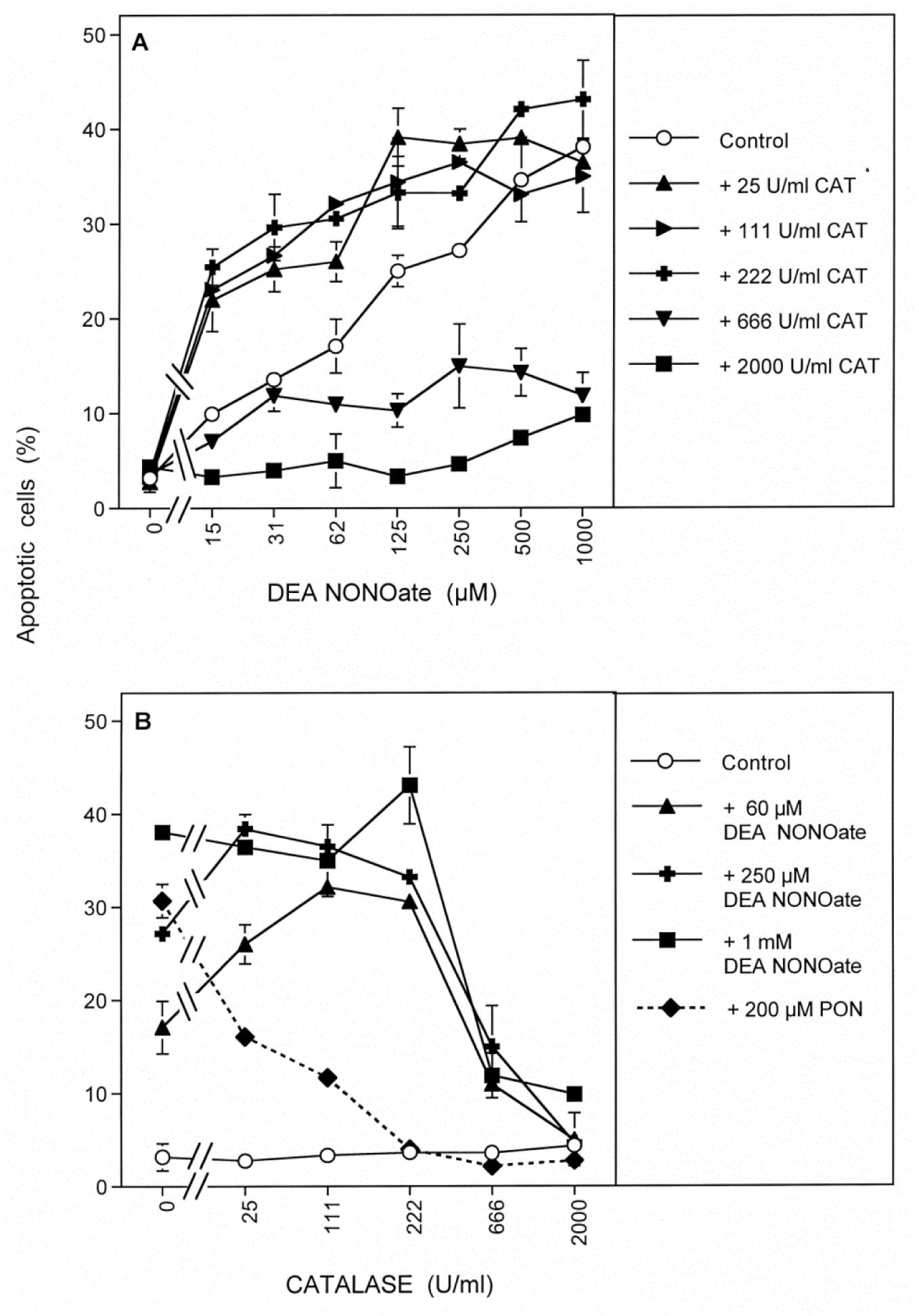
Decomposition of cell-derived H_2_O_2_ by catalase enhances NO/peroxynitrite signaling. 12 500 src oncogene-transformed murine 208Fsrc3 cells per assay were treated with 20 ng/ml TGF-beta1 for 1 hr and then received the indicated additions and increasing concentrations of the NO donor DEA NONOate, in the absence or in the presence of the indicated concentrations of catalase. The percentage of apoptotic cells was determined 3 h after addition of DEA NONOate. The results are plotted in dependence on the concentration of DEA NONate (A) and the dependence on the concentration of catalase (B). In addition, the effect of catalase on 200 µM exogenously added peroxynitrite was determined (B). The data show that catalase up to a concentration of 222 U/ml enhances NO-mediated apoptosis induction in the concentration range of the NO donor up to 0.25 mM. Higher concentrations of catalase inhibit NO/peroxynitrite-dependent apoptosis induction. Protection against apoptosis induction by exogenous peroxynitrite requires much less catalase than protection against peroxynitrite generated close to the cell membrane through NO/superoxide anion interaction. *Statistical analysis: The stimulatory effect of 25-222 U/ml catalase on apoptosis induction mediated by 0.015-0.125 mM DEA NONOate, the inhibition apoptosis induction mediated by 0.125-1 mM DEA NONOate through 666 u/ml catalase and the inhibition of DEA NONOate-mediated apoptosis induction at all concentrations of the NO donor through 2000 U/ml catalase were highly significant (p < 0.001)*.

These data also illuminate the dual mode of interaction between catalase and NO/peroxynitrite signaling. Low concentrations of soluble catalase have the potential to abrogate the negative effects of H_2_O_2_ on NO/peroxynitrite signaling, leading to an enhancement of apoptosis signaling. In contrast, very high concentrations of soluble catalase protect the cells against peroxynitrite that is formed in close vicinity of the cell membrane of the superoxide anion-generating cells.

Superoxide anions are required for peroxynitrite generation through their reaction with NO. Superoxide anions are, however, also the source for H_2_O_2_ that is generated through their dismutation and that seems to counteract NO-dependent processes. This dual role of superoxide anions and their determining function for the outcome of the experiments are analysed in Supplementary Figures S5 and S6.

These findings demonstrate that H_2_O_2_ causes a strong inhibitory effect on NO- mediated apoptosis signaling. Thereby these two agents seem to interact in an antagonistic mode. In addition, under specific steric conditions a POD-dependent effect, potentially driven by H_2_O_2_ interferes with NO/peroxynitrite signaling as well. Dependent on its concentration, catalase may abrogate the antagonism between H_2_O_2_ and NO or interfere with NO/peroxynitrite signaling.

### The role of ferrous/ferric ions for the antagonistic action between NO and H_2_O_2_ and vice versa

Established reactions between ferrous/ferric ions with NO [68–75] as well as with hydrogen peroxide [76, 77]) were the basis for the speculation that iron might play a crucial role for the antagonistic effects between NO and hydrogen peroxide and vice versa. For a direct experimental verification or falsification of this hypothesis, it was tested, whether the removal of iron by the iron chelator deferoxamine would abrogate the modulatory effects of a) exogenous NO on H_2_O_2_-dependent HOCl-mediated apoptosis induction, b) exogenous NO on inhibition of HOCl-mediated apoptosis induction by excess H_2_O_2_ and c) H_2_O_2_ on NO-dependent, peroxynitrite-mediated apoptosis induction.

Treatment of MKN-45 gastric carcinoma cells with 100 mM 3-AT allowed for reactivation of HOCl-mediated apoptosis-inducing signaling, as seen by the strong inhibitory effect of the HOCl scavenger taurine and the minor effect of peroxynitrite decomposition catalyst FeTPPS (Figure 12 A). Addition of the exogenous NO donor SNP caused inhibition of HOCl-mediated apoptosis induction in a concentration-dependent mode. Inhibition was complete at 62 µM SNP. Further increase in the SNP concentration caused resumption of NO/peroxynitrite-mediated apoptosis induction, as shown by the inhibitory effect of FeTPPS at higher concentrations of SNP. The HOCl scavenger taurine not only prevented HOCl-mediated apoptosis induction, but it also increased the efficiency of NO/peroxynitrite-mediated apoptosis induction, leading to a left-ward shift of the dose response curve for NO-dependent peroxynitrite signaling.

**Figure 12.**
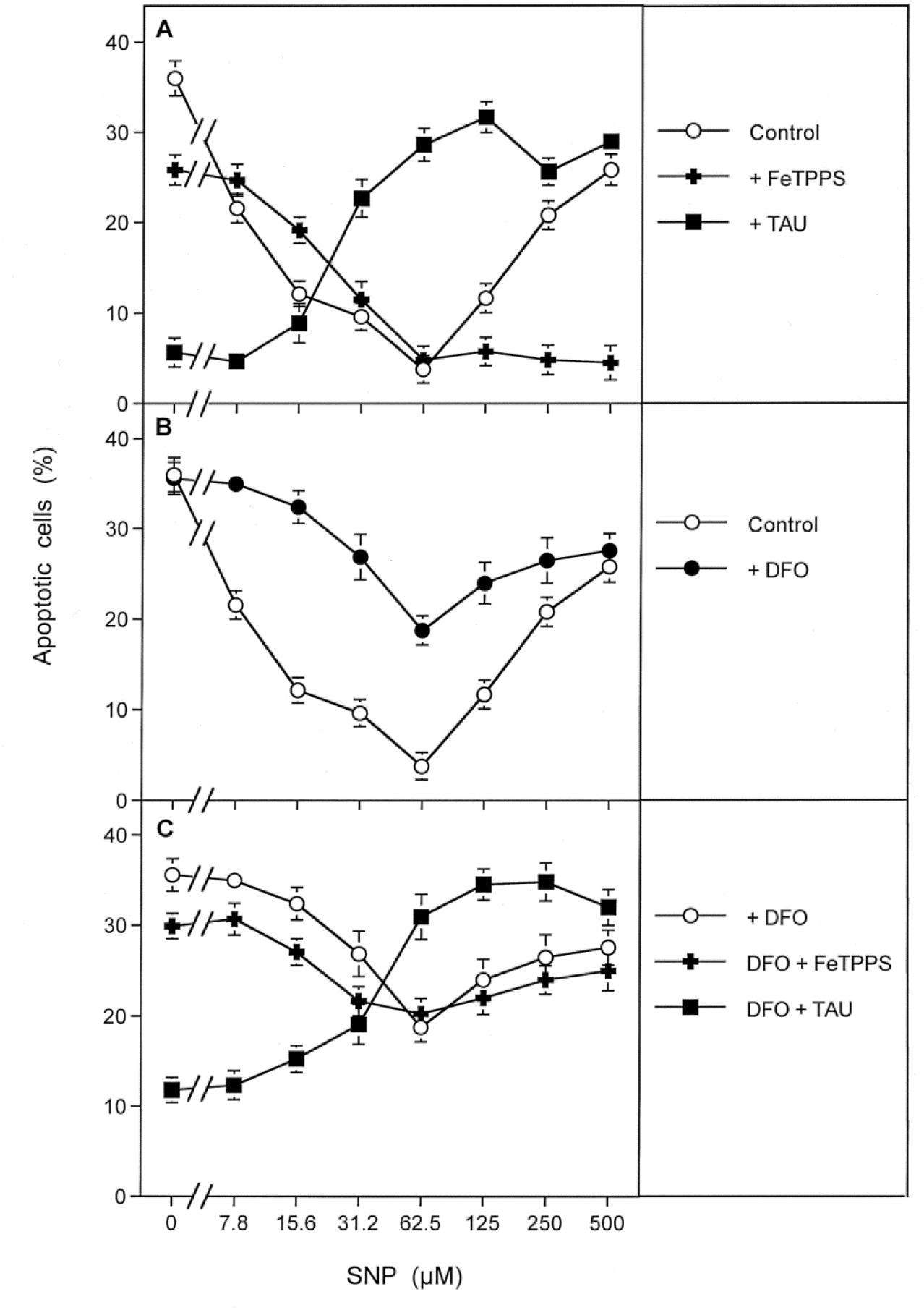
The iron chelator deferroxamine interferes with the NO-mediated antagonism towards H_2_O_2_-dependent HOCl signaling. 12 500 MKN-45 cells/assay were cultivated in the presence of 100 mM 3-AT and received either no inhibitors (“control”) or 25 µM FeTPPS, 50 mM taurine or 50 µM DFO. SNP was added at increasing concentrations and the percentages of apoptotic cells were determined after 4 h. A. Apoptosis induction in the presence of 3-AT was due to HOCl signaling. Increasing concentrations of the NO donor SNP first caused NO-dependent, peroxynitrite-independent inhibition of HOCl signaling, followed by resumption of NO/peroxynitrite signaling, as deduced from the inhibition profiles. Note that the presence of the HOCl scavenger taurine caused a leftward shift of the curve indicating NO/peroxynitrite signaling. This effect of the inhibitor is explained by the role of HOCl in iron-mediated consumption reactions according to reaction #16 m(and following) in Figure 19. B. Removal of iron from the system causes a strong abrogation of the effects of the NO donor. C. In the presence of DFO, both pathways are active. *Statistical analysis: The effects of SNP and the counteraction by DFO are highly significant (p<0.001)*.

The iron chelator DFO strongly interfered with the inhibition of HOCl-mediated apoptosis induction by the NO donor (Figure 12 B), pointing to a central role of iron in NO-mediated inhibition of H_2_O_2_-dependent, HOCl-mediated apoptosis induction. Thereby the degree of interference by DFO was inversely related to the concentration of the NO donor. For example, 7.7 and 15.5 µM SNP caused 44 and 72 % inhibition of HOCl-mediated apoptosis induction. This inhibitory effect of SNP was nearly completely counteracted by DFO (93 and 82.6 % abrogation of inhibition). In contrast, the higher concentration of SNP (62 µM) inhibited HOCl-mediated apoptosis induction to background activity. This inhibitory effect was counteracted by DFO only to 45 %.

Inhibition of HOCl signaling by excess free H_2_O_2_ was abrogated through the application of the NO donor SNP (Supplementary Figure S 7). After optimal HOCl synthesis was achieved, it was inhibited by further increase in the concentration of the NO donor. Both of these NO-dependent processes directed towards H_2_O_2_ were abrogated by DFO, thus pointing to a central role of iron in this process.

NO-dependent, peroxynitrite-mediated apoptosis induction in the tumor cell line MKN-45 mediated by an exogenous NO donor requires inhibition of protective catalase. 10 mM 3-AT were required and sufficient to allow substantial apoptosis induction in the presence of the NO donor. Apoptosis induction was dependent on the concentration of the fast decaying NO donor DEA NONOate. Inhibition of NOX1 by AEBSF or decomposition of peroxynitrite by FeTPPS completely inhibited DEA NONOate-dependent apoptosis induction, thus confirming that NO/peroxynitrite signaling had been active in the absence of scavengers (Figure 13 A). The increase in free tumor cell-derived H_2_O_2_ through increasing the concentration of the catalase inhibitor 3-AT from 10 mM to 50 mM caused a strong rightward shift of the concentration/response curve (Figure 13 B). This is fully explained by consumption of NO by free H_2_O_2_. This shift was completely reverted when either the generation of H_2_O_2_ was slowed down by mild inhibition of NOX1 through 5 µM AEBSF, or through application of 50 µM of the iron chelator DFO. These findings point to a consuming role of H_2_O_2_ and iron on the effective concentration of NO.

**Figure 13.**
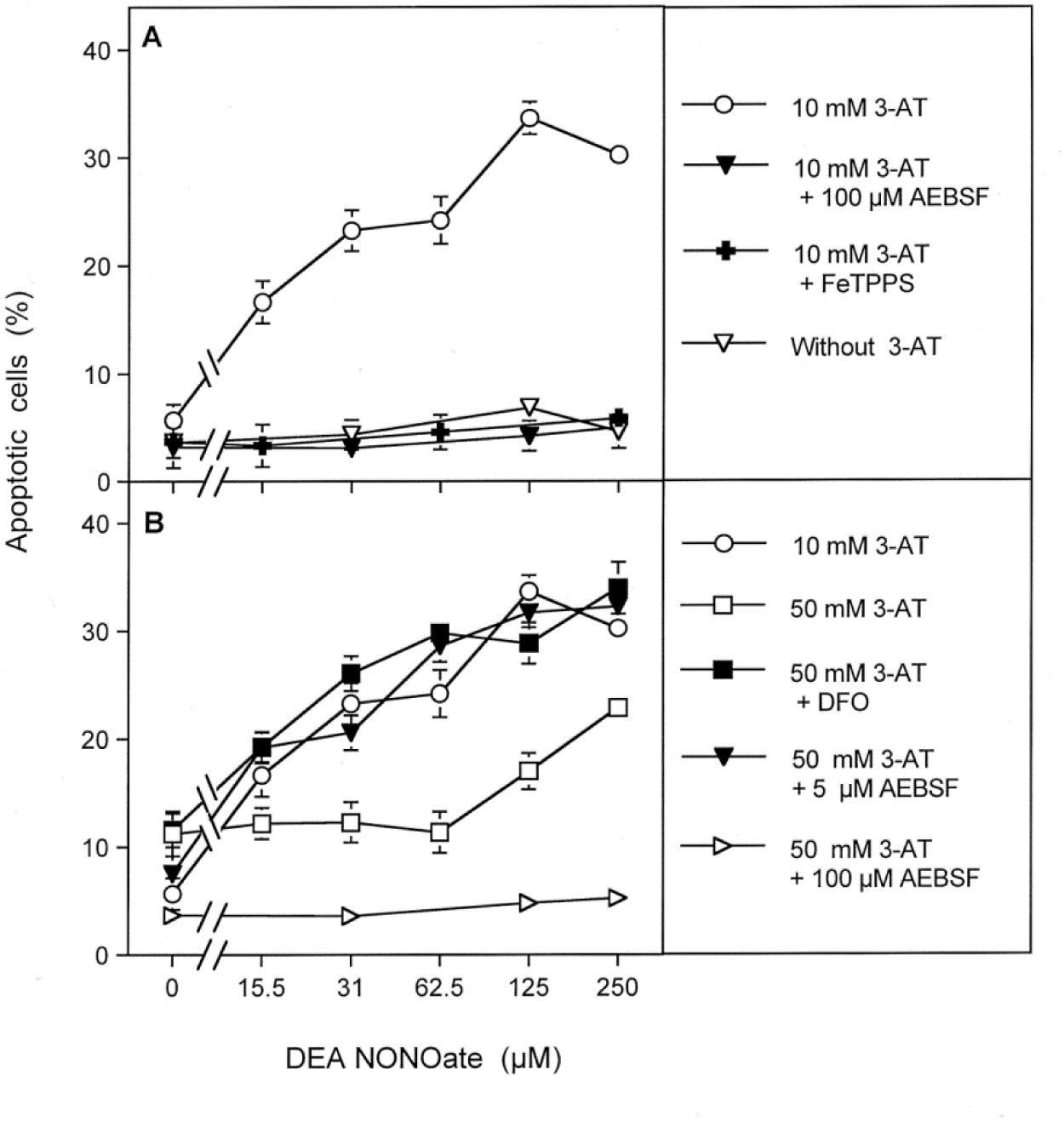
The iron chelator deferroxamine interferes with the antagonistic effect of H_2_O_2_ towards NO-mediated, peroxynitrite-dependent apoptosis-inducing signaling. 5 000 MKN-45 cells/assay remained without 3-AT or received either 10 mM 3-AT (A) or 50 mM 3-At (B). In addition, where indicated, AEBSF was added at a concentration of either 5 µM or 100 µM, FeTPPS at a concentration of 25 µM and DFO at a concentration of 50 µM. DEANONOate was added at increasing concentrations and the percentages of apoptotic cells were determined after 3 h. A. In the presence of 10 mM 3-AT, the NO donor DEANONOate causes concentration-dependent induction of apoposis (p<0.001), which is completely inhibited by the NOX-inhibitor AEBSF (p<0.001) and the peroxynitrite decomposition catalyst FeTPPS (p<0.001), confirming that apoptosis induction was due to the action of peroxynitrite. B. An increase in the concentration of 3-AT up to 50 mM first caused inhibition of DEAENONOate-mediated apoptosis compared to the situation for 10 mM 3-AT (p<0.001), before the curve resumed at concentrations of DEAENONOate higher than 62.5 µM. These effects of 50 mM 3-AT compared to 10 mM 3-AT are explained by an increased concentration of tumor cell-generated H_2_O_2_, which leads to consumption of NO. Gradual inhibition of NOX-1 with the suboptimal concentration of 5 µM AEBSF abrogates the effects of 50 mM 3-AT (p<0.001) through decreasing NOX-1 activity and subsequent dismutation of superoxide anions to H_2_O_2_. DFO also interferes with the effect of 50 mM 3-AT (p< 0.001).by abrogation of the iron-catalyzed consumption of NO by H_2_O_2_.

In line with these findings, NO-mediated, peroxynitrite-dependent apoptosis induction was inhibited by increasing concentrations of H_2_O_2_-generating glucose oxidase. This inhibitory effect was completely abrogated by DFO (Supplementary Figure S8) at all concentrations of GOX

### H_2_O_2_ antagonizes apoptosis induction by preformed peroxynitrite

Exogenous application of chemically synthesized peroxynitrite allowed to study the effect of tumor cell-derived H_2_O_2_ on apoptosis induction by preformed exogenous peroxynitrite. This experimental set-up therefore did not require intermediate NO/superoxide anion interaction in the experimental system, as utilized in previous experiments. Also, in the experimental setting where preformed peroxynitrite was added, this compound was present at all sites of the test system, in contrast to its more restricted localization to the membrane in the case of its generation through superoxide anions and NO.

MKN-45 cells were treated with increasing concentrations of neutralizing antibody directed towards their protective catalase, in the absence or presence of the NOX1 inhibitor AEBSF. The cells were then challenged with 12.5 µM or 25 µM peroxynitrite, and apoptosis induction was subsequently determined. As can be seen from Figure 14, MKN-45 cells without antibody treatment or treated with irrelevant control antibody were not affected by peroxynitrite. This finding confirms the protective capacity of their membrane-associated catalase against exogenous peroxynitrite [28, 29]. Increasing concentrations of neutralizing antibody directed towards catalase allowed an initial increase in apoptosis induction by exogenous peroxynitrite, followed by gradual decline of the apoptotic response with further increasing concentration of the antibodies. The resultant optimum curve was broader for the higher concentration of peroxynitrite. This is in line with a consumption reaction between peroxynitrite and H_2_O_2_. When the generation of superoxide anions and their dismutation product H_2_O_2_ had been prevented through AEBSF-mediated inhibition of NADPH oxidase, the dependence of apoptosis induction by peroxynitrite on the concentration of catalase inhibiting antibodies changed from an optimum curve to a plateau curve. This indicates that H_2_O_2_ generated by the cells (through dismutation of extracellular, NOX- derived superoxide anions) at high concentrations of antibodies against catalase had caused the inhibition of the peroxynitrite effect. A negative effect of superoxide anions on peroxynitrite was directly ruled out, as AEBSF did not affect peroxynitrite-mediated apoptosis induction in the lower concentration range of antibodies. If superoxide anions would have directly interfered with peroxynitrite, such interference should have been especially prominent at low concentrations of antibodies and the resultant low concentration of effective peroxynitrite (due to suboptimal prevention of peroxynitrite decomposition by catalase under these conditions. The interaction between peroxynitrite and H_2_O_2_ under these specific experimental conditions can be explained by the findings by Di Mascio et al. [78], extended by previous own work [43, 47]

**Figure 14.**
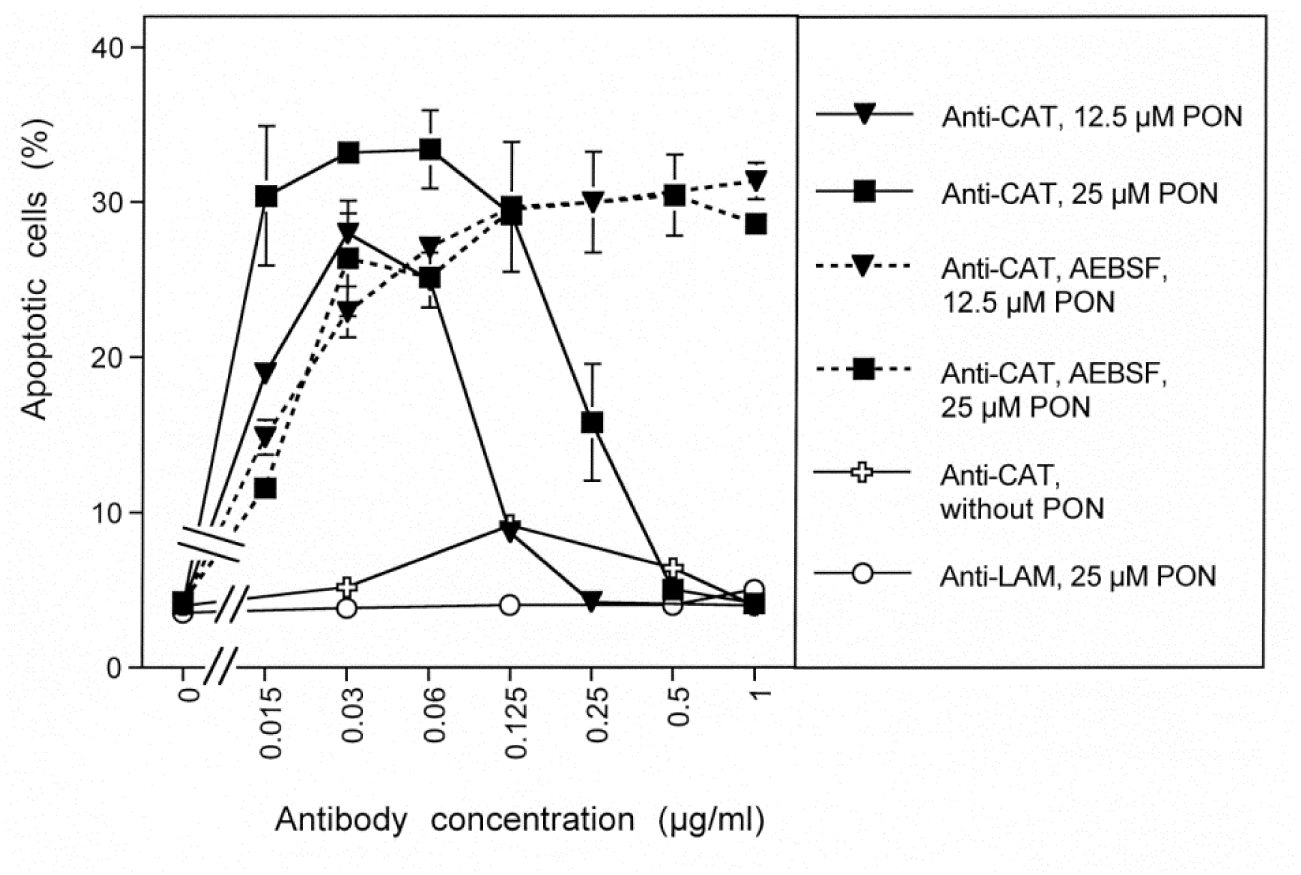
Antagonism between H_2_O_2_ and peroxynitrite. 6000 MKN-45 cells/100 µl were treated with the indicated concentrations of neutralizing monoclonal antibody directed against catalase for 20 min. Control assays received anti-laminin as irrelevant control antibody. Where indicated, NOX1 was completely inhibited by 100 µM AEBSF. Assays remained either free of peroxynitrite or received 12.5 or 25 µM peroxynitrite. Apoptosis was measured after 1.5 h. Control assays without antibody and without peroxynitrite were indistinguishable from assays containing either specific or nonspecific antibody and that were not treated with peroxynitrite. The experiment shows that the tumor cells are protected towards exogenous peroxynitrite by their membrane-associated catalase. Specific antibody-mediated inhibition of membrane-associated catalase sensitized the cells for apoptosis induction by peroxynitrite, dependent on the concentration of antibodies. At higher antibody concentrations, the effect of peroxynitrite was antagonized. This antagonistic effect seemed to be caused by cell-derived H_2_O_2_, as it increased with the concentration of antibody directed against catalase and was abrogated when superoxide anion production was blocked by AEBSF. *Statistical analysis: Apoptosis induction by 12.5 µM peroxynitrite in the presence of 0.015-0.06 µg/ml anti-catalase, apoptosis induction by 25 µM peroxynitrite in the presence of 0.015-0.25 µg/ml anti-catalase and the enhancing effect of AEBSF at concentrations of anti-Catalase higher than 0.06 µg/ml (in the case of 12.5 µM peroxynitrite) and of anti-catalase higher than 0.125 µg/ml (in the case of 25 µM peroxynitrite) were highly significant (p < 0.001). Anti-laminin did not induce significant sensitivity for peroxynitrite-dependent apoptosis induction, nor did anti-catalase cause apoptosis induction in the absence of peroxynitrite*.

### Modulation of intercellular ROS/RNS-dependent apoptosis-inducing signaling

The mutual interrelationship between H_2_O_2_ and NO allowed to predict that modulation of the concentrations of critical reaction partners like superoxide anions by the NOX1 inhibitor AEBSF, H_2_O_2_ by the catalase mimetic EUK-134 or NO by the NOS inhibitor L-NAME should have distinct consequences for the quality, as well as efficiency of intercellular ROS/RNS signaling of tumor cells. Thereby, increasing cell density and an increasing rate of extracellular superoxide anion production mention dismutation under discussion should have a positive effect on HOCl signaling up to an optimum, followed by H_2_O_2_-mediated inhibitory effects. In contrast, a negative effect of increasing cell density on NO/peroxynitrite signaling was to be expected [8, 55 79]. These initial findings were systematically analysed in the experiment described in Figure 15. MKN-45 cells at relatively low cell density (62 500 cells/ml) responded to increasing concentrations of 3-AT with apoptosis induction in the mode of a plateau curve (Figure 15 A). Inhibitor studies under these conditions showed a broad initial range of NO/peroxynitrite signaling, followed by HOCl signaling. Increasing the cell density to our standard density of 125 000 cells/ml, changed the concentration-response curve to an optimum curve (Figure 15 B). In parallel, the initial concentration range of 3-AT that preferentially allowed NO/peroxynitrite signaling became more narrow, whereas HOCl signaling was first increased and then inhibited by higher concentrations of 3-AT. Further doubling of the cell density to 250 000 cells/ml completely abrogated detectable NO/peroxynitrite signaling and favoured exclusive HOCl signaling (Figure 15 C). At concentrations of 3-AT above 100 mM apoptosis induction seemed to be independent of HOCl and NO/peroxynitrite signaling at this high cell density, most likely due to direct apoptosis induction by very high concentrations of H_2_O_2_.

**Figure 15.**
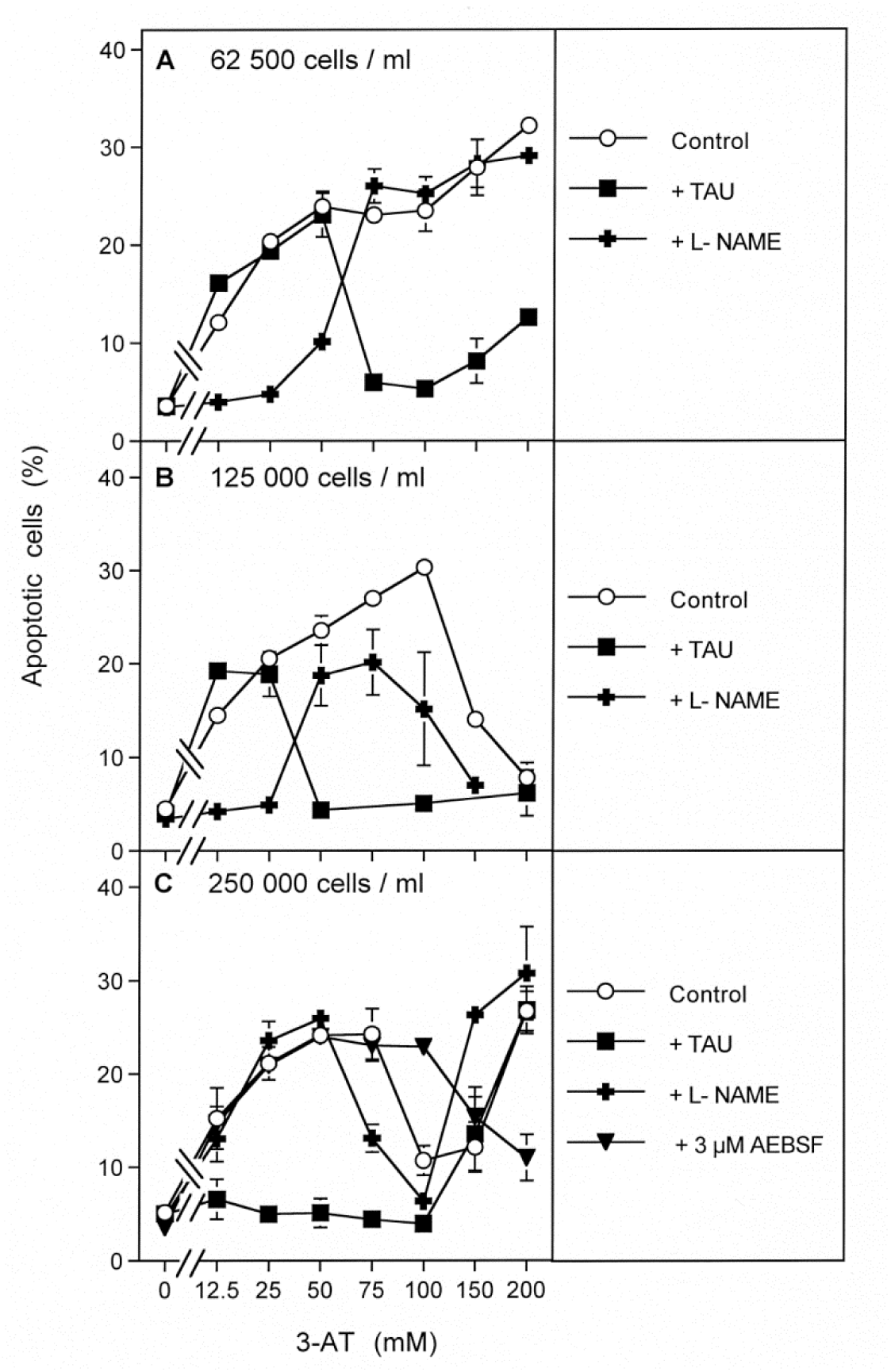
Modulation of intercellular ROS/RNS signaling by the density of tumor cells. MKN-45 tumor cells at the indicated densities were seeded in 96 well plates (100 µl volume). The assays either received no inhibitor (“control”), or 50 mM of the HOCl scavenger taurine, 2.4 mM of the NOS inhibitor L-NAME or 3 µM of the NOX inhibitor AEBSF. The catalase inhibitor 3-AT was added at the indicated concentrations and the percentages of apoptotic cells were determined after 6.5 h. This experiment shows that at the lowest cell density (A), apoptosis induction follows the mode of a plateau curve, with NO/peroxynitrite signaling up to 50 mM 3-AT, followed by HOCl signaling. At the intermediate cell density (B), the apoptotic response of the cells is characterized by an optimum curve. The concentration range that allows specifically NO/peroxynitrite signaling is more narrow than under A. At the highest cell density (C), the optimum curve is more narrow than under B and is essentially due to HOCl signaling. It is followed by apoptosis induction that is independent of HOCl- or NO/peroxynitrite signaling. *Statistical analysis: A. Apoptosis induction mediated by 3-AT, as well as its inhibition by L-NAME in the lower concentration range of 3-AT and by taurine in the higher concentration range are highly significant (p<0.001). B. The change to an optimum curve at higher concentrations of 3-AT and the differential effects of the inhibitors are highly significant (p<0.001). C. The change in signaling chemistry compared to B is highly significant (p<0.001)*.

The next series of experiments analysed the signaling chemistry of a dense population of tumor cells at increasing concentrations of 3-AT, when i) the superoxide anion production was gently reduced through mild inhibition of NOX1 by AEBSF, or ii) excess H_2_O_2_ was removed by fine-tuned concentrations of the catalase mimetic EUK-134 or iii) the NO donor SNP was added at low concentrations. Figure 16 A shows that the dense population of tumor cells responded to increasing concentrations of 3-AT exclusively with HOCl signaling and in the mode of an optimum curve. NO/peroxynitrite signaling did not contribute to apoptosis induction, as seen from the lack of inhibition by the NOS inhibitor L-NAME. When the cells were treated with low concentrations of the NOX1 inhibitor AEBSF (Figure 16 B, C), i) the optimum curve was changed to a plateau-type response curve and ii) NO/peroxynitrite signaling was reactivated in the lower concentration range of 3-AT while HOCl signaling was maintained in the subsequent higher concentration range of 3-AT.The concentration range in which HOCl signaling was substituted by NO/peroxynitrite signaling was dependent on the concentration of AEBSF. Removal of excess H_2_O_2_ through application of discrete concentrations of the catalase mimetic EUK-134 caused an effect analogous to that of suboptimal inhibition of superoxide anion generation by AEBSF (Figure 16 D, E), i. e. it changed the curve to a plateau-like form and reactivated NO/peroxynitrite signaling on the expense of HOCl signaling. At 2 µM EUK-134, NO/peroxynitrite signaling was dominant up to 75 mM 3-AT and HOCl signaling was only detectable at 100 mM 3-AT.

**Figure 16.**
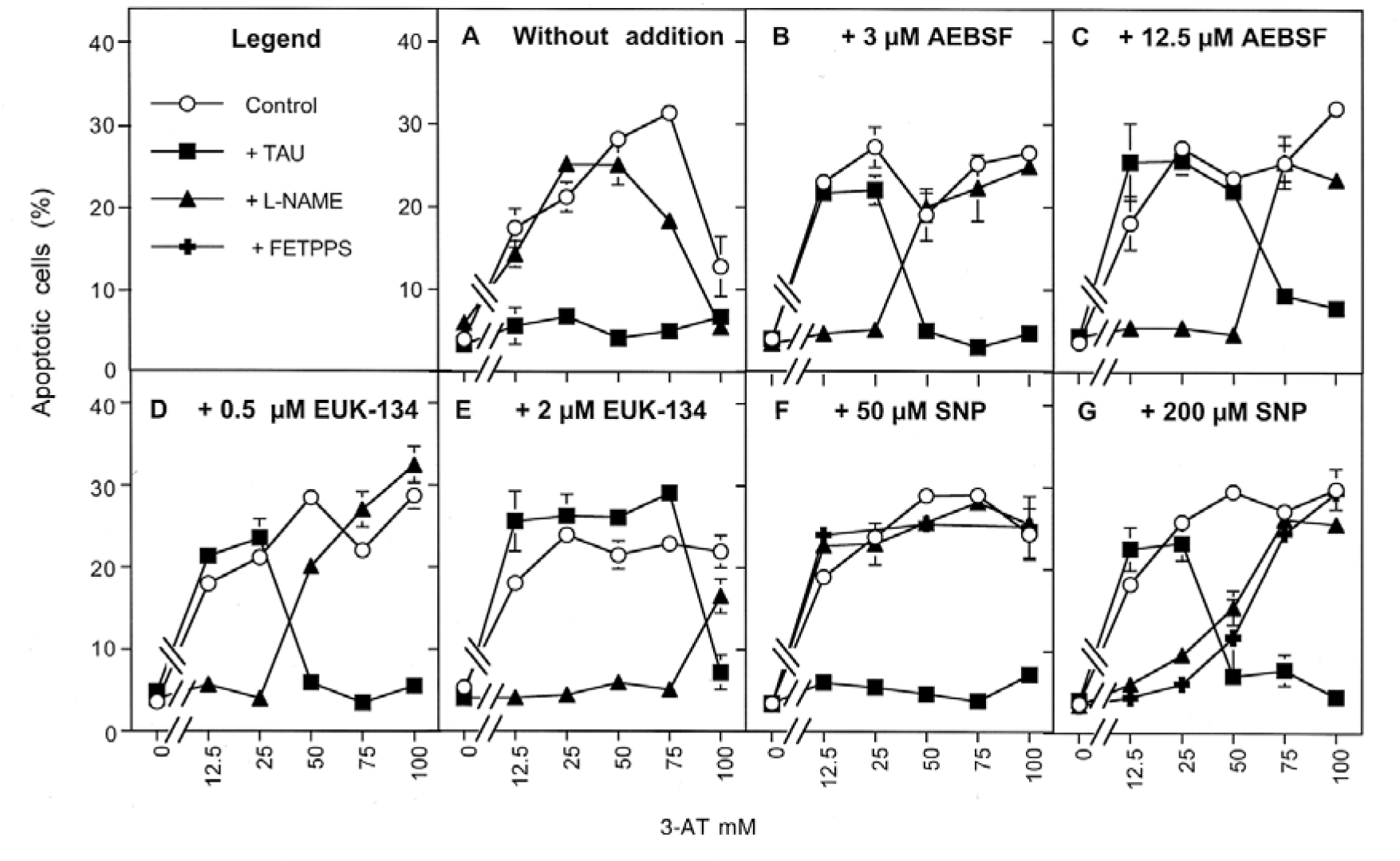
Modulation of apoptosis induction through partial inhibition of NOX1, mild removal of H_2_O_2_ or addition of an NO donor. MKN-45 cells (250 000 cells/ml) in 100 µl assays received either no inhibitors (“control”), 50 mM of the HOCl scavenger taurine (TAU), 2.4 mM of the NOS inhibitor L-NAME, or 25 µM of the peroxynitrite decomposition catalyst FeTPPS. Assays remained without further addition (A), 3µM or 12.5 µM of the NOX inhibitor AEBSF (B, C), 0.5 µM or 2 µM of the catalase mimetic EUK-134 (D, E) or 50µM or 200 µM of the NO donor SNP (F, G). The catalase inhibitor 3-AT was added at increasing concentrations. The percentages of apoptotic cells were determined after 5 h. The results show that 3-AT induces apoptosis in the dense culture in the mode of an optimum curve and essentially dependent on HOCl signaling (A) (p<0.001). Low concentrations of the NOX inhibitor AEBSF (B,C) shifted the optimum curve to a plateau curve (p<0.001). and allowed for selective NO/peroxynitrite signaling at lower concentrations of 3-AT (p<0.001). The concentration range of 3-AT in which NO/peroxynitrite signaling was allowed, was dependent on the concentration of AEBSF (p<0.001). Low concentrations of the catalase mimetic EUK-134 (D, E) shifted the optimum curve also to a plateau curve and restored NO/peroxynitrite signaling up to 25 mM 3-AT (D) (p<0.001) or 75 mM 3-AT (E) (p<0.001). 50 µM SNP (F) shifted the curve to a plateau curve (p<0.001)., which was completely dependent on HOCl signaling (p<0.001), wherease 200 µM SNP (G) also allowed for NO/peroxynitrite signaling up to 25 mM 3-AT (p<0.001).

Addition of 50 µM of the NO donor SNP caused a change of the optimum curve to a plateau curve, while HOCl signaling was still maintained over the complete concentration range of 3-AT (Figure 16 F). This finding shows that under these conditions exogenous NO had interfered with H_2_O_2_-mediated inhibition of HOCl signaling, but had not inhibited HOCl signaling or replaced it by NO/peroxynitrite signaling. Increasing the concentration of the NO donor up to 200 µM then allowed for reactivation of NO/peroxynitrite signaling, which was followed by HOCl signaling at 50 mM 3-AT and higher concentrations (Figure 16 G). Importantly, the findings presented in Figure 15 show that reactivated NO/peroxynitrite signaling was always driven by NOS-derived NO, as seen from the inhibition by L-NAME. Even if NO/peroxynitrite signaling was achieved by the exogenous NO donor SNP, the resultant signaling was due to endogenous NO. This points to the modulating role of SNP. The results demonstrated in Figures 15 and 16 confirm the negative impact of H_2_O_2_ on a) NO/peroxnitrite signaling and on b) HOCl signaling at high concentrations of H_2_O_2_, as both effects can be abrogated by mild reduction of the H_2_O_2_ concentration. The data show that a mild reduction of the H_2_O_2_ concentration can be efficiently achieved through mild inhibition of the activity of NOX1 (thus lowering the dismutation reaction), through discrete decomposition of H_2_O_2_ by a catalase mimetic or through controlled application of an NO donor. These findings therefore point to consumption of H_2_O_2_ by free NO and vice versa.

Based on these findings, it was obvious that it should be possible to rationally modulate the signaling quality of tumor cells through modulating the concentrations of central signaling-relevant molecules. As shown in Figure 17 A, C, MKN-45 gastric carcinoma cells at a density of 125 000 cells/ml responded to increasing concentrations of 3-AT with apoptosis induction in the mode of an optimum curve. Apoptosis was determined by initial NO/peroxynitrite signaling, followed by HOCl signaling. A slight inhibition of NOX1 activity through 10 µM AEBSF, in combination with addition of 0.75 mM of the NOS substrate arginine, caused a shift to exclusive NO/peroxynitrite signaling and followed the mode of a plateau-type curve (Figure 17 B). In a complementary experiment, the increase in superoxide anion production [79, 80] and POD release [81] through the addition of exogenous TGFß1 caused a shift to mere HOCl signaling (Figure 17 D). These findings might be the clue for the understanding of the variable signaling profiles between different groups of tumor cells that are illustrated in the next chapter.

**Figure 17.**
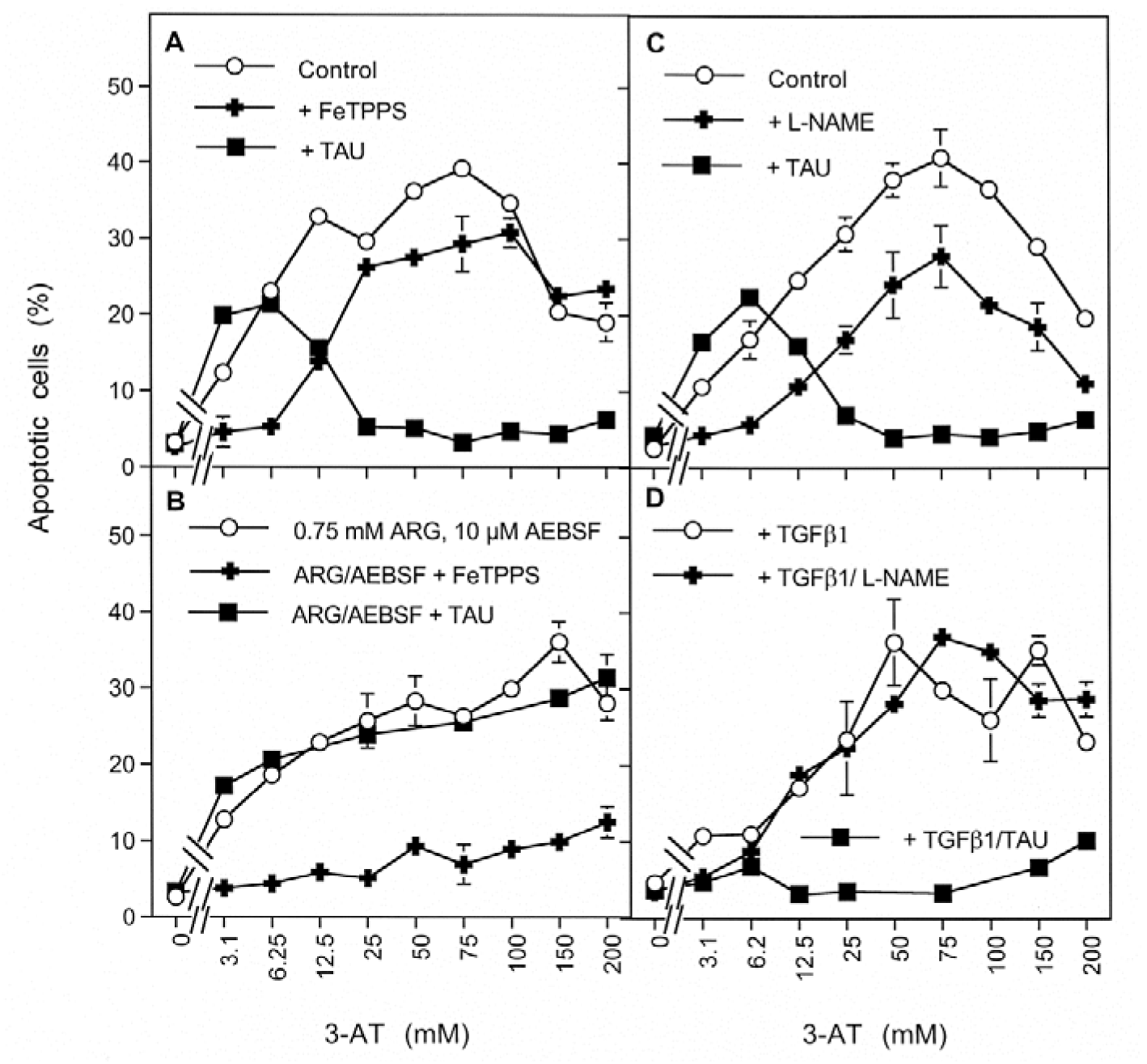
Modulation of intercellular ROS/RNS signaling through modulation of the concentrations of tumor cell-derived superoxide anions and NO. A, C: 12 500 MKN-45 cells per 100 µl assay received no addition (“control”), or 25 µM of the peroxynitrite decomposition catalyst FeTPPS, 2.4 mM of the NOS inhibitor L-NAME or 50 mM of the HOCl scavenger taurine (TAU), as indicated. B: The assays received 0.75 mM arginine (ARG) plus 10 µM AEBSF, either alone or in combination with 25 µM FeTPPS or 50 mM taurine. D: The assays received 20 ng/ml TGF-beta-1, either alone or in combination with 2.4 mM L-NAME. The catalase inhibitor 3-AT was added at the indicated concentrations and the percentages of apoptotic cells were determined after 4.5 h. The results under A and C show that the tumor cells at the selected density responded to 3-AT in the mode of an optimum curve (p<0.001), characterized by initial NO/peroxynitrite signaling (p<0.001), followed by HOCl signaling (p<0.001). B: The parallel increase in cell derived NO through addition of arginine, in combination with mild inhibition of superoxide anion generation by 10 µM AEBSF, shifted the curve to a plateau curve of apoptosis induction (p<0.001) that was completely dependent on NO/peroxynitrite signaling (p<0.001). D. Enhancement of NOX1 activity by TGF-beta1 (Temme and Bauer, 2013) shifted apoptosis induction completely to HOCl signaling (p<0.001) and in the mode of an optimum curve (p<0.001).

### Variability of the quality of intercellular ROS/RNS-dependent apoptosis-inducing signaling of human tumor cell lines

Human tumor cell lines have been found to show different signaling profiles after addition of increasing concentrations of either the catalase inhibitor 3-AT or neutralizing antibodies directed towards catalase [28]. After inhibition of membrane-associated catalase, tumor lines like L929 or Gumbus essentially reactivated HOCl signaling, MKN-45 cells initially reactivated NO/peroxynitrite signaling followed by HOCl signaling at higher degrees of catalase inhibition, whereas lines like SKNMC showed NO/peroxynitrite signaling exclusively. This type of analysis is extended in Supplementary Figure S9. Again, certain tumor cells show NO/peroxynitrite signaling at lower concentrations of the catalase inhibitor, followed by HOCl signaling at higher concentrations of the inhibitor, whereas other cell lines seem to be dependent on NO/peroxynitrite signaling at all concentrations of the catalase inhibitor. When optimal superoxide anion production was warranted through addition of exogenous TGF-beta1, tumor cell lines like SISO or SIHA (cervical carcinoma) and the melanoma cell lines IGR-37 and IGR-39 showed mainly HOCl signaling in the presence of 100 mM 3-AT (Supplementary Figure S10), whereas the mammary carcinoma cell lines CAL-51m HCC-1937 and MCF-7, as well as the ovarial carcinoma cell line FU-OV-1 showed NO/peroxynitrite signaling exclusively under these conditions (Supplementary Figure S11).

When clumps of transformed “target cells” were surrounded by high numbers of “effector cells” (an experimental setting recently described by Pottgiesser et al., 2015; please see Supplementary Figure 12 for details), tumor cells such as SISO and SIHA that responded with HOCl signaling after optimal catalase inhibition also caused apoptosis induction through HOCl signaling in the transformed target cells. In contrast, tumor cells such as CAL-51, Fu-OV-1, BG-1, that preferentially showed NO/peroxynitrite signaling after inhibition of their protective catalase caused apoptosis induction through the NO/peroxynitrite pathway in the transformed target cells (Supplementary Figure S13). These findings are indicative for differential release of POD versus NO from different tumor cell lines. (Importantly, tumor cell lines from both categories of effector functions (i. e. mediating either NO/PON or HOCl signaling in target cells) as determined in the previous figures, were able to respond to NO/peroxynitrite-mediated apoptosis induction after application of an exogenous NO donor, provided their protective catalase was inhibited (Supplementary Figures S14 and S15)).

Finally, the requirements for specific apoptosis induction in tumor cells through intercellular ROS/RNS signaling and the determinants for preferential NO/peroxynitrite or HOCl signaling were analysed through the use of SHEP neuroblastoma and SISO cervical carcinoma cells. These two cell lines were chosen as under optimal conditions for autocrine apoptosis induction and in the presence of a catalase inhibitor, SHEP cells died through NO/peroxynitrite signaling and SISO cells exclusively through HOCl signaling. The analysis performed here combined clumps of target cells (low number, high density) surrounded by effector cells (high number, low density), following the approach described in Supplementary Figure 12. This approach allowed to study homologous and heterologous combinations of target and effector cells from these two distinct tumor cell populations (Supplementary Figures 16-24). The results confirm that inhibition or knockdown of protective catalase is absolutely essential to exert the target cell function, but is not necessary for the effector function. Furthermore, the data show that the distinct effector function of the effector cells determines whether the target cells die by NO/peroxynitrite or HOCl signaling. Thererfore, clumps of either SISO or SHEP target cells died from HOCl signaling, when their catalase was inhibited and they were cocultured with excess SISO effector cells. Vice versa, clumps of either SISO or SHEP target cells died from NO/peroxynitrite signaling, when they were cocultured (in the presence of catalase inhibitor) with excess SHEP effector cells (Supplementary Figures 16, 17). The activity of the effector cells was completely mimicked by addition of either MPO or the NO donor SNP to target cells (Supplementary Figures 18, 19), indicating that peroxidase and NO were the discriminative and sufficient effector molecules in this system. Therefore, siRNA-mediated knockdown of DUOX in SISO cells and nNOS in SHEP cells, but not the reciprocal approach, completely abrogated their effector functions. Knockdown of DUOX or NOS in target cells did not affect their target cell function, whereas knockdown of NOX1 in target cells completely prevented apoptosis induction in target cells. This finding confirms that NOX1-derived superoxide anions are the crucial element for efficient and selective target cell function of tumor cells, provided their catalase is inactivated or knocked down (Supplementary Figure 22, 23).

In line with the antagonistic actions of NO and H_2_O_2,_ as characterized in this manuscript, apoptosis induction in tumor cells with preferential HOCl signaling was initially inhibited by exogenous NO, before NO/peroxynitrite signaling was overt at higher concentrations of the NO donor (Supplementary Figures S25, S26 A). In contrast, apoptosis induction in NO/peroxynitrite signaling-dominated tumor cells was enhanced by exogenous NO (Supplementary Figure S26 B).

## Discussion

This study confirms the chemical biology and the multiple interconnections of the apoptosis-inducing HOCl signaling pathway (Figure 18 A) and the NO/peroxynitrite signaling pathway (Figure 18 B). Both pathways share the dependency on NOX1-derived extracellular superoxide anions (reactions #1 and #3 under A; reaction #5 in B) and are finalized by the generation of hydroxyl radicals that cause lipid peroxidation and subsequent induction of the mitochondrial pathway of apoptosis (reaction #4 under A, reaction #8 under B) [16]. These pathways differ markedly from each other with respect to the central roles of either peroxidase/HOCl or NO/peroxynitrite [6, 7, 10, 11].

**Figure 18.**
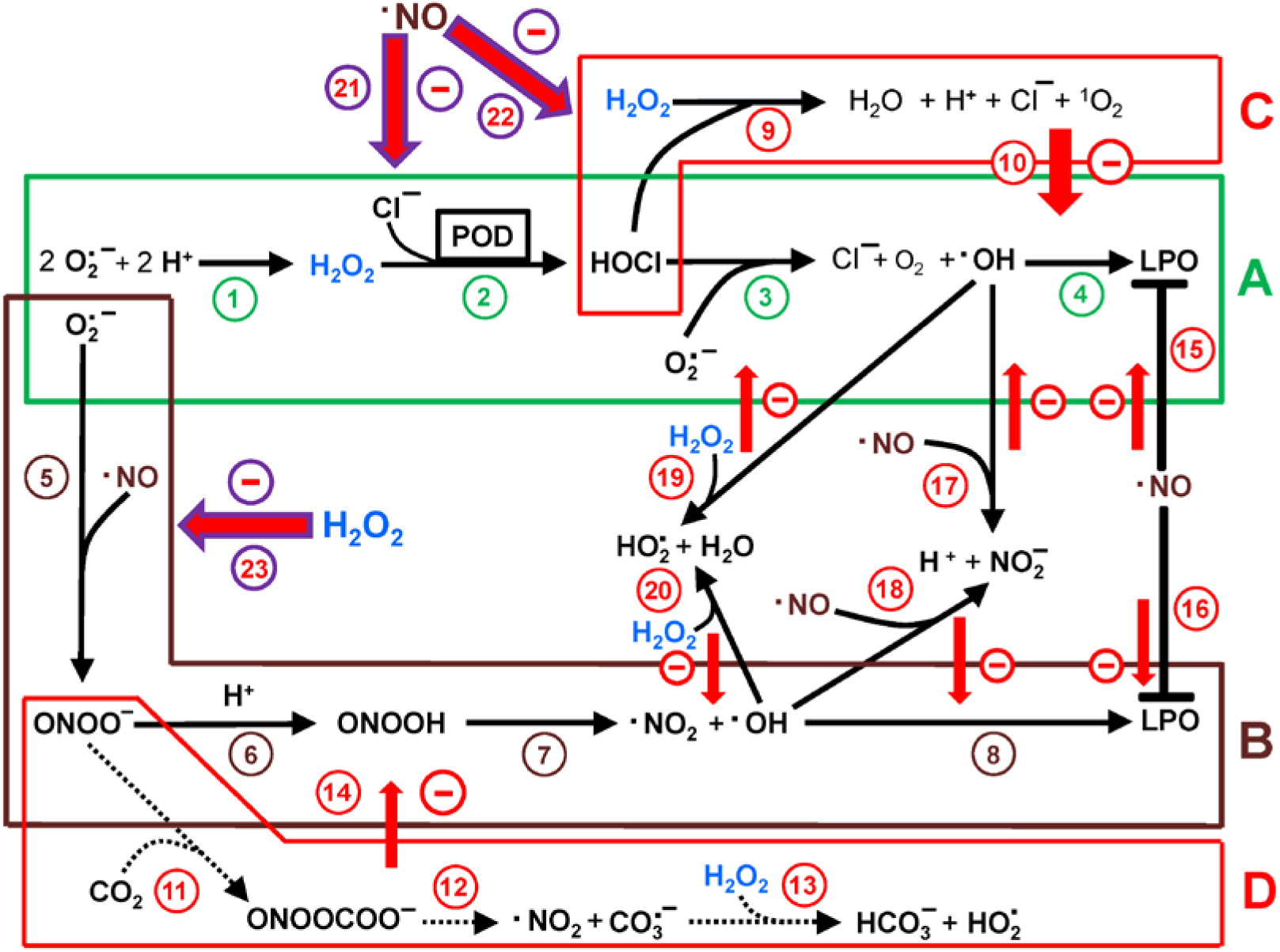
The effect of NO and H_2_O_2_ on intercellular ROS/RNS-dependent apoptosis-inducing signaling. *A. HOCl signaling*. The dismutation of superoxide anions generated by NOX1 (#1) leads to the formation of H_2_O_2_. DUOX-coded peroxidase (POD) uses H_2_O_2_ as substrate for HOCl synthesis (#2). The interaction between HOCl and superoxide anions (#3) causes the generation of hydroxyl radicals, which may cause lipid peroxidation (LPO) (#4) and subsequent induction of the mitochondrial pathway of apoptosis. *B. NO/peroxynitrite signaling*. The reaction between superoxide anions and NO leads to the formation of peroxynitrite (#5). Protonation of peroxynitrite (#6) by proton pumps in the cell membrane leads to the formation of peroxynitrous acid (#7), which spontaneously decomposes into hydroxyl radicals and NO_2_ (#6). This causes lipid peroxidation (LPO) (#8) and induction of apoptosis. *C. The effect of excess H_2_O_2_ on HOCl signaling*. Excess H_2_O_2_ interacts with HOCl (#9) and in this way blunts HOCl signaling (#10). *D. Consumption of peroxynitrite by CO_2_.* CO_2_ efficiently consumes peroxynitrite through reactions #11-13. This reaction has a negative effect on NO/peroxynitrite signaling (#14). Additional negative interactions: NO has the potential to react with lipid peroxides (#15, 16) and hydroxyl radicals (#17, 18). H_2_O_2_ can react with hydroxyl radicals (#19, 20). Reactions #15-20 might negatively interfere with HOCl- or NO/peroxynitrite signaling. The focus of our analysis are the negative effects of NO on HOCl signaling (#21), the consumption of H_2_O_2_ through NO, followed by abrogation of H_2_O_2_-mediated inhibition of HOCl signaling (#22) and the negative effect of H_2_O_2_ on NO/peroxynitrite signaling (#23).

This study confirms that the HOCl and the NO/peroxynitrite signaling pathways are mutually exclusive. Exclusivity and mutual negative interference of the two pathways become apparent when exogenous NO is added to transformed cells with ongoing HOCl signaling. The added NO first completely inactivates HOCl signaling before it establishes NO/peroxynitrite signaling (see Figure 4). The mutual exclusivity of the two signaling pathways is also apparent when gradual inhibition of membrane-associated catalase of MKN-45 gastric carcinoma cells first reactivates NO/peroxynitrite signaling selectively and then, with increasing concentrations of free H_2_O_2_, allows for HOCl signaling, with parallel suppression of NO/peroxynitrite signaling (Figure 6). The data from this study allow to establish a model for this mutual exclusivity, as discussed below.

H_2_O_2_ is the central protagonist for HOCl signaling, as it is the substrate for DUOX- peroxidase-dependent HOCl synthesis (Figure 18 A, reaction #2). However, excess H_2_O_2_ causes an antagonistic effect on HOCl signaling, as shown by the negative effect of part C on part A in Figure 18. Previous studies have shown that this antagonistic effect is mainly due to the direct reaction between HOCl and H_2_O_2_ [10, 27]. This important modulating step is confirmed in this study and is shown to be independent of iron.

The chemical biology of NO/peroxynitrite action is determined by several factors and particularly by site-specific effects. NO/peroxynitrite signaling is counteracted by oxidation of NO by molecular oxygen [11, 82] (not shown in Figure 18). In close vicinity to the cell membrane with its proton pumps, NO/peroxynitrite signaling according to Figure 18 B is favoured, leading to the centrally important peroxynitrous acid, which generates hydroxyl radicals and NO_2_ [13]. More distant of the cell membrane, the more efficient reaction between peroxynitrite and CO_2_ (Figure 18 D) [19, 20] most likely prevents the formation of peroxynitrous acid and in this way reduces the efficiency of NO/peroxynitrite signaling.

Based on its established reaction potential [50, 51], NO might theoretically interfere with lipid peroxidation by hydroxyl radicals through reactions #15 and #16 and/or reactions #17 and #18, whereas H_2_O_2_ might scavenge hydroxyl radicals through reactions #19 and #20 [83, 84]. However, based on our data, none of these reactions seems to have a recognizable effect on apoptosis induction through HOCl or NO/peroxynitrite signaling under the conditions used for our cell systems. Otherwise an increase in NO i) should not allow enhancement of NO/peroxynitrite signaling and ii) should not allow to rescue HOCl signaling from supraoptimal inhibition at high concentrations of H_2_O_2_ (reaction #22).

The experimental system of apoptosis induction by chemically synthesized exogenous peroxynitrite (Figure 14) seems to be an exception. The special situation under this particular experimental system seems to allow an interaction between tumor cell-derived H_2_O_2_ and exogenous peroxynitrite, in line with the findings established by Di Mascio [78] and further extended in references [43, 47].

The central new themes of this study are i) the inhibition of HOCl signaling by NO (reaction # 21 in Figure 18), ii) the interference of NO with H_2_O_2_-mediated inhibition of HOCl signaling, leading to an enhancement of HOCl signaling (reaction #22), and the inhibition of NO/peroxynitrite signaling by H_2_O_2_ (reaction # 23). It is shown that these three activities are biochemically interconnected. This allows to explain the paradoxically seeming nature of these effects.

The inhibitory effect of exogenous or cell-derived NO on HOCl signaling cannot be explained by i) inhibition of peroxidase by NO [85], ii) scavenging of HOCl by NO, iii) scavenging of superoxide anions by NO, iv) scavenging of hydroxyl radicals by NO or v) inhibition of lipid peroxidation, as otherwise the rescue of HOCl signaling from the inhibitory effects of excess H_2_O_2_ through an increase in the NO concentration would not be feasible. As deduced from the inhibition profile, this rescue is based on establishment of functional HOCl signaling and therefore requires peroxidase, HOCl, superoxide anions, hydroxyl radicals and subsequent lipid peroxidation. However, a concentration-dependent consumption of H_2_O_2_ by NO could well explain the inhibition of HOCl signaling (by removal of the POD substrate H_2_O_2_ and thus preventing HOCl synthesis), as well as the abrogation of H_2_O_2_-dependent inhibition of HOCl signaling (through removal of excess H_2_O_2_ and thus bringing HOCl synthesis from supraoptimal inhibiton back to optimal values). The consumption of H_2_O_2_ by NO is in good agreement with the effects of the catalase mimetic EUK-134 that are analogous to those of NO, as EUK-134 inhibits HOCl signaling, but also can bring supraoptimal inhibition of HOCl signaling through excess H_2_O_2_ back to its optimum, provided the concentration of EUK-134 is well-balanced. As no direct consumption reaction between H_2_O_2_ and NO is known, this interaction seems to be based on indirect effects. These will be elucidated here.

As the effects of NO on HOCl signaling were observed when exogenous NO donors were applied or when the cell-derived NO concentration was increased i) through increasing the NOS substrate arginine, ii) by increasing the expression of NOS through interferon, and iii) by increasing the available concentration of NO through inhibiton of NO dioxygenase by various inhibitors, the central role of NO in this process is clearly established.

Furthermore, it was assured that both effects of NO on HOCl signaling are based on NO, but not on its reaction product peroxynitrite, as the peroxynitrite decomposition catalyst FeTPPS did not interfere with either of these NO effects, whereas prevention of NO generation did, whenever cell-derived NO rather than NO released by a NO donor was utilized in the experiments.

The reverse effect, i. e. H_2_O_2_-mediated inhibition of NO/peroxynitrite signaling was achieved through i) increasing the concentration of free H_2_O_2_ by increasing the cell density of NOX1 expressing malignant cells, ii) addition of H_2_O_2_-generating glucose oxidase, iii) increasing the inhibition of membrane-associated catalase of tumor cells or iv) enhancing NOX-1 activity through low dose gamma irradiation, and thus cause a net increase in free H_2_O_2_ derived from dismutation of NOX1-derived superoxide anions. In all of these regimes, a negative effect of H_2_O_2_ on NO/peroxynitrite signaling was observed. The dependency on the concentration of the NO donor as determined in Figure 13 B indicates a titration effect of H_2_O_2_ on the concentration of NO that is finally entering NO/peroxynitrite signaling, as an increase in the effectiveness of the NO donor was starting at a higher concentration of the NO donor when a higher concentration of free H_2_O_2_ was reached through application of the higher concentration of the catalase inhibitor. In a complementary approach, the reduction of free H_2_O_2_ through the addition of soluble catalase caused an increase in NO/peroxynitrite signaling, which is in line with the preceding results. The enhancing effects of catalase were only detectable in the concentration range of catalase that did not interfere with NO/peroxynitrite signaling per se. This part of the results confirms the multiple reaction potentials of catalase that may target its classical substrate H_2_O_2_, but also NO or peroxynitrite.

The central modulatory effects addressed in this study i. e. i) the inhibition of HOCl signaling by NO, ii) the interference of NO with H_2_O_2_-mediated inhibition of HOCl signaling, leading to reactivation of HOCl signaling, and iii) the inhibition of NO/peroxynitrite signaling by H_2_O_2_ cannot be explained by known reactions based on the classical chemical biology of NO/peroxynitrite and H_2_O_2_/HOCl. The finding that each one of these three modulatory effects was strongly abrogated after removal of free iron through deferoxamine was the clue for understanding these effects. Iron-mediated reactions therefore seem to be the central point for the understanding of the underlying mechanism(s) and seem to be involved in one or more central steps during these complex reactions. The inclusion of established reaction potentials of iron in Figure 19 allows to propose a scheme for mutual consumption of NO and H_2_O_2_, in line with our experimental findings. One central element in the interactions studied here is the reaction between H_2_O_2_ and ferrous ion, leading to the classical Fenton reaction (reaction #12 in Figure 19 A) [76, 77]

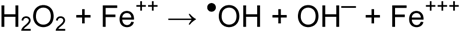

As H_2_O_2_ is a far-ranging ROS, most of these hydroxyl radicals will not reach the cell membrane and therefore do not cause damage. However, ^●^OH has a chance to react with ^●^NO (reaction #13) [50, 51].

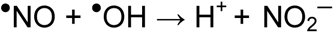

Ferric iron, resulting from the Fenton reaction can also interact with NO (reaction #14) [71–75], leading to free ferrous iron through reaction #15.

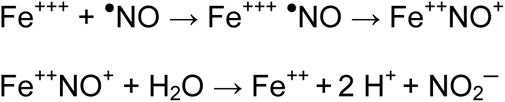

Free ferrous iron then may contribute to the next cycle of Fenton reaction. This cyclic reaction may lead to the consumption of maximally two molecules of NO, minimally one molecule of NO, after Fenton chemistry of one molecule of H_2_O_2_. As long as both compounds (H_2_O_2_ and NO) are present, this cycle may continue, as ferrous iron is constantly recycled in this process. The compound with the higher concentration, i. e. H_2_O_2_ or NO, will remain at a certain concentration after cycling is completed. Therefore, high concentrations of NO might consume H_2_O_2_ and thus prevent HOCl synthesis (reaction # 3), or consume excess H_2_O_2_ (reaction #18) and in this way restore optimal HOCl signaling through prevention of H_2_O_2_-mediated inhibition of HOCl signaling (reaction #6). In contrast, high concentrations of H_2_O_2_ may lead to consumption of NO through reactions #13 and 15 and thus prevent the formation of peroxynitrite (reaction #8) and its apoptosis-inducing consequences (reactions #10, 11).

**Figure 19.**
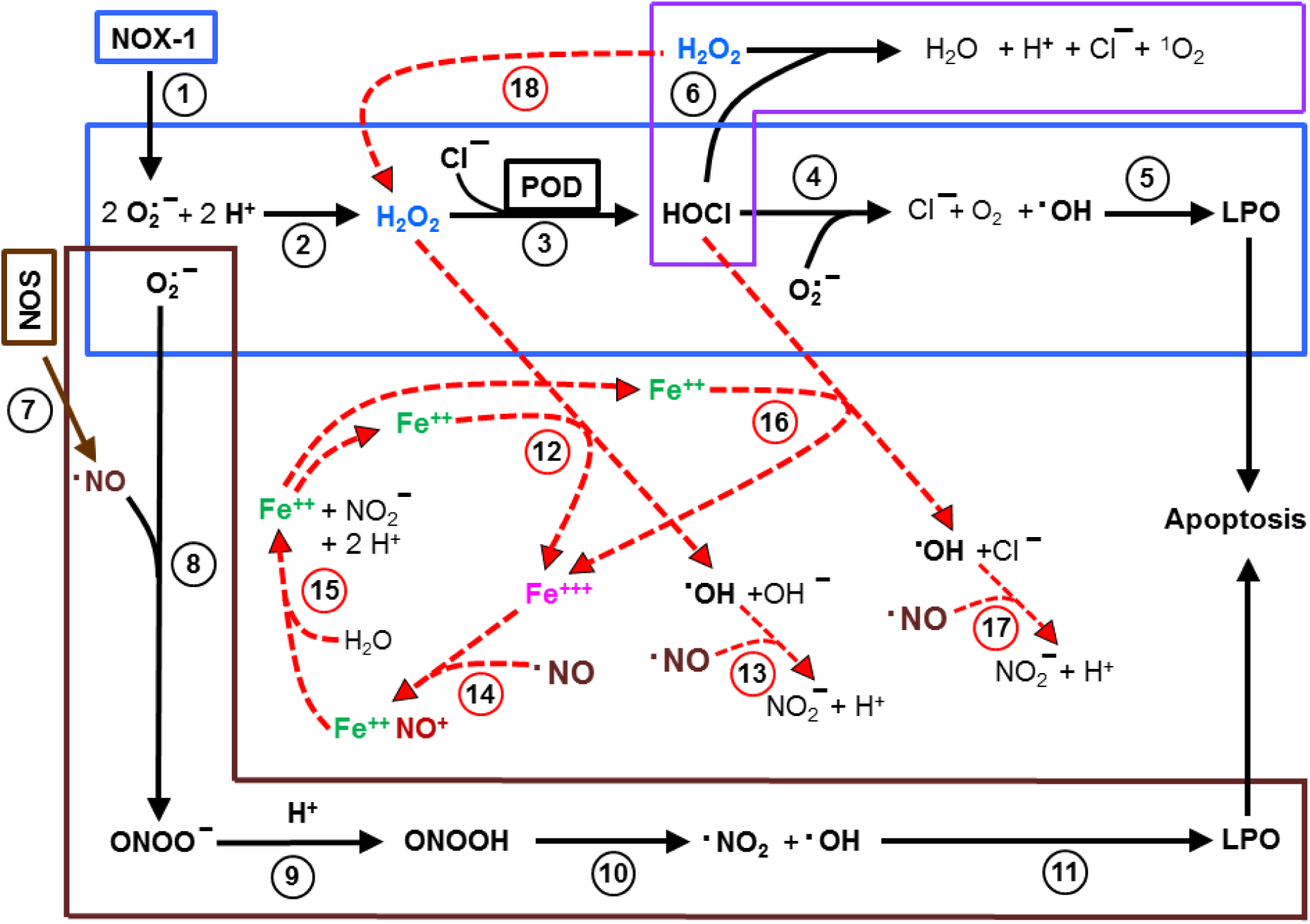
Iron-catalyzed mutual consumption of NO and H_2_O_2_/HOCl. NOX-1-derived superoxide anions (#1) drive HOCl signaling (#2 -5) which leads to lipid peroxidation (LPO) and apoptosis induction. Excess H_2_O_2_ leads to the consumption of HOCl and thus blunts HOCl signaling (#6). NOS-derived NO (#7) and NOX-1-derived superoxide establish NO/peroxynitrite signaling (#8-11). Ferrous iron (Fe^++^) and H_2_O_2_ establish Fenton chemistry (#12). The resultant hydroxyl radicals may react with NO (#13). Ferric iron (Fe^+++^) resulting from Fenton reaction reacts with NO (#14). After reaction of the resulting Fe^++^NO^+^ complex with water (#15), ferrous iron is regenerated and ready for the next round of action. Alternatively, ferrous iron may react with HOCl (#16), followed by the reaction of hydroxyl radicals with NO (#17) and recycling ferric iron back to ferrous iron through reaction steps #14, 15. These interactions describe the mechanisms underlying the inhibition of HOCl signaling by NO, the interference of NO with H_2_O_2_-mediated inhibition of HOCl signaling and the interference of H_2_O_2_ with NO/peroxynitrite signaling. They thus explain the mutual exclusivity of NO/peroxynitrite and HOCl signaling.

This basic cycle of mutual iron-driven consumption of H_2_O_2_ and NO seems to extend to a more complex situation, as soon as peroxidase-dependent HOCl synthesis becomes effective. In the presence of optimal concentrations of peroxidase and appropriate concentrations of H_2_O_2_, peroxidase will efficiently use H_2_O_2_ as substrate for HOCl synthesis. The utilization of H_2_O_2_ by peroxidase will drastically reduce H_2_O_2_ - based Fenton chemistry (reaction #12). However, as HOCl has a thousandfold higher efficiency to undergo Fenton chemistry than H_2_O_2_ [86], the cycle of NO consumption should be easily maintained, driven by HOCl/iron reaction (reaction #16).

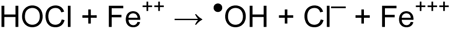

With an excess of NO in the system, the concentration of HOCl may thus be completely diminished (following the reaction sequence #16, 17, 14, 15, 16). This will lead to inhibition of classical HOCl signaling (reactions #4,5). It therefore can be concluded that under conditions of optimal HOCl synthesis, the negative effect of NO on HOCl signaling will be primarily through consumption of HOCl rather than prevention of HOCl synthesis through consumption of the peroxidase substrate H_2_O_2_, though both processes lead to the same biological effect. The role of HOCl in the iron-mediated scenario described in Figure 19 is strongly supported by the findings that removal of HOCl by taurine can substantially enhance NO/peroxynitrite signaling, as shown in Figures 4, 6, and 9. The analogous effect of the catalase mimetic EUK-134 under these experimental conditions might therefore be an indirect one– acting through prevention of HOCl synthesis through decomposition of H_2_O_2_.

Under conditions of relative excess of H_2_O_2_ that interferes with HOCl signaling, H_2_O_2_ may again become the target for consumption by NO again, as the concentration of free HOCl is strongly reduced through its reaction with H_2_O_2_ (reaction #6). Under these conditions, iron-mediated consumption of H_2_O_2_ by NO is more likely than iron-mediated consumption of HOCl. If sufficient concentrations of NO are supplied, the consumption of H_2_O_2_ by NO will gradually lead to abrogation of H_2_O_2_ -mediated inhibition of HOCl signaling and allow for HOCl/superoxide anion interaction. With further availability of free NO, this pathway can then be predicted to be preferentially inhibited by NO through consumption of HOCl.

Figure 19 defines all the conditions that seem to be relevant for our central findings of NO-mediated inhibition of HOCl signaling, NO-mediated abrogation of H_2_O_2_- mediated inhibition of HOCl signaling and H_2_O_2_-mediated inhibition of NO/peroxynitrite signaling. Delicate interactions define the dominant pathway and thus explain the mutual exclusivity of these two major signaling pathways.

Finally, this study confirms for a large number of bona fide human tumor cells that HOCl- as well as NO/peroxynitrite signaling are under stringent control by membrane-associated catalase. Therefore, inhibition, inactivation or genetic knockdown of catalase has been suggested as selective, redox-driven method for tumor therapy [6, 7, 10, 13, 28, 30, 39, 40–45]. The data on the mutual interactions between the two signaling pathways allows to understand the mechanisms underlying the preferences of tumor cells for either HOCl or NO/peroxynitrite signaling after inhibition or inactivation of catalase. These data also confirm that the “target cell function” of malignant cells, i.e. its ability to allow and drived apoptosis-inducing signaling is defined by active NOX in combination with inactive NOX, whereas the “effector function”, the potential to contribute essential signaling components like NO or the peroxidase domain of DUOX is independent of the malignant state and is determined by the concentration of these components. Taken together, this information might be helpful to determine conditions that enhance ROS/RNS-driven tumor cell apoptosis and prevent potential interfering mechanisms during tumor therapy. It is important to keep in mind, that this complex control system of redox-controlled tumor cell apoptosis is only one part within the context of tumor cell control. ROS/RNS-dependent apoptosis induction in tumor cells is dependent on abrogation of intracellular counteracting processes that interfere with lipid peroxidation. This dominant regulatory system is controlled by the concentration of intracellular glutathione and glutathione peroxidase, driven by the activity of glutathione-synthesizing enzymes and the xC transporter [18]. It needs also be kept in mind, that successful tumor treatment not only requires specific induction of tumor cell apoptosis, but is also essentially dependent on the induction of selective immunological processes, activated by specific signals of dying tumor cells [30]. These different levels of tumor control seem to be connected to a significant extend by redox biology with its many facettes of molecular interactions, elucidated by a fascinating scientific process that started 51 years ago with the demonstration of H_2_O_2_ in human tissue [88–90].

## Supporting information

BauerNOH2O2.pdf

## Acknowledgements

The professional finalization of the graphs by Jürgen Brandel (Freiburg) and the financial support by the Medical Faculty of the University of Freiburg for publication gratefully costs is acknowledged.

## References

1. Irani K, Xia Y, Zweier JL, Sollott SJ, Der CJ, Fearon ER, Sundaresan M, Finkel T, Goldschmidt-Clermont PJ: Mitogenic signalling by oxidants in Ras-transformed fibroblasts. Science 275 (1997) 1649–1652.

2. Irani K, Goldschmidt-Clermont PJ: Ras, Superoxide and Signal Transduction. Biochem. Pharmacol. 55 (1998) 1339–1346.

3. Lopez-Lazaro M: Excessive superoxide anion generation plays a key role in carcinogenesis. Int. J. Cancer 120 (2007) 1378–1380.

4. Suh Y-A, Arnold RS, Lassegue B, Shi J, Xu X, Sorescu D, Chung AB, Griendling KK and Lambeth JD: Cell transformation by the superoxide-generating oxidase Mox1. Nature 401: 79–82, 1999.

5. Yang JQ, Li S, Domann FE, Buettner G and Oberley LW: Superoxide generation in *v-Ha-ras*-transduced human keratinocyte HaCaT cells. Mol Carcinogenesis 26: 180–188, 1999.

6. Bauer G. Tumor cell protective catalase as a novel target for rational therapeutic approaches based on specific intercellular ROS signaling. Anticancer Res. 32 (2012) 2599–2624.

7. Bauer G. Targeting extracellular ROS signaling of tumor cells. Anticancer Res. 34 (2014) 1467–1482.

8. Herdener M, Heigold S, Saran M and Bauer G: Target cell-derived superoxide anions cause efficiency and selectivity of intercellular induction of apoptosis. Free Rad Biol Med 29: 1260–1271, 2000.

9. Heigold S, Sers C, Bechtel W, Ivanovas B, Schäfer R, Bauer G. Nitric oxide mediates apoptosis induction selectively in transformed fibroblasts compared to nontransformed fibroblasts. Carcinogenesis 23 (2002) 929–941.

10. Bauer G. HOCl and the control of oncogenesis. J. Inorganic. Biochem. 179 (2018 a) 10–23. doi:10.1016/j.jinorgbio.2017.11.005

11. Bauer G. Nitric oxide contributes to selective apoptosis induction in malignant cells through multiple reaction steps. Critical Reviews in Oncogenesis 21 (2016) 365–398, 2016

12. Bauer G, Chatgilialoglu C, Gebicki JL, Gebicka L, Gescheidt G, Golding BT, Goldstein S, Kaizer J, Merenyi G, Speier G and Wardman P: Biologically relevant small radicals. Chimia 62: 1–9, 2008.

13. Bauer G. Increasing the endogenous NO level causes catalase inactivation and reactivation of intercellular apoptosis signaling specifically in tumor cells. Redox Biol. 6 (2015) 353–371.

14. Saran M and Bors W: Oxygen radicals as chemical messengers: a hypothesis. Free Rad Res Comm 7: 213–220, 1989.

15. Saran M and Bors W: Signalling by O2- and NO: How far can either radical, or any specific reaction product, transmit a message under *in vivo* conditions? Chem Biol Interact 90: 35–45, 1994.

16. Bauer G. Central signaling elements of intercellular reactive oxygen/nitrogen species-dependent induction of apoptosis in malignant cells. Anticancer Res 37 (2017 a) 499–514.

17. Kundrát P, Bauer G, Jacob P and Friedland W. Mechanistic modelling suggests that the size of preneoplastic lesions is limited by intercellular induction of apoptosis in oncogenically transformed cells. Carcinogenesis 33: 253–259, 2012.

18. Bauer G. Inhibition of membrane-associated catalase, extracellular ROS/RNS signaling and aquaporin/H_2_O_2_-mediated intracellular glutathione depletion cooperate during apoptosis induction in the human gastric carcinoma cell line MKN-45. Antioxidants 2021, 10, 1585. https://doi.org/10.3390/antiox10101585

19. Denicola ABA, Freeman BA, Trujillo M, Radi R. Peroxynitrite reaction with carbon dioxide/bicarbonate: kinetics and influence on peroxynitrite-mediated reactions. Arch. Biochem. Biophys. 333 (1996) 49–58.

20. Goldstein S, Czapski G. Formation of peroxynitrate from the reaction of peroxynitrite with CO_2_: Evidence for carbonate radical production. J. Am. Chem. Soc. 120 (1998) 3458–3463.

21. Deichman GI and Vendrov EL: Characteristics of in vitro transformed cells essential for their in vivo survival, selection and metastatic activity. Int. J. Caner 37: 401–409, 1986.

22. Deichman GI, Kluchareva TE, Matveeva VA, Kushlinsky NE, Bassalyk LS and Vendrov EL: Clustering of discrete cell properties essential for tumorigenicity and metastasis. I. Studies of syrian hamster embryo fibroblasts spontaneously transformed in vitro. Int. J. Cancer 44: 904–907, 1989.

23. Deichman G, Matveeva VA, Kashkina LM, Dyakova NA, Uvarova EN, Nikiforov MA and Gudkov AV: Cell transforming genes and tumor progression: *in vivo* unified secondary phenotypic cell changes. Int J Cancer 75: 277–283, 1998.

24. Deichman G: Natural selection and early changes of phenotype of tumor cells in vivo: Acquisition of new defense mechanisms. Biochem 65: 78–94, 2000.

25. Deichman G: Early phenotypic changes of in vitro transformed cells during in vivo progression: possible role of the host innate immunity. Sem Cancer Biol 12: 317–326, 2002.

26. Bechtel W and Bauer G: Catalase protects tumor cells against apoptosis induction by intercellular ROS signaling. Anticancer Res 29: 4541–4557, 2009

27. Bechtel W and Bauer G: Modulation of intercellular ROS signaling of human tumor cells. Anticancer Res 29: 4559–4570, 2009

28. Heinzelmann S and Bauer G. Multiple protective functions of catalase against intercellular apoptosis-inducing ROS signaling of human tumor cells. Biol. Chem. 391, 675–693, 2010.

29. Böhm B, Heinzelmann S, Motz M, Bauer G. Extracellular localization of catalase is associated with the transformed state of malignant cells. Biol. Chem. 396 (2015) 1339–1356.

30. Bauer G. Signal amplification by tumor cells: clue to the understanding of the antitumor effects of cold atmospheric plasma and plasma-activated medium. IEEE Transactions on Radiation and Plasma Medical Sciences 2 (2018b) 87–98

31. Gebicka L, Didik J. Catalytic scavenging of peroxynitrite by catalase. Int. J. Inorg. Biochem. 103 (2009) 1375–1379.

32. Brunelli L, Yermilov V and Beckman JS. Modulation of catalase peroxidatic and catalytic activity by nitric oxide. Free Rad.Biol Med 30: 709–714, 2001.

33. Wink DA, Mitchell JB. Chemical biology of nitric oxide: insights into regulatory, cytotoxic, and cytoprotective mechanisms of nitric oxide. Free Radic Biol & Medicine 1998;25: 434–456.

34. Bauer G. SiRNA-based analysis of the abrogation of the protective function of membrane-associated catalase of tumor cells. Anticancer Res 37 (2017b) 567–582.

35. Bauer G. HOCl-dependent singlet oxygen and hydroxyl radical generation modulate and induce apoptosis of malignant cells. Anticancer Res. 33 (2013) 3589–3602.

36. Y. Kono and I. Fridovich, “Superoxide radical inhibits catalase”, J Biol Chem, vol. 257, pp- 5751–5754, 1982.

37. N. Shimizu, K. Kobayashi and K. Hayashi, “The reaction of superoxide radical with catalase. Mechanism of the inhibition of catalase by superoxide radical”, J Biol Chem, vol. 259, pp. 4414–4418, 1984.

38. L. Gebicka, D. Metodiewa and J. L. Gebicki, “Pulse radiolysis of catalase in solution. I. Reactions of O ^.-^ with catalase and its compound I”, Int J Rad Biol, vol. 55, pp. 45–50, 1989.

39. Riethmüller M, Burger N, Bauer G: Singlet oxygen treatment of tumor cells triggers extracellular singlet oxygen generation, catalase inactivation and reactivation of intercellular apoptosis-inducing signaling. Redox Biol, 6 (2015) 157–168.

40. Scheit K and Bauer G. Direct and indirect inactivation of tumor cell protective catalase by salicylic acid and anthocyanidins reactivates intercellular ROS signaling and allows for synergistic effects. Carcinogenesis 36 (2015) 400–411.

41. Bauer G, Motz M: The antitumor effect of single-domain antibodies directed towards membrane-associated catalase and superoxide dismutase. Anticancer Res. 36 (2016) 5945–5956.

42. Bauer G, Graves DB. Mechanisms of selective antitumor action of cold atmospheric plasma-derived reactive oxygen and nitrogen species. Plasma Process. Polymer. 13 (2016) 1157–1178.

43. Bauer G. The synergistic effect between hydrogen peroxide and nitrite, two long-lived molecular species from cold atmospheric plasma, triggers tumor cells to induce their own cell death. Redox Biol, 26, 101291 (2019). https://doi.org/10.1016/j.redox.2019.101291

44. Bauer G. Intercellular singlet oxygen-mediated bystander signaling triggered by long-lived species of cold atmospheric plasma and plasma-activated medium. Redox Biol 26, 101301, 2019. https://doi.org/10.1016/j.redox.2019.101301

45. Bauer G, Sersenova D, Graves DB and Machala Z. Cold atmospheric plasma and plasma-activated medium trigger RONS-based tumor cell apoptosis. Scientific Reports 9, 14210, 2019. https://doi.org/10.1038/s41598-019-50291-0

46. Bauer G, Sersenova D, Graves DB and Machala Z. Dynamics of singlet oxygen-triggered, RONS-based apoptosis induction after treatment of tumor cells with cold atmospheric plasma or plasma-activated medium. Scientific Reports 9: 1393, 2019. https://doi.org/10.1038/s41598-019-50329-3

47. Bauer G. Autoamplificatory singlet oxygen generation sensitizes tumor cells for intercellular apoptosis-inducing signaling. Mech. Ageing Develop. 172 (2018d) 59–77.

48. Wink DA, Hanbauer I, Krishna MC, Degraff W, Gamson J, Mitchell JB. Nitric oxide protects against cellular damage and cytotoxicity from reactive oxygen species. PNAS 90: 9813–9817, 1993

49. Pacelli R, Wink DA, Cork JA, Krishna MC, DeGraff W, Friedman N, Tsokos M, Samuni A, Mitchell JB. Nitric oxide potentiates hydrogen peroxide-induced killing of Escherichia coli. J Exp Med 182:; 1469–1479, 1995.

50. Dalloz F, Maupoil V, Lecour S, Briot F, and Rochette L. In vitro studies of interactions of nitric oxide donor drugs with superoxide and hydroxyl radicals. Mol Cell Biochem 177: 193–200, 1997.

51. Xu KY. Nitric oxide protects nitric oxide synthase function from hydroxyl radical-induced inhibition. Biochem Biophys Acta 1481: 156–166, 2000.

52. Wink DA and Ford PC: Nitric oxide reactions important to biological systems: A survey of some kinetics investigations. Methods: A companion to Methods in Enzymology 7: 14–20, 1995

53. Connick RE: The interaction of hydrogen peroxide and hypochlorous acid in acidic solutions containing chloride ion. J Am Chem Soc 69: 1509–1514, 1947.

54. Held AM, Halko DJ and Hurst JK: Mechanisms of chlorine oxidation of hydrogen peroxide. J. Am. Chem. Soc. 100: 5732–5740, 1978

55. Haberstroh K, Heigold S and Bauer G. Transformed cell-derived reactive oxygen species support and inhibit nitric oxide-mediated apoptosis induction. Int J Oncol. 21:145–151, 2002.

56. Jürgensmeier J, Schmitt CP, Viesel E, Höfler P and Bauer G: TGF-ß-treated normal fibroblasts eliminate transformed fibroblasts by induction of apoptosis. Cancer Res 54: 393–398, 1994.

57. Bauer G, Bereswill S, Aichele P and Glocker E. Helicobacter pylori protects oncogenically transformed cells from reactive oxygen species-mediated intercellular induction of apoptosis. Carcinogenesis 35 (2014) 1582–1591.

58. Kerr, JFR, Wyllie AH, and Currie AR. Apoptosis: a basic biological phenomenon with wide-ranging implications in tissue kinetics. Br J Cancer 26: 239–257, 1972.

59. Elmore S. Apoptosis: A review of programmed cell death. Toxicol Pathol 35: 495–515, 2007.

60. Beck E, Schäfer R and Bauer G: Sensitivity of transformed fibroblasts for intercellular induction of apoptosis is determined by their transformed phenotype. Exp Cell Res 234: 47–56, 1997.

61. Pottgiesser S, Heinzelmann S and Bauer G. Intercellular HOCl-mediated apoptosis induction in malignant cells: interplay between NOX1-dependent superoxide anion generation and DUOX-related HOCl-generating peroxidase activity. Anticancer Res. 35: 5927–5943, 2015.

62. Gardner PR, Martin LA, Hall D and Gardner AM: Dioxygen-dependent metabolism of nitric oxide in mammalian cells. Free Rad Biol Med 31: 191–204, 2001

63. Hallstrom CK, Gardner AM and Gardner PR: Nitric oxide metabolism in mammalian cells: substrate and inhibitor profiles of a NADPH-cytochrome P450 oxidoreductase-coupled microsomal nitric oxide dioxygenase. Free Rad Biol Med 37: 216–228, 2004.

64. Gardner PR: Assay and characterization of the NO dioxygenase activity of flavohemoglobins. In: Globins and Other Nitric Oxide-reactive Proteins. Meth Enzymol 436: 217–237, 2008.

65. Schmidt K and Mayer B. Consumption of nitric oxide by endothelial cells: Evidence for the involvement of a NAD(P)H-, flavin and heme-dependent dioxygenase reaction. FEBS Lett 577: 199–204, 2004.

66. Karupiah G, Cie Q-W, Buller RML, Nathan C, Duarte C and MacMicking JD: Inhibition of viral replication by interferon-γ-induced nitric oxide synthase. Science 261: 1445–1448, 1993.

67. Santiskulvong C and Rozengurt E: Galardin (GM 6001), a broad-spectrum matrix metalloproteinase inhibitor, blocks bombesin- and LPA-induced EGF receptor transactivation and DNA synthesis in rat-1 cells. Exp Cell Res 290: 437–446, 2003.

68. Wink DA and Ford PC: Nitric oxide reactions important to biological systems: A survey of some kinetics investigations. Methods: A companion to Methods in Enzymology 7: 14–20, 1995

69. Wink DA and Mitchell JB: Chemical biology of nitric oxide: insights into regulatory, cytotoxic, and cytoprotective mechanisms of nitric oxide. Free Radic Biol & Medicine 25: 434–456, 1998 a)

70. Wink DA and Mitchell JB: Nitric oxide and cancer: An introduction. Free Radic Biol & Medicine 34: 951–954, 2003

71. Farias-Eisner R, Chaudhuri G, Aeberhard E, Fukuto JM. The chemistry and tumoricidal activity of nitric oxide/hydrogen peroxide and the implications to cell resistance/susceptibility. J Biol Chem 271: 6144–5151, 1996.

72. Goretski J, Hollocher TC (1991) Catalysis of nitrosyl transfer by denitrifying bacteria is facilitated by nitric oxide. Biochem Biophys Res Commun 175: 901–905

73. Stamler JS, Singel DJ, Loscalzo J (1992) Biochemistry of nitric oxide and its redox-activated forms. Science 258: 1898–1902

74. Graziano M, Beligni MV, Lamattina L. Nitric Oxide Improves Internal Iron Availability in Plants. Plant Physiol 130: 1852–1859, 2002

75. Graziano M and Lamattina L: Nitric oxide and iron in plants: an emerging and converging story. Trends in Plant Science 10: 4–8, 2005.

76. Winterbourn C: Toxicity of iron and hydrogen peroxide: the Fenton reaction. Toxicol Letters 82/83: 969–974, 1995

77. Neyens E and Baeyens J: A review of classic Fenton s peroxidation as an advanced oxidation technique. Journal of Hazardous Materials B98: 35–50, 2003

78. Di Mascio P, Bechara EJH, Medeiros MHG, Briviba K and Sies H: Singlet molecular oxygen production in the reaction of peroxynitrite with hydrogen peroxide. FEBS Lett 355: 287–289, 1994.

79. Bauer G. Low dose irradiation enhances specific signaling components of intercellular reactive oxygen-mediated apoptosis induction. J. Phys. Conf. ser. 261 012001, 2011

80. Temme J and Bauer G: Low-dose gamma irradiation enhances superoxide anion production by nonirradiated cells through TGF-β1-dependent bystander signaling. Rad. Res. 179: 422–432, 2013.

81. Abdelrazzak AB, Pottgiesser SJ, Hill MA, ÒNeCill P and Bauer G. Enhancement of peroxidase release from non-malignant and malignant cells through low-dose irradiation with different radiation quality. Radiation Res 185: 199–213, 2016.

82. Goldstein S and Czapski G. Kinetics of nitric oxide autooxidation in aqueous solution in the absence and presence of various reductants. The nature of the oxidizing intermediates. J Am Chem Soc 117: 12078–12084, 1995 a

83. Christensen H, Sehested K, Corfitzen H. Reactions of hydroxyl radicals with hydrogen peroxide at ambient and elevated temperature. J. Phys. Chem. 86 (1982) 1588–1590.

84. Buxton, G.V.; Greenstock, C.L.; Helman, W.P.; Ross, A.B. Critical reviews of rate constants for reactions of hydrated electrons, hydrogen atoms and hydroxyl radicals (.OH/.O^−^) in aqueous solution. J. Phys. Chem. Ref. Data, 1988, 17, 513–886.

85. Abu-Soud HM and Hazen SL. Nitric oxide modulates the catalytic activity of myeloperoxidase. J Biol Chem 275: 5425–5430, 2000 a.

86. Folkes LK, Candeias LP, Wardman P. Kinetics and mechanisms of hypochlorous acid reactions. Arch. Biochem. Biophys. 323 (1995) 120–126.

87. Scheit K and Bauer G. Synergistic effects between catalase inhibitors and modulators of nitric oxide metabolism on tumor cell apoptosis. Anticancer Res 34: 5337–5350, 2014.

88. Sies H, Chance B. The steady state level of catalase compound I in isolated hemoglobin-free perfused rat liver. FEBS Letters 1970;11:172–176.

89. Sies H, Jones DP. Reactive oxygen species (ROS) as pleiotropic physiological signaling agents. Nature Reviews Mol. Cell. Biol. 2020; 21: 363–383.

90. Sies H. Findings in redox biology: From H_2_O_2_ to oxidative stress. J Biol Chem 2020; 295:13458–13473

