## Supplementary material for "Iron-catalyzed antagonism between NO and H_2_O_2_ / HOCl causes mutual exclusivity of HOCl- and NO/peroxynitrite-mediated apoptosis-inducing signaling": BauerNOH2O2.pdf

Hydrogen peroxide and nitric oxide: protagonists and antagonists of reactive oxygen species/reactive nitrogen species (ROS/RNS)-mediated intercellular apoptosis-inducing signaling

Georg Bauer


**Contents of Supplementary Material:**

**Supplementary Results**

*1. Additional complexity of interaction between the NO donor SNP and HOCl signaling*
*2. Interaction of SNP-derived NO with HOCl signaling of the murine fibrosarcoma cell line L929*
*3. Three distinct effects of SNP-derived NO on apoptosis-inducing signaling in MKN-45 cells depend on the concentration of the NO donor*
*4. Inhibition of HOCl signaling by the NOD inhibitor quercetin and its derivatives quercitrin, isoquercitrin and rutin.*
*5. Dual role of superoxide anions for NO/peroxynitrite signaling*
6. *The potential role of ferrous/ferric ions for the antagonistic action between hydrogen peroxide and NO*
*7. Tumor cells from different tissues show differences in the quality of their apoptosis-inducing signaling chemistry*
*8. The determining role of the effector function for the quality of intercellular ROS/RNS-dependent apoptosis-inducing signaling. The use of transformed target cells.*
*9. Intercellular induction of apoptosis in bona fide tumor cells.Inhibition of protective catalase is necessary to activate the target function, whereas the effector function determines the quality of signaling.*
*10. The effect of exogenous NO on the quality of intercellular ROS/RNS signaling of tumor cells with different signaling qualities.*

**Supplementary Materials and Methods**

**Supplementary References**

**Supplementary Results**

*1. Additional complexity of interaction between the NO donor SNP and HOCl signaling*

The analysis shown in Figure 4 in the main document led to the unexpected result that the peroxynitrite decomposition catalyst FeTPPS interfered not only with NO/peroxynitrite signaling, but also resulted in a leftward shift of the curve that described HOCl-mediated apoptosis induction in the presence of increasing concentrations of the NO donor SNP. Finally, complete inhibition of apoptosis was reached. On the first sight, this might be misinterpreted as a direct effect of FeTPPS on HOCl signaling. This particular effect of FeTPPS was repeated in a follow-up experiment (Supplementary Figure S1). The application of the singlet oxygen scavenger histidine and of discrete, low concentrations of the NOX1 inhibitor AEBSF and the catalase mimetic EUK-134 also caused a leftward shift of the curve. In the case of these scavengers, apoptosis induction increased again after the point of maximal inhibition. In line with previous findings [1-3], the following explanation for the observed effects is suggested: NO derived from SNP not only seems to interfere with HOCl signaling through consumption of $H_2O_2$, but also might allow for peroxynitrite formation. As a result, the complex interaction between $H_2O_2$ and peroxynitrite [3-5] then may generate singlet oxygen. As transformed cells carry a distinct concentration of catalase on their surface – not sufficient to inhibit HOCl signaling, but sufficient to modulate HOCl signaling [6], the generation of singlet oxygen under the experimental conditions can lead to inactivation of catalase [7-9]. Subsequently, the concentration of free $H_2O_2$ is enhanced and leads to increased consumption of NO. As a result, higher concentrations of the NO donor are required

Supplementary Figure S1

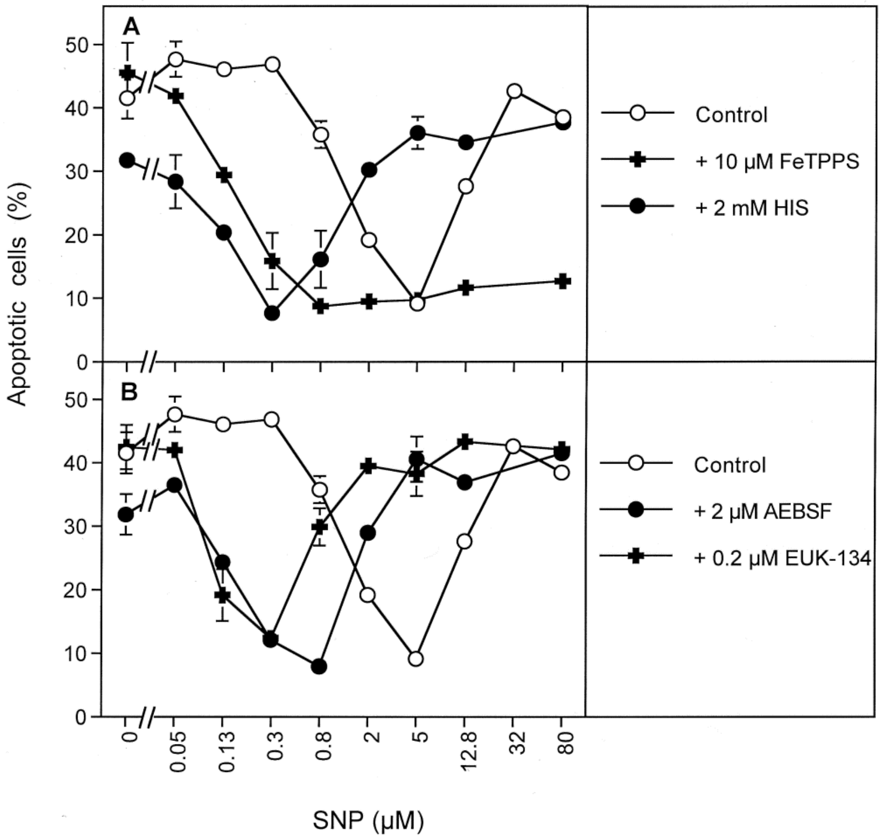


Supplementary Figure S1. Singlet oxygen-mediated effects. 12 500 208Fsrc3 cells in 96 well plates, 100 µl complete medium and 20 ng/ml TGF-beta1 were incubated in the presence of the indicated concentrations of SNP. The assays contained the indicated concentrations of the peroxynitrite decomposition catalyst FeTPPS, the singlet oxygen histidine (HIS), the NOX inhibitor AEBSF and the catalase mimetic EUK-134. Control assays remained free of inhibitors. The percentages of apoptotic cells were determined after 19.5 hrs. The observed shift of the inhibition curves is due to singlet oxygen generation through the interaction between $H_2O_2$ and peroxynitrite, leading to the inactivation of catalase. The resultant increase in free $H_2O_2$ causes and increased consumption of NO. The effects of the inhibitors are highly significant ($p<0.001$).

for inhibition of $H_2O_2$-dependent HOCl signaling. Therefore, the SNP-dependent decrease in HOCl signaling in the control curve reflects the counteraction between NO and $H_2O_2$ derived from transformed cells with inactivated catalase, whereas the curve in the presence of FeTPPS reflects the interaction between NO and $H_2O_2$ in the presence of partially protective catalase, as FeTPPS prevented peroxynitrite and subsequent singlet oxygen generation.

*2. Interaction of SNP-derived NO with HOCl signaling of the murine fibrosarcoma cell line L929*

The murine fibrosarcoma cell line L929 establishes HOCl signaling after inhibition of catalase [10, 11]. When the effects of two-fold dilution steps of the catalase inhibitor 3-AT were analysed according to a standard protocol, a smooth optimum curve of apoptosis induction was recognized (Supplementary Figure S2 A). However, application of very fine steps of variation of the 3-AT concentrations within the optimal range of apoptosis induction demonstrated that the true optimum curve was exceptionally sharp (Supplementary Figure S2 A). This finding illustrates the limitation of standard analysis and the potential of a sharper analysis based on a more refined analysis.

Addition of SNP to assays with increasing concentrations of 3-AT confirmed that apoptosis induction at the suboptimal and optimal concentration of 3-AT was directly inhibited by SNP, whereas SNP brought supraoptimal apoptosis induction at 50 mM 3-AT back to its optimum. Optimal apoptosis induction was then inhibited upon further application of higher concentrations of the NO donor (Supplementary Figure S2 B). These findings confirm the antagonistic action between NO and $H_2O_2$-dependent HOCl signaling, as shown for other cell lines in the main manuscript.

Supplementary Figure S2

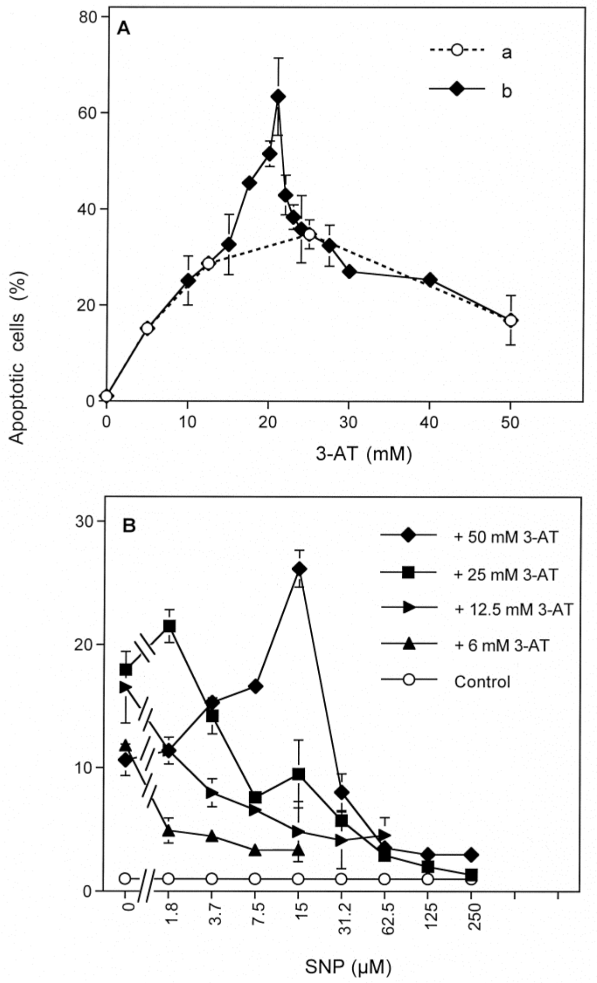


Supplementary Figure S2: The NO donor SNP antagonizes $H_2O_2$-dependent HOCl signaling of the murine fibrosarcoma cell line L929.

A. Determination of the optimum for apoptosis induction by 3-AT. L929 cells were cultivated in the presence of 20 ng/ml TGF-beta and the indicated concentrations of 3-AT for 45 hs. Under a, twofold dilution steps of 3-AT were applied, whereas under b smaller differences in the concentration of the catalase inhibitor were applied. The more refined analysis allows to determine the sharp optimum curve of apoptosis induction. B. The NO donor SNP inhibits apoptosis induction mediated by suboptimal and optimal concentrations of 3-AT. When added to supraoptimal concentrations of 3-AT it enhances apoptosis until the optimum is reached and then inhibits apoptopsis in a concentration-dependent mode.

*Statistical analysis: Apoptosis induction by 3-AT in Supplementary Figure S2 A and B, the difference between a and b (SupplementaryFigure S2A) in the concentration range of 20 mM 3-AT, as well as the effects of SNP on 3-AT-mediated apoptosis induction were highly significant ($p < 0.001$).*

*3. Three distinct effects of SNP-derived NO on apoptosis-inducing signaling in MKN-45 cells depend on the concentration of the NO donor*

As shown in Supplementary Figure S3 A, MKN-45 respond to increasing concentrations of the catalase inhibitor 3-AT with apoptosis induction in the mode of an optimum curve. As shown by the inhibition profile and in line with Figure 7 in the main manuscript, apoptosis induction was inhibited by AEBSF and therefore seems to be dependent on NOX1-derived superoxide anions at all concentrations of the catalase inhibitor. In the lower concentration range of 3-AT, NO/peroxynitrite signaling is reactivated. This is followed by HOCl signaling that shows a final decrease at high concentrations of 3-AT. This supraoptimal decline is explained by consumption of HOCl by excess $H_2O_2$ under the conditions of extreme inhibition of catalase [11]. In line with Figure 7 in the main manuscript, NO is been shown to i) enhance NO/peroxynitrite signaling at low concentrations of 3-AT, ii) inhibit HOCl signaling at intermediate concentrations of 3-AT, and to iii) interfere with supraoptimal decline of HOCl signaling at very high concentrations of 3-AT (Supplementary Figure S3 B). These three distinct effects are all strictly dependent on the concentration of the NO donor. This is in line with a role of NO in each one of these biochemical steps. It is important to keep in mind that the SNP-dependent increase in apoptosis induction in range of higher concentrations of 3-AT is not due to establishment of NO/peroxynitrite signaling, but is due to uprogation of the supraoptimal decline of HOCl signaling by SNP (please see Figure 7 in the main manuscript for details).

Supplementary Figure S3

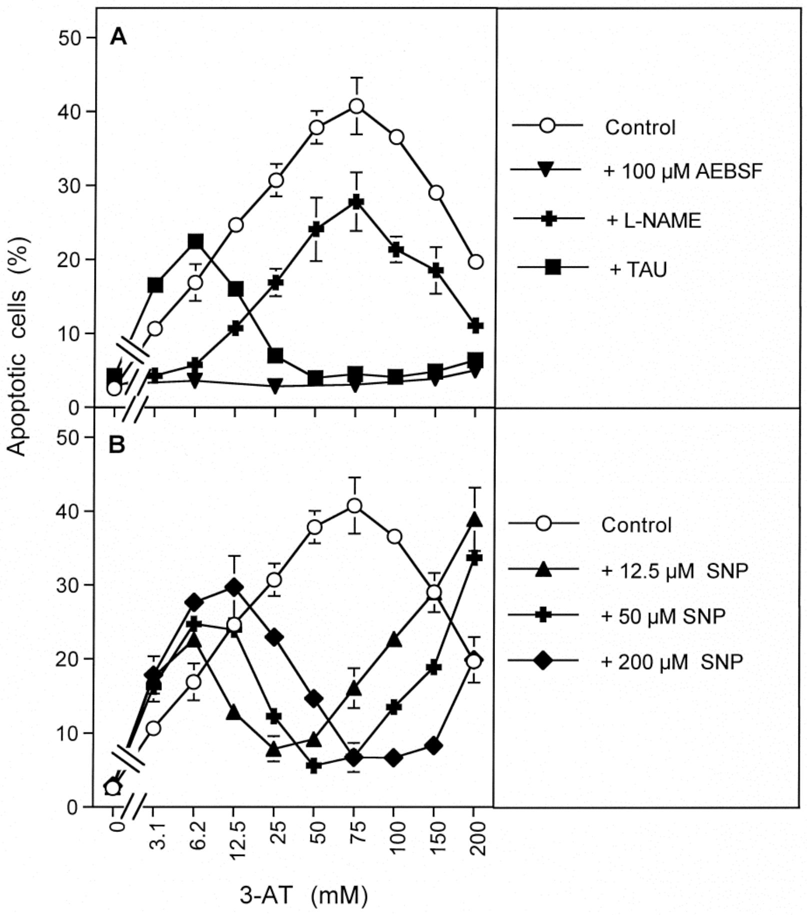


Supplementary Figure S3. The effects of SNP are dependent on its concentration.
12 500 MKN-45 gastric carcinoma cells in 96 well plates, 100 µl medium remained without additions (controls) or received 100 µM AEBSF, 2.4 mM L-NAME, 50 mM taurine or the indicated concentrations SNP. Then 3-AT was added at the indicated concentrations and the percentages of apoptosis were determined after 5 hs. The assays under A show that catalase inhibition leads to NOX1-dependent apoptosis induction, based on NO/peroxynitrite signaling at low 3-AT concentrations and on HOCl signaling at high concentrations. Apoptosis induction in controls appears as optimum curve, due to consumption of HOCl by excess H2O2 at high concentrations of 3-AT. The assays under B demonstrate that the NO donor SNP increases NO/peroxynitrite signaling, inhibits HOCl signaling at intermediate concentrations of 3-AT and counteracts the supraoptimal inhibition of HOCl signaling at high concentrations of 3-AT in a concentration-dependent mode.
*Statistical analysis: The enhancement of NO/peroxynitrite signaling, inhibition of HOCl signaling and abrogation of supraoptimal decline by NO were highly significant ($p<0.001$).*

*4. Inhibition of HOCl signaling by the NOD inhibitor quercetin and its derivatives quercitrin, isoquercitrin and rutin.*

Treatment of L929 fibrosarcoma cells with the catalase inhibitor 3-AT causes establishment of the HOCl signaling pathway. Addition of increasing concentrations of the NOD inhibitor quercetin or its glycosylated derivatives quercitrin, isoquercitrin and rutin caused concentration-dependent inhibition of apoptosis induction, followed

Supplementary Figure S4

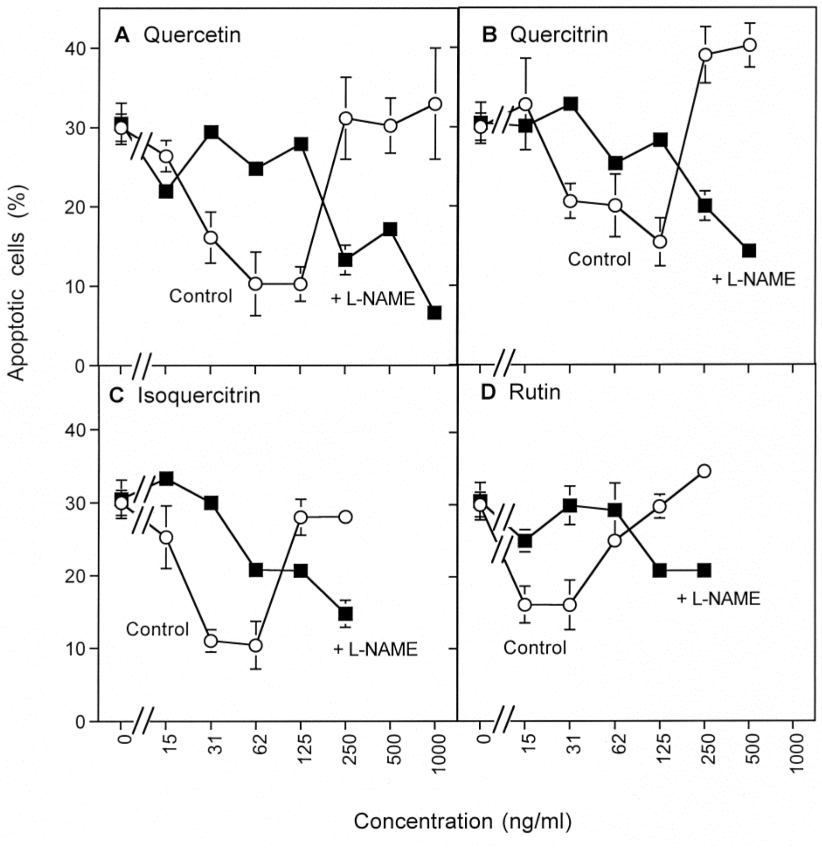


Supplementary Figure S4. Inhibition of NO dioxygenase (NOD) by quercetin and its derivatives inhibits $H_2O_2$-dependent HOCl signaling and establishes NO/peroxynitrite signaling in 3-AT-treated L929 fibrosarcoma cells

L929 cells in the presence of 20 ng/ml TGF-beta1 and 25 mM 3-AT were treated with the indicated concentrations of the NOD inhibitors quercetin, quercitrin, isoquercitrin and rutin, both in the absence and presence of 2.4 mM of the NOS inhibitor L-NAME and cultivated for 22 hs. Determination of apoptosis induction showed that all four compounds inhibited 3-AT-mediated apoptosis induction in an NO-dependent mode. After reaching maximal inhibition of the initial HOCl-dependent process, the compounds established NO-dependent apoptosis signaling. *Statistical analysis: Inhibition of 3-AT-mediated apoptosis induction by all four compounds, resumption of apoptosis induction at high concentrations of the compounds as well as inhibition of both processes by L-NAME were highly significant ($p < 0.001$).*

by reestablishment of apoptosis induction at higher concentrations of these compounds. Inhibition of apoptosis-inducing signaling as well as resumption of apoptosis induction was efficiently inhibited by the NOS inhibitor L-NAME: These findings are in line with the concept that increasing concentrations of NO after inhibition of NOD inhibit HOCl signaling and establish NO/peroxynitrite signaling when applied at higher concentrations. The quercetin moiety of the compounds studied seems to be required and sufficient to trigger the observed effects.

*5. Dual role of superoxide anions for NO/peroxynitrite signaling*

NO/peroxynitrite signaling depends on the concentrations of the two reaction partners NO and superoxide anions. These interact and form peroxynitrite. It is obvious that an increase in the concentration of superoxide anions might enhance peroxynitrite formation in the presence of sufficiently high concentrations of NO. However, an increase in the concentration of superoxide anions also leads to an increased $H_2O_2$ concentration, which might then consume NO and thus inhibit NO/peroxynitrite signaling. As low dose gamma radiation efficiently enhances NOX1-dependent superoxide anion generation [12, 13], this instrument was chosen for the modulation of the concentration of extracellular superoxide anions.

When untreated 208Fsrc3 cells or 208Fsrc3 cells irradiated with increasing doses of gamma irradiation where treated with the slowly decaying NO donor SNP, apoptosis induction was dependent on the concentration of SNP (Supplementary Figure S5 A).

Supplementary Figure S5

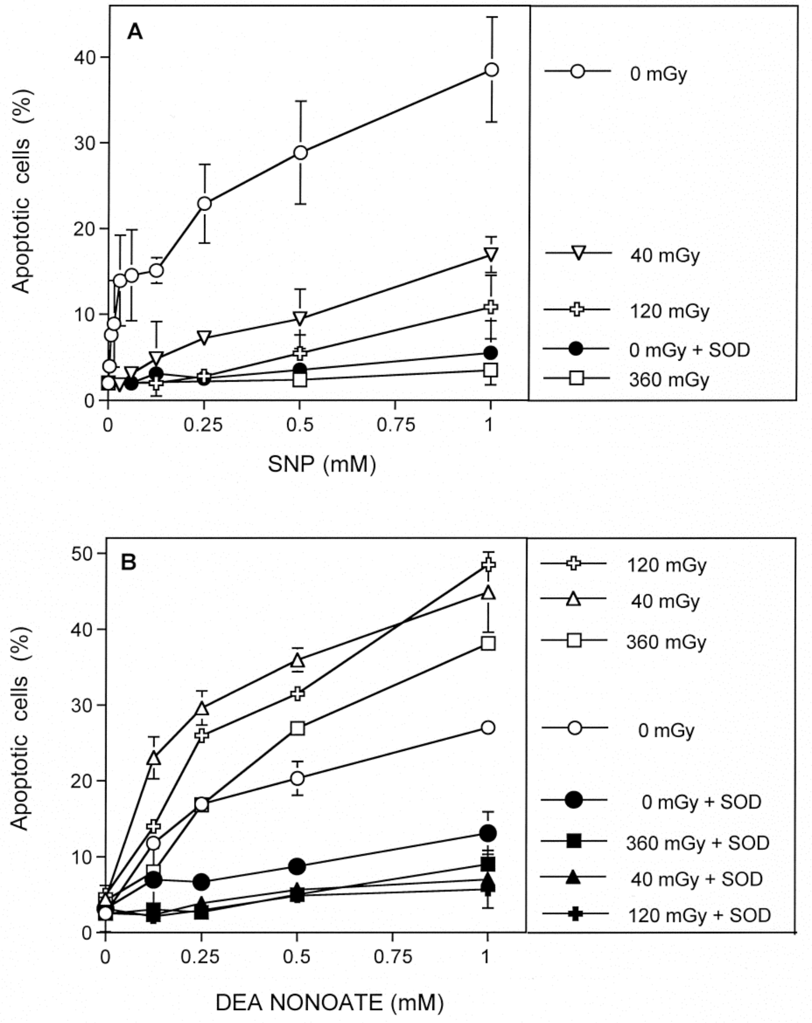


Supplementary Figure S5
Differentiation between the effects of superoxide anions and $H_2O_2$ on NO/peroxynitrite signaling
Unirradiated 208Fsrc3 cells and 208Fsrc3 cells irradiated with the indicated dose of gamma radiation were seeded at a density of 3000 (A) or 4000 cells/assay (B). Where indicated, assays received 100 U/ml MnSOD. The indicated concentrations of the NO donor SNP (A) or DEA NONOate (B) were added. Apoptosis induction was determined after 48hs (A) or 12.5 hs (B).
Supplementary Figure S5 A shows at 48 hs that apoptosis induction is maximal in unirradiated cells and is suppressed by irradiation in a dose-dependent mode.When the fast decaying NO donor DEA NONOate is applied (B), irradiation enhances NO-mediated apoptosis induction up to a radiation dose of 120 mGy. Increasement to 360 mGy leads to mild inhibition.
*Statistical analysis: A: Apoptosis induction by SNP and its inhibition by irradiation and by SOD were highly significant ($p < 0.001$). B: Apoptosis induction by DEA NONOate and its enhancement by 40 mGy and 120 mGy irradiation, as well as inhibition by SOD were highly significant at all concentrations of DEA NONOate, whereas enhancement of DEA NONOate-dependent apoptosis induction by 360 mGy irradiation was only highly significant at 1 mM DEA NONOate.*

Apoptosis induction under these conditions was inhibited by SOD, indicating that the reaction between NO and superoxide anions, i. e. formation of peroxynitrite, was required. Pretreatment with increasing doses of gamma irradiation caused inhibition of apoptosis induction dependent on the dose. 40 mGy treatment caused already more than 50 percent inhibition, 360 mGy caused complete inhibition. These findings show that in the presence of limiting concentrations of NO, an increase in superoxide anions after irradiation has no chance to increase peroxynitrite formation and apoptosis induction, as the concomittant increase in $H_2O_2$ seems to efficiently counteract NO/peroxynitrite signaling through consumption of NO.

This picture changed significantly when the fast decaying NO donor DEA NONOate was applied instead of the slowly decaying NO donor SNP (Supplementary Figure S5 B). DEA NONOate caused concentration-dependent apoptosis induction, mediated by peroxynitrite, as the reaction was blocked by SOD. Pretreatment of the cells with gamma irradiation enhanced DEA NONOate-dependent apoptosis induction. Thereby 40 mGy gave optimal enhancement, whereas a further increase in dose lowered the enhancing effect. These data show that in the presence of abundant NO, released by the fast decaying NO donor, an increase in superoxide anions allows for more peroxynitrite formation and an enhanced biological effect. The negative effect of excess $H_2O_2$ due to irradiation is also significant, but cannot outcompete the positive effect on NO/peroxynitrite signaling.

In line with these results, pretreatment of 208Fsrc3 cells with low dose irradiation had a significant effect on their interaction with SNP. As shown in Supplementary Figure S6, autocrine apoptosis induction was initially increased by SNP, followed by a strong inhibition of apoptosis. Further increase in SNP caused resumption of apoptosis induction. This finding is identical to the data shown in Figure 4 of the main manuscript. Pretreatment with low dose irradiation (100 mGy) caused a rightward shift of the curve, indicative of a higher overall concentration of $H_2O_2$ that requires higher concentrations of SNP-derived NO for counteraction.

Supplementary Figure S6

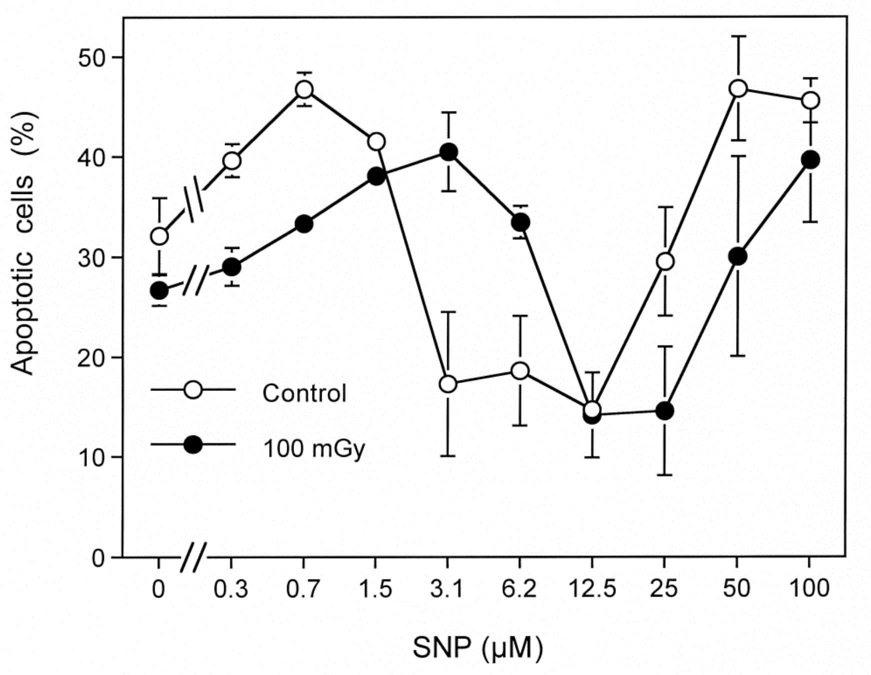


Supplementary Figure S6. Low dose radiation induces consumption of NO.
208Fsrc3 cell were irradiated with 100 mGy gamma irradiation or remained untreated. The cells were then seeded at a density of 12 500 cells per assay in 96 well plates with 100 µl medium containing 20 ng/ml TGF-beta1. The indicated concentrations SNP were added and apoptosis induction was determined after 27 hrs. The result shows that autocrine apoptosis induction by the transformed cells is slightly enhanced at very low concentrations of SNP, followed by inhibition at higher concentrations. At very high concentrations, apoptosis induction resumed again. This result was analogous to that described in Figure 4, main manuscript. Irradiated cells, which generate higher concentrations of superoxide anions (and therefore also more H2O2) showed the same signaling profile, however at higher concentrations of SNP. This shift indicates the consumption of NO which can be compensated by higher concentrations of the NO donor SNP.
*Statistical analysis: The shift induced by irradiation was highly significant ($p<0.001$).*

6. *The potential role of ferrous/ferric ions for the antagonistic action between hydrogen peroxide and NO*

Previous experiments (Figure 7 in the main manuscript) have shown that 200 mM 3-AT caused supraoptimal apoptosis induction in MKN-45 cells, i. e. a situation in which excess $H_2O_2$ partially inhibited HOCl signaling through interaction with HOCl. The data presented in Supplementary Figure S7 A show that increasing concentrations of SNP initially shift supraoptimal apoptosis induction up to an optimum and then cause its inhibition. Then, finally, NO/peroxynitrite signaling is resuming. The presence of the iron chelator desferroxamine interferes with SNP-mediated enhancement of supraoptimal apoptosis induction as well as with SNP-mediated inhibition of optimal apoptosis induction (Supplementary Figure S7). This finding indicates the involvement of iron in both processes.

DEA NONOate-dependent apoptosis induction in MKN 45 tumor cells in the presence of the catalase inhibitor 3-AT was inhibited by increasing concentrations of $H_2O_2$-generating glucose oxidase (GOX) (Supplementary Figure S8). After complete inhibition, GOX-dependent apoptosis resumed. The inhibitory effect of GOX-derived $H_2O_2$ on NO/peroxynitrite signaling was completely abrogated in the presence of deferroxamine, indicating that this process was dependent on iron. This figure also shows that the concentration-dependency of GOX-mediated apoptosis induction (in the presence of 3-AT, but absence of the NO donor) was shifted rightward through DEA NONOate. Therefore, the balancing effect of $H_2O_2$ on NO-dependent apoptosis-inducing processes seems to be reciprocal.

Supplementary Figure S7

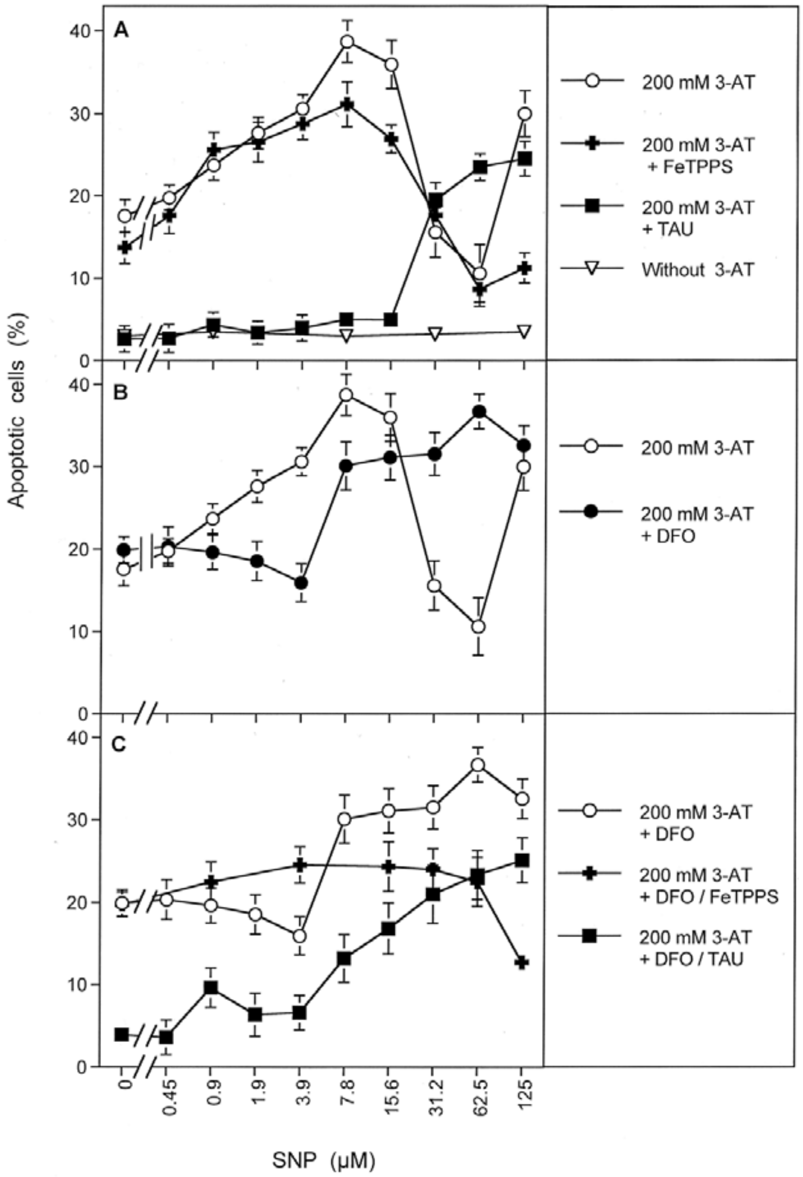


Supplementary Figure S7. The interference of NO with $H_2O_2$-mediated inhibition of HOCl signaling at high concentrations of the catalase inhibitor 3-AT is mediated by iron. 12 500 MKN-45 cells in 96 well plates with 100 µl medium received 200 mM 3-AT or remained free of 3-AT. Where indicated, the following additions were made: FeTPPS (20 µM), taurine (5 mM), DFO (50 µM). All assays received the indicated concentrations of SNP. The percentages of apoptotic cells were determined after 4.5 hrs.

*Statistical analysis: the interfering effect of DFO is highly significant ($p<0.001$).*

Supplementary Figure S8

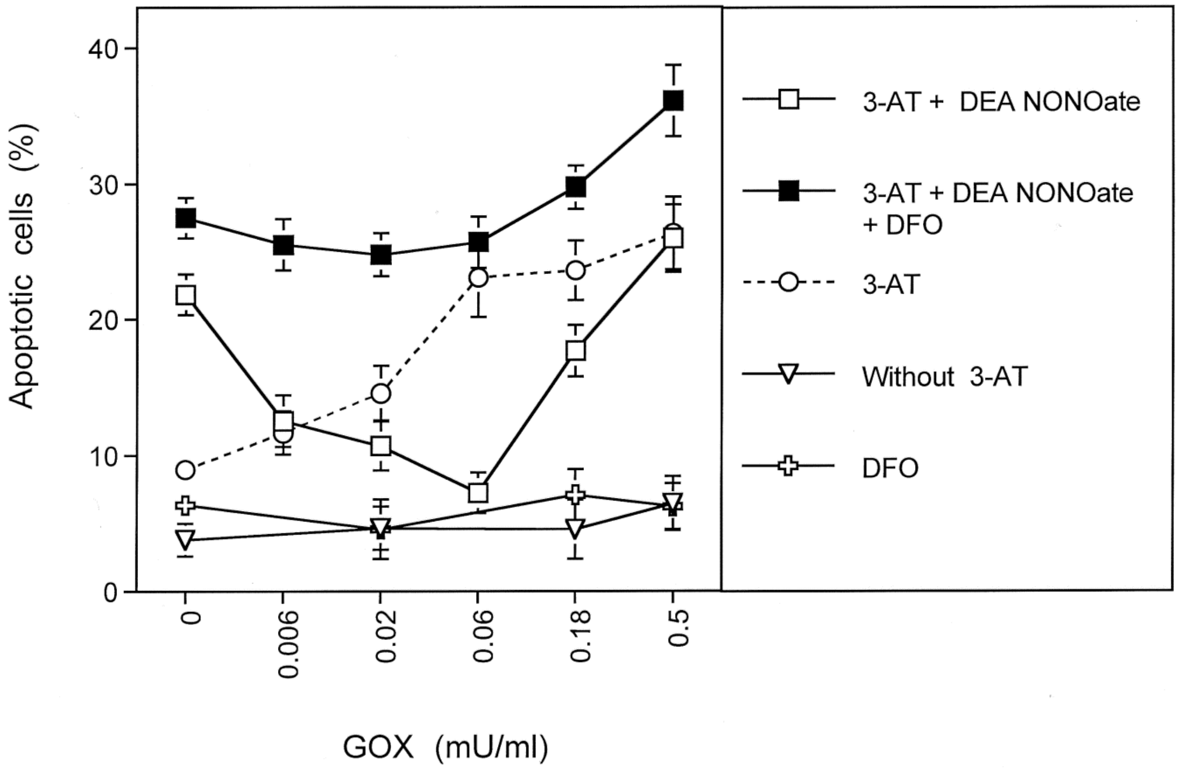


Supplementary Figure S8. Counteraction of $H_2O_2$ towards NO/peroxynitrite signaling is dependent on iron. 12 500 MKN-45 cells in 96 well plates, 100 µl medium, remained without 3-AT or received 10 mM 3-AT. Where indicated, 200 µM DEA NONOate and/or 50 µM DFO were added. The percentages of apoptotic cells were determined after 3 hrs.

*Statistical analysis: the interfering effect of DFO is highly significant ($p<0.001$).*

*7. Tumor cells from different tissues show differences in the quality of their apoptosis-inducing signaling chemistry*

Different signaling profiles after addition of increasing concentrations of the catalase inhibitor 3-AT are summarized in Supplementary Figure S9. Whereas prostate carcinoma cells PC-3 showed an optimum curve for apoptosis induction, dependent on the concentration of 3-AT, the cervical carcinoma cell line SISO and the neuroblastoma cell line SHEP responded with constant increase in apoptosis

Supplementary Figure S9

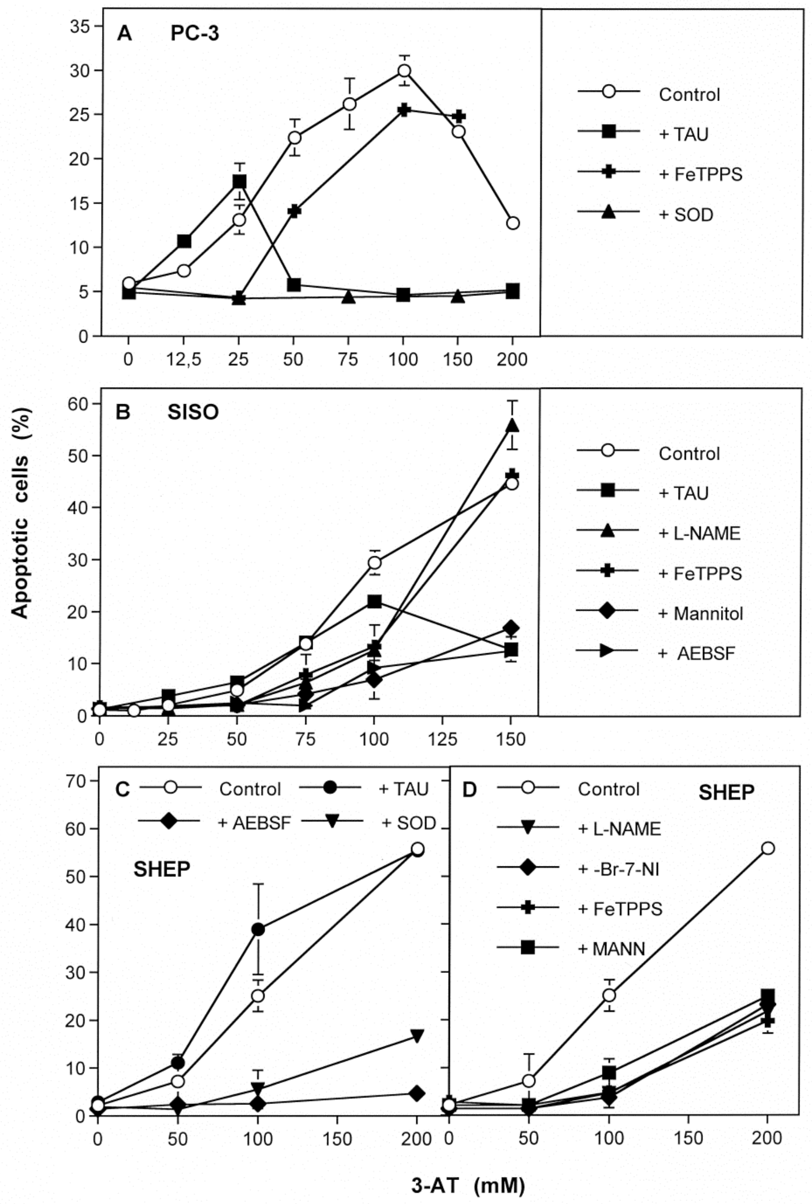


Supplementary Figure S9. Apoptosis induction in tumor cells: variable responses.

The human prostate carcinoma cell line PC-3 (A) (10 000 cells/assay), cervical carcinoma cell line SISO (B) (12 500 cells per assay) and the neuroblastoma cell line SHEP (C) (12 500 cells per assay) were seeded in 96 well plates and indicated in the absence or presence of the indicated inhibitors/scavenger and increasing concentrations of 3-AT. Concentrations of inhibitors: taurine (TAU) 50 mM; FeTPPS 20 µM, SOD 100 U/ml, L-NAME 2.4 mM, mannitol 20 mM, AEBSF 100 µM, 3-Br-7-nitroimidazole (3-Br-7-NI) 20 µM. 3-Br-7-NI is a specific inhibitor of nNOS. The percentages of apoptotic cells were determined after 5.5 hrs (A), or 22 hrs (B), or 7 hrs (C). PC-3 and SISO cells first show NO/peroxynitrite signaling and then HOCl signaling, but differ in the characteristic of the concentration-dependency curve: Optimum curve, in the case of PC-3 (indicative of $H_2O_2$-mediated inhibition of HOCl signaling at high concentrations of catalase inhibitor) and constant increasing response in the case of SISO cells. SHEP cells show NO/peroxynitrite signaling exclusively after catalase inhibition.

*Statistical analysis: apoptosis induction mediated by 3-AT and the specific inhibitions as outlined in the text were highly significant.*

induction. Whereas PC-3 and SISO cells showed apoptosis induction through the NO/peroxynitrite signaling pathway at lower concentrations of 3-AT and through the HOCl signaling pathway at higher concentrations, SHEP cells showed NO/peroxynitrite signaling at all concentrations of the catalase inhibitor.

When optimal superoxide anion production was warranted through addition of exogenous TGF-beta1, tumor cell lines like SISO or SIHA (cervical carcinoma) and the melanoma cell lines IGR-37 and IGR-39 showed mainly HOCl signaling in the presence of 100 mM 3-AT (Supplementary Figure S10), whereas the mammary carcinoma cell lines CAL-51, HCC-1937 and MCF-7, as well as the ovarial carcinoma cell line FU-OV-1 showed NO/peroxynitrite signaling exclusively under these conditions (Supplementary Figure S11).

Supplementary Figure S10

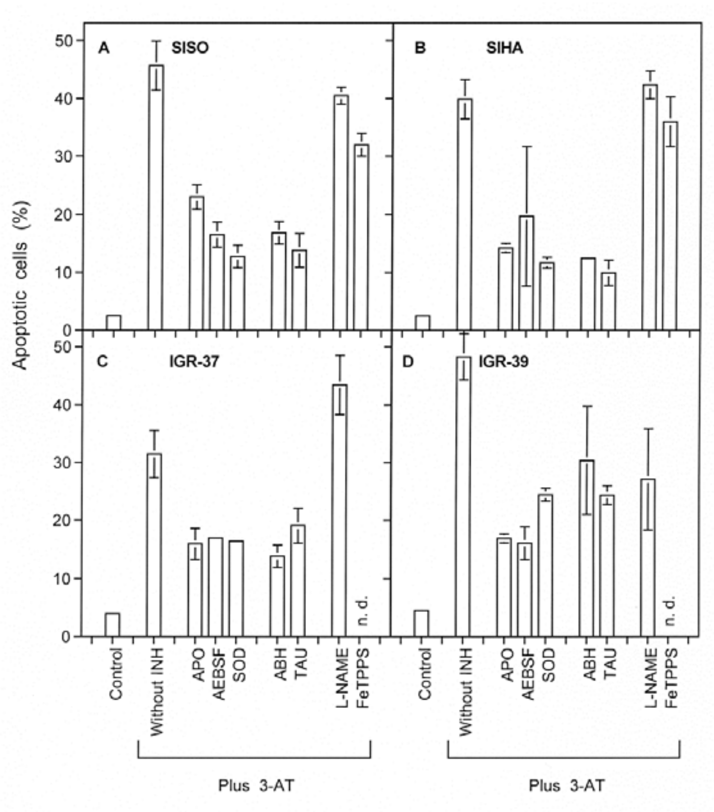

Legend to Supplementary Figure S10. Autocrine apoptosis induction by tumor cells.
25 000 cells in 200 µl medium, 48 well plates, of the human cervical carcinoma cell lines SISO and SIHA, and the melanoma cell lines IGR-37 and IGR-39 were seeded in the presence of 20 ng/ml TGF-beta and the indicated inhibitors/scavengers. Where indicated, 3-AT was added at a concentration of 100 mM. The percentages of apoptotic cells were determined after 24 hrs in A, C, D and 36 hrs in B.
Concentrations of inhibitors: apocynin (APO) 50 µg/ml; AEBSF (100 µM), SOD (100 U/ml; ABH 150 µM; taurine (TAU) 50 mM, L-NAME 2.4 mM; FeTPPS 25 µM.
Under the conditions of the test, SISO, SIHA and IGR-37 cells only show HOCl signaling, wherease IGR-39 cells show indications of both pathways.
*Statistical analysis: apoptosis induction mediated by 3-AT and its inhibition by compounds that interfere with HOCl signaling was highly significant*

Suppplementary Figure S11

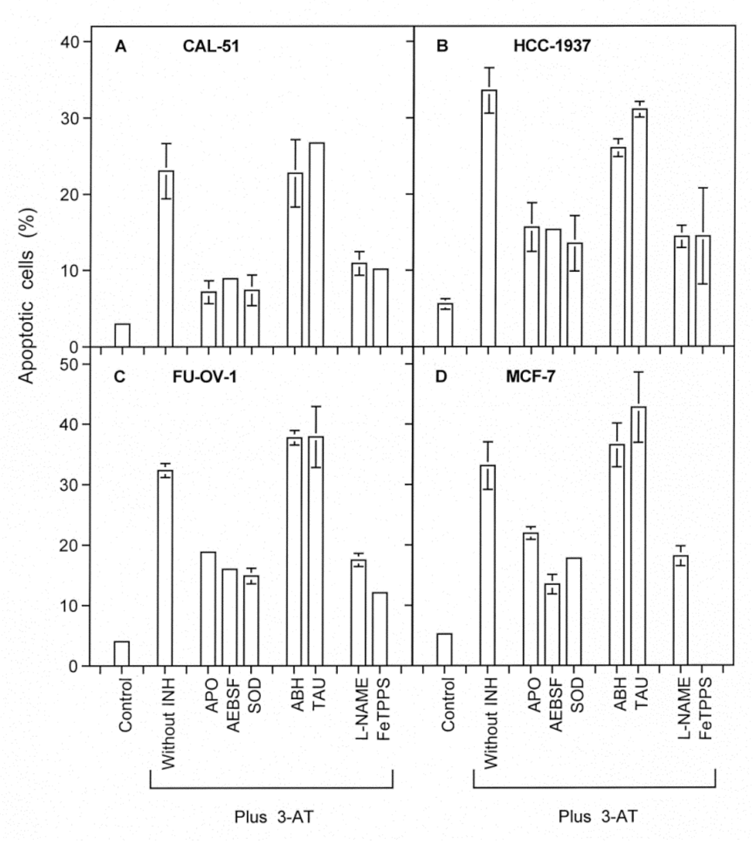


Supplementary Figure S11. Autocrine apoptosis induction by tumor cells.
25 000 cells in 200 µl medium, 48 well plates, of the human mammary carcinoma cell lines CAL-51, HCC-1937 and MCF-7, and the ovarial cell line FU-OV-1 were seeded in the presence of 20 ng/ml TGF-beta and the indicated inhibitors/scavengers. Where indicated, 3-AT was added at a concentration of 100 mM. The percentages of apoptotic cells were determined after 24 hrs in A, C, D and 36 hrs in B.
Concentrations of inhibitors: apocynin (APO) 50 µg/ml; AEBSF (100 µM), SOD (100 U/ml; ABH 150 µM; taurine (TAU) 50 mM, L-NAME 2.4 mM; FeTPPS 25 µM.

All four tumor cell lines seem to respond with NO/peroxynitrite signaling after catalase inhibition.

*Statistical analysis: apoptosis induction by 3-AT and its inhibition by inhibitors of the NO/peroxynitrite pathway were highly significant Statistical analysis: the interfering effect of DFO is highly significant (p<0.001).*

*8. The determining role of the effector function for the quality of intercellular ROS/RNS-dependent apoptosis-inducing signaling. The use of transformed target cells.*

Pottgiesser et al. [14] have established an analytical cell culture system that allows to differentiate between the "target function" and the "effector function" during intercellular ROS/RNS-mediated apoptosis induction. This system is schematically presented in Supplementary Figure S12. The "target cell function" thereby is characterized as the ability to respond to intercellular ROS/RNS signaling with induction of apoptosis. It has been found to be restricted to malignant cells and depend on the generation of extracellular superoxide anions by NOX1. If tumor cells, i. e. cells from late stages of oncogenesis are used as target cells, their membrane-associated catalase needs to be inhibited for activation of the target function. The "effector cell function" has been characterized as the potential to release either DUOX-coded peroxidase or NO. The effector function can be exerted by non-malignant and malignant cells of different stages. For experimental analysis, clumps of transformed cells are surrounded by a large volume of medium. Under these conditions, peroxidase and/or NO released from the cells is diluted into the medium and thus the signaling pathways are not established, despite active NOX1. Surrounding the clumps with large numbers of cells, but at low density, allows to supplement for the necessary POD or NO to establish apoptosis induction in the clumps, without enabling the effector cells to show their own target cell potential during the experiment. As the specific target function is restricted to NOX1 expressing cells, nontransformed cells in clumps surrounded by effector cells do not show apoptosis induction. The effector function is independent of the malignant state. Cells seeded under the conditions of effector cells in this assay show no apoptosis

Supplementary Figure S12

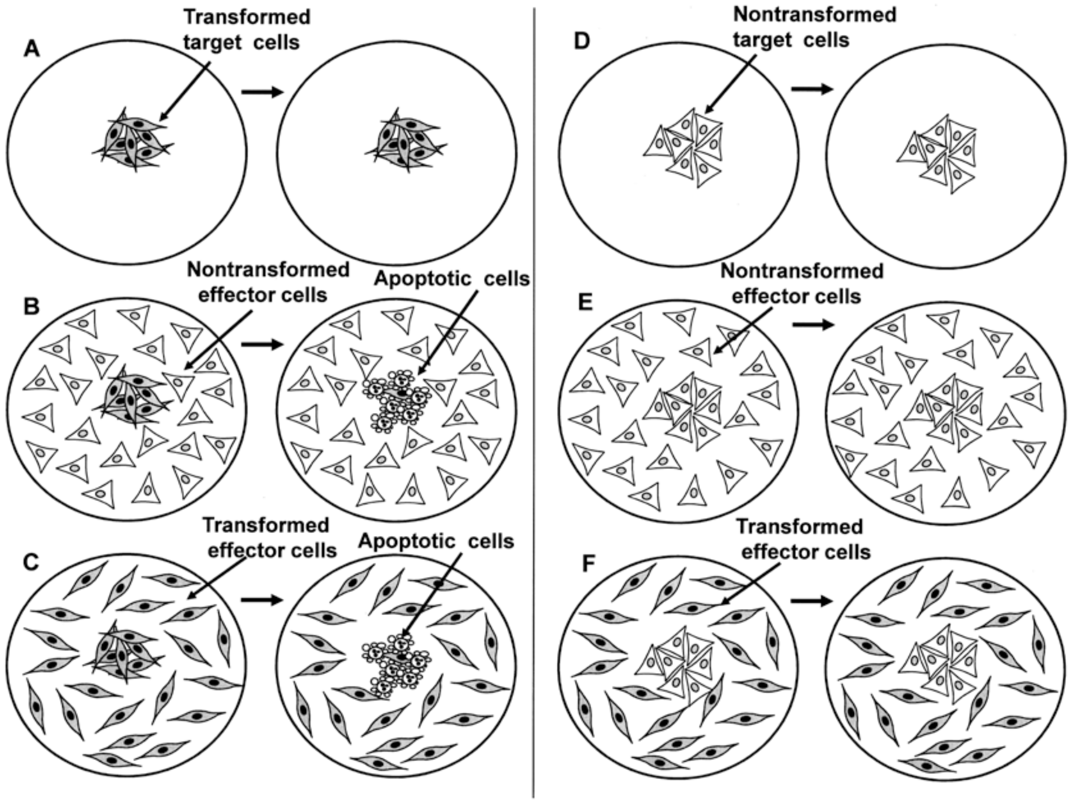


Supplementary Figure S12: **Schematic presentation of the cell system used for the experimental dissection of target and effector cell function of intercellular ROS-mediated induction of apoptosis**. Part I: Analysis of nontransformed and transformed effector cells. Transformed (A) or nontransformed (D) cells were seeded as clumps of high local density (250 cells/$mm^2$), but low total cell number (2000 cells) in 12 well tissue culture clusters and 1 ml medium containing 20 ng/ml TGFβ1. In addition, target cells were surrounded with either 15 000 nontransformed (B,E) or transformed (C, F) effector cells. The effector cells had a starting density of 40 cells/$mm^2$. Clumps of target cells cultivated alone did not show significant induction of apoptosis (A, D). Transformed target cells, surrounded by either nontransformed (B) or transformed (C) effector cells show substantial induction of apoptosis, whereas clumps of nontransformed target cells did not. This experimental system allows to define target and effector cell functions of defined cell lines. ***This approach was applied for the analysis shown in Supplementary Figure S13***.

Part II: Analysis of tumor cell effector and target cell function. These experiments follow the principle outlined in this figure, with slight modifications: The assays were performed in 24 well plates, with only 0.5 ml medium, 1500 target cells at high density (250 cells/$mm^2$) and 8000 effector cells. No TGF-beta was added. This approach was applied in the experiments described in Supplementary Figures 16-24. Inhibition of catalase was essential to allow expression of the target function. In combination with inhibitors or siRNA-treatment of cells this approach allowed to study the signaling chemistry underlying intercellular induction of apoptosis (Supplementary Figures S21-S24).

induction during the time of the experiment, even if they are transformed. This failure is due to their low density that does not allow sufficient generation of $H_2O_2$.

The experiment described in Supplementary Figure S13 used four types of cells as target cells. that were seeded as clumps. These were nontransformed 208F rat fibroblasts, their src-oncogene-transformed derivative 208Fsrc3, nontransformed rat epithelial cells ROSE 199 and their HRAS-transformed derivative ROSE A2/5. In the control part under A, the clumps either remained without effector cells or were surrounded by an excess of 208F, 208Fsrc3 or ROSE A2/5 cells of low density. Clumps seeded alone showed no or only a minor apoptotic response. Clumps of nontransformed cells showed no apoptotic response, even if they were surrounded by effector cells. In contrast, clumps of transformed cells responded with apoptosis in the presence of each one of the three types of effector cells ($p<0.001$). Inhibitor studies thereby showed that the apoptotic response of the transformed cells in the presence of effector cells was strongly inhibited in the presence of the HOCl scavenger taurine ($p<0.001$), whereas the NOS inhibitor L-NAME only had a marginal inhibitory effect. These findings show that the effector cells used in the experiment described under A mainly supported the signaling process with peroxidase that was released by the effector cells and acted in trans selectively on the transformed cells. The use of various tumor cells as effector cells in this system showed that the cervical tumor lines SIHA and SICO exerted effector function mainly through peroxidase ($p<0.001$), whereas the mammary carcinoma cell lines CAL-51 and the ovarial carcinoma cell lines Fu-OV-1 and BG-1 contributed to signaling essential through the release of NO (p<0.001).

Supplementary Figure S13

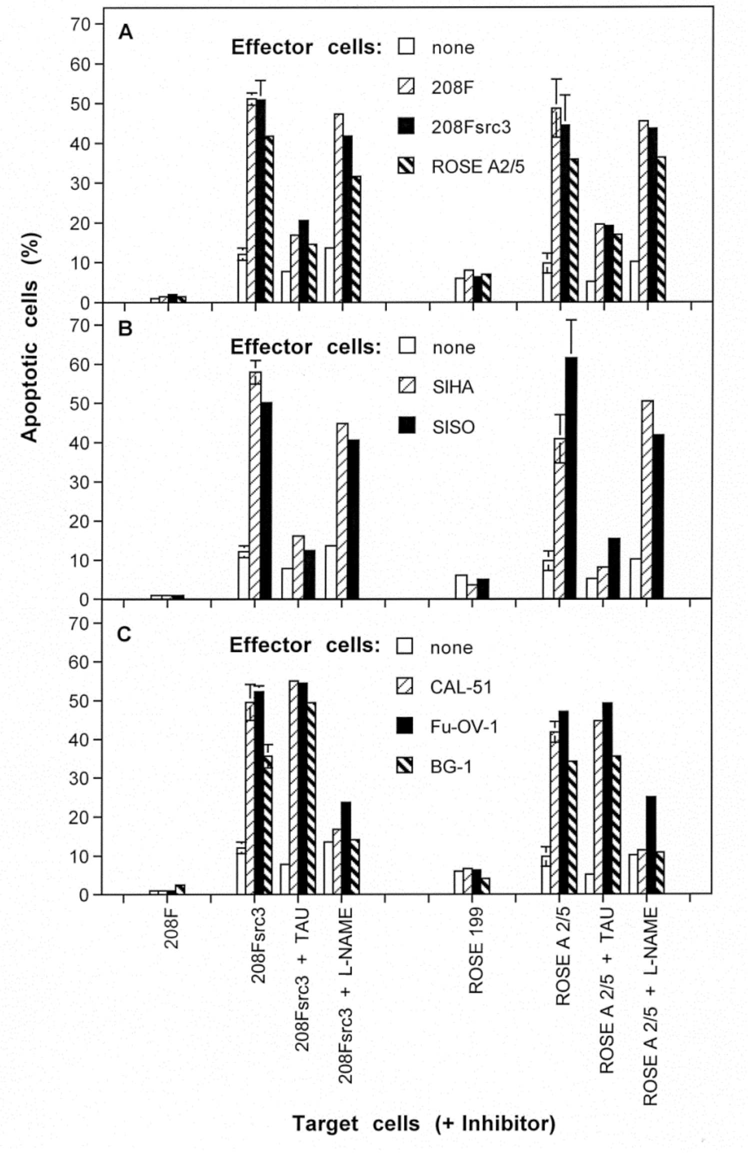


Supplementary Figure S13. Determination of target and effector function of nontransformed and transformed rat fibroblasts and epithelial cells. The experiment was performed as described in Supplementary Figure S12 in 12 well tissue culture clusters. Nontransformed rat fibroblasts 208 F, their src-oncogene-transformed descendants 208Fsrc3, nontransformed rat epithelial cells ROSE 199 and their transformed counterparts ROSE A2/5 were seeded as clumps of high cell density, low total number. They were surrounded by 15 000 effector cells (high number, low density) or cultivated in the absence of effector cells. Where indicated, 50 mM taurine

(TAU) or 2.4 mM L-NAME were added. The percentages of apoptotic cells in clumps were determined after 2 days.
The results show that the target cell function can only be exerted by transformed cells, whereas the effector function is demonstrated for nontransformed cells and transformed cells. Effector function is also shown for the human tumor cell lines SIHA and SISO (cervical carcinoma), CAL-51 (mammary carcinoma) and Fu-OV-1 and BG-1 (ovarial carcinoma). The effector cells determine whether the transformed target cells are killed by HOCl-signaling or NO/peroxynitrite signaling. Epithelial cells and fibroblasts respond analogously. Epithelial effector cells have the same effect on transformed fibroblasts as on transformed epithelial cells and vice versa. These data show the general significance of the signaling system described by us.
*Statistical analysis: significance of specific findings is indicated in the text.*

These data show that the dominating release of either POD or NO from defined tumor cells corresponded to the type of apoptotic response after inhibition of catalase, i. e. preference for HOCl signaling or NO/peroxynitrite signaling.

Importantly, tumor cell lines from both categories determined in the previous figures were able to respond to NO/peroxynitrite-mediated apoptosis induction after application of an exogenous NO donor, provided their protective catalase was inhibited (Supplementary Figures S14 and S15.

Supplementary Figure S14

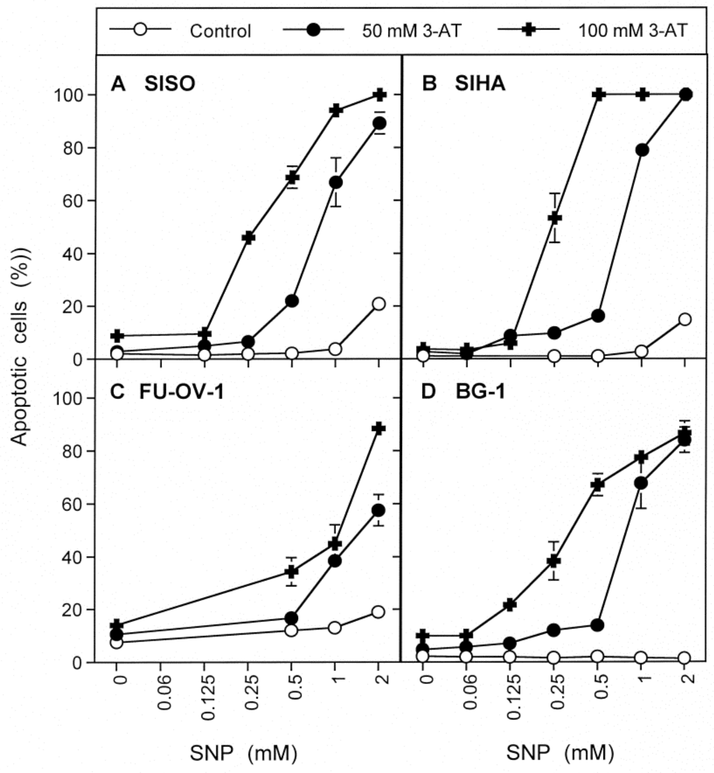

Legend to Supplementary Figure S14. Apoptotic response of tumor cells to an exogenous NO donor requires inhibition of their catalase and is independent of their own effector cell characteristic. The cervical carcinoma cell lines SISO and SIHA, and the ovarial carcinoma cell lines FU-OV-1 and BG-1 were seeded in 48 well tissue culture clusters (5000 cells/assay, 400 µl medium), in the absence of 3-AT or in the presence of either 50 mM or 100 mM 3-AT. SNP was added at the indicated concentrations and the percentages of apoptotic cells were determined after 24 hrs.
The result shows that the apoptotic response under the conditions of the experiment requires inhibition of catalase and the presence of the NO donor.
*Statistical analysis: Concentration-dependent induction of apoptosis by SNP in the presence of 3-AT was highly significant (p<0.001).*

Supplementary Figure S15

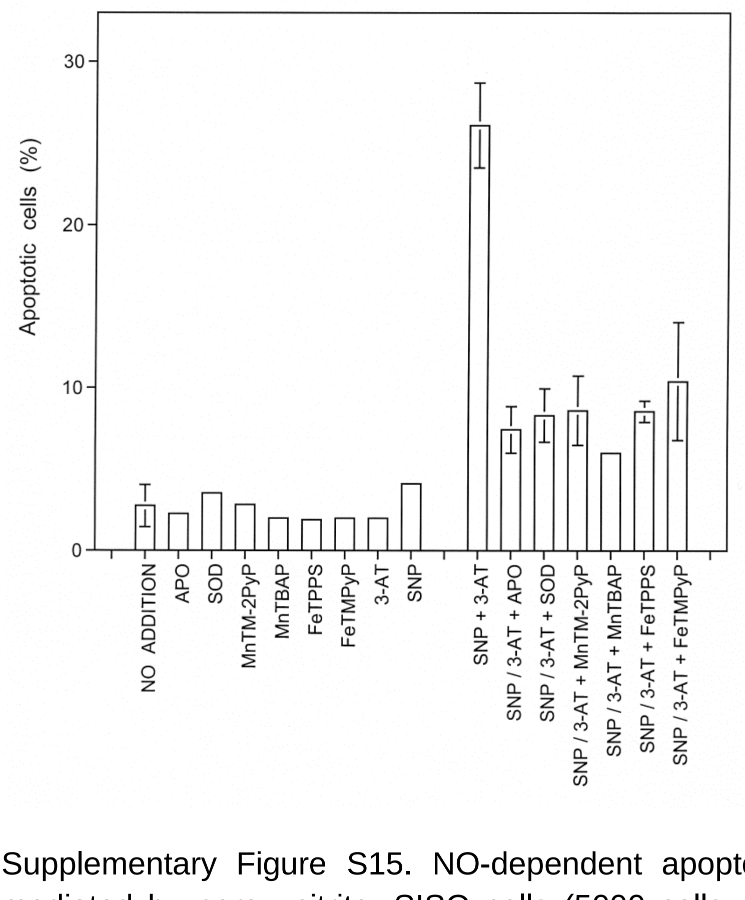


Supplementary Figure S15. NO-dependent apoptosis induction in SISO cells is mediated by peroxynitrite. SISO cells (5000 cells per assay, 48 well plate, 400 µl medium) were cultivated in the absence or presence of inhibitors, as well as in absence or presence of 25 mM 3-AT plus 1 mM SNP. The percentages of apoptotic cells were determined after 20 hrs. The following inhibitors were used: Inhibitor of NOX1: apocynin, 50 µg/l; scavenger of superoxide anions: SOD (100 U/ml); MnTM-2pyP (20 µM), MnTBAP (20 µM), Decomposition catalyst of peroxynitrite: FeTPPS (25 µM); FeTMPyP (25 µM). The results show that the apoptotic response towards the NO donor requires inhibition of catalase and is mediated by superoxide anions derived from NOX1 that interact with NO to generate peroxynitrite.

*Statistical analysis: Apoptosis induction by SNP plus 3-AT, as well as its inhibition by all inhibitors applied was highly significant ($p<0.001$).*

*9. Intercellular induction of apoptosis in bona fide tumor cells.Inhibition of protective catalase is necessary to activate the target function, whereas the effector function determines the quality of signaling.*

The next experiments were focussing on the elucidation of the characteristics of target and effector cell functions in bona fide tumor cells. Therefore, the experimental system presented in Supplementary Figures S12 and S13 was modified. Clumps of tumor cells were used (instead of transformed cells as in the previous experiment) and surrounded by tumor cells, or not. Two prototype tumor cell lines (with respect to their intercellular autocrine apoptosis-inducing ROS/RNS signaling) were used: SISO cells show HOCl preferentially at optimal inhibition of their protective catalase, whereas SHEP cells die through NO/peroxynitrite signaling in the presence of catalase inhibitor (please find details in Supplementary Figure S9). Clumps of SISO cells or SHEP cells showed strong apoptotic response as target cells only if they were i) surrounded by effector cells and ii) in the presence of the catalase inhibitor 3-AT (Supplementary Figures S16 and S17). SISO cells and SHEP cells acted equally well as effector cells with regard to the strength of apoptosis induction in the target cells. However, the quality of intercellular ROS/RNS signaling achieved by the effector cells was completely different. SHEP effector cells allowed apoptosis induction preferentially through the NO/peroxynitrite pathway both in SHEP and SISO target cells, whereas SISO effector cells caused apoptosis induction in these target cells essentially through the HOCl signaling pathway. The effector functions of SHEP and SISO effector cells can be completely mimicked by adding MPO or the NO donor SNP instead of the effector cells (Supplementary Figure S18, 19). The addition of MPO mimicked the effector function of SISO cells, whereas the NO donor mimicked the effector function of SHEP cells. These findings demonstrate that these two effector molecules are sufficient to determine the quality of intercellular

Supplementary Figure S16

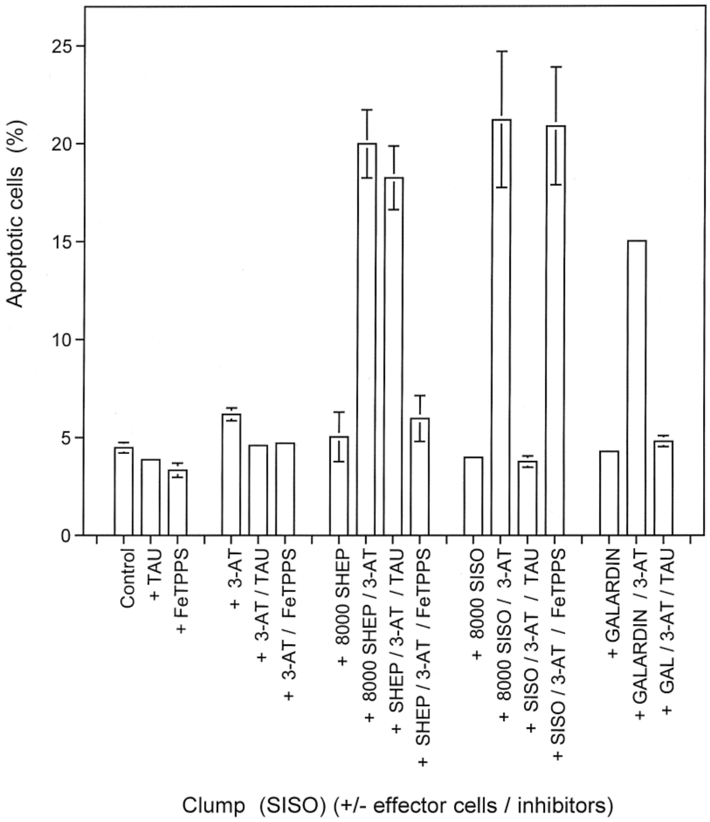


Supplementary Figure S16. The effect of SHEP or SISO effector cells on SISO target cells. Clumps of 1500 SISO cells were seeded and remained either isolated or were surrounded by 8000 SHEP or SISO cells, in the absence or presence of the indicated inhibitors/scavengers (3-AT: 100 mM; taurine (TAU): 50 mM; FeTPPS (25 µM). In addition, clumps without surrounding effector cells received 10 µM of the protease inhibitor galardin without further addition or together with 3-AT or 3-AT plus taurine.
After 15 hrs, the percentages of apoptotic cells were determined. Clumps in the absence of effector cells showed background apoptosis induction. The presence of either SHEP or SISO effector cells caused remarked apoptosis induction, provided the catalase inhibitor 3-AT was present. As seen from the differential inhibitory action of the HOCl scavenger taurine and the peroxynitrite decomposition catalyst FeTPPS, the presence of SHEP effector cells induced apoptosis in SISO target cells through NO/peroxynitrite signaling, wherease SISO effector cells caused apoptosis through HOCl signaling. In the presence of the matrix metalloprotease inhibitor galardin, the peroxidase domain of DUOX is not released [14, 15] and diluted in the medium.

*Statistical analysis: Apoptosis induction in the presence of SHEP or SISO effector cells plus 3-AT, as well as in the presence of galardin plus 3-AT was highly significant (p<0.001). The inhibition of apoptosis induction by FeTPPS in the case of SHEP effector cells, and by taurine, in the case of SISO effector cells or galardin was highly significant (p<0.001).*

Supplementary Figure S17

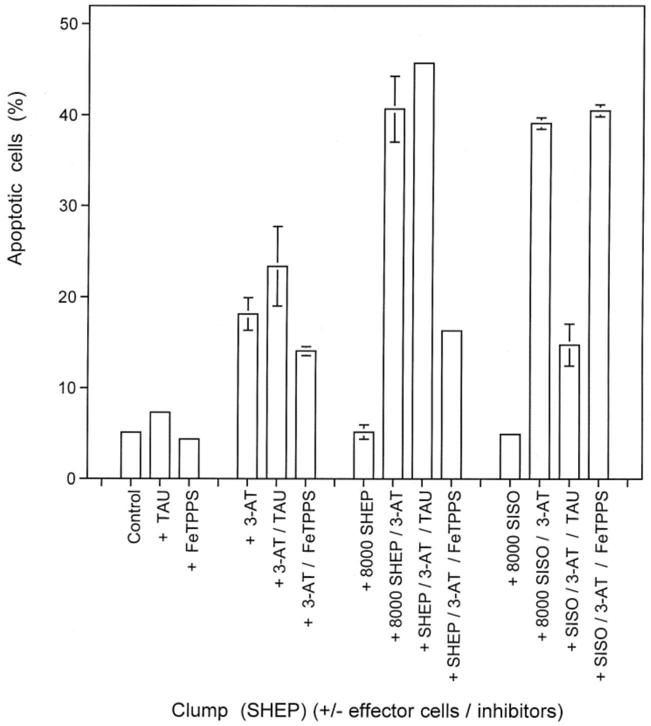


Supplementary Figure S17. The effect of SHEP or SISO effector cells on SEEP target cells. Clumps of 1500 SHEP cells were seeded and remained either isolated or were surrounded by 8000 SHEP or SISO cells, in the absence or presence of the indicated inhibitors/scavengers (3-AT: 100 mM; taurine (TAU): 50 mM; FeTPPS (25 µM).

After 15 hrs, the percentages of apoptotic cells were determined. Clumps in the absence of effector cells showed background apoptosis induction. The presence of either SHEP or SISO effector cells caused remarked apoptosis induction, provided the catalase inhibitor 3-AT was present. As seen from the differential inhibitory action of the HOCl scavenger taurine and the peroxynitrite decomposition catalyst FeTPPS, the presence of SHEP effector cells induced apoptosis in SHEP target cells through NO/peroxynitrite signaling, wherease SISO effector cells caused apoptosis through HOCl signaling.

*Statistical analysis: Apoptosis induction mediated by SHEP or SISO effector cells in the presence of 3-AT, as well as the differential inhibitory effect of FeTPPS and taurine were highly significant (p<0.001).*

Supplementary Figure S18

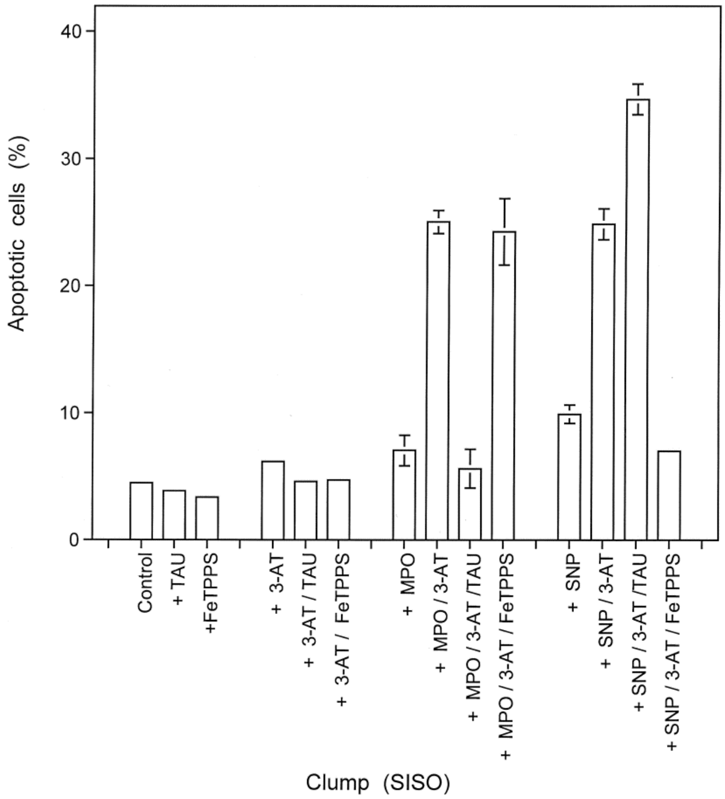


Supplementary Figure S18. MPO or SNP mimick the effector function of SISO or SHEP cells on SISO target cells. Clumps of 1500 SISO cells were seeded and remained either isolated or received either 200 mU/ml MPO or 0.5 mM of the NO donor SNP, in the absence or presence of the indicated inhibitors/scavengers (3-AT: 100 mM; taurine (TAU): 50 mM; FeTPPS (25 µM). After 15 hrs, the percentages of apoptotic cells were determined. Clumps in the absence of effector cells showed background apoptosis induction. The presence of either MPO or SNP caused remarked apoptosis induction, provided the catalase inhibitor 3-AT had been present. As seen from the differential inhibitory action of the HOCl scavenger taurine and the peroxynitrite decomposition catalyst FeTPPS, the presence of MPO caused apoptosis in SISO cells through HOCl signaling, whereas SNP induced apoptosis through NO/peroxynitrite signaling.

*Statistical analysis: Apoptosis induction mediated by MPO in the presence of 3-AT, and its inhibition by taurine, as well as mediated by SNP in the presence of 3-AT and its inhibition by FeTPPS were highly significant (p<0.001).*

Supplementary Figure S19

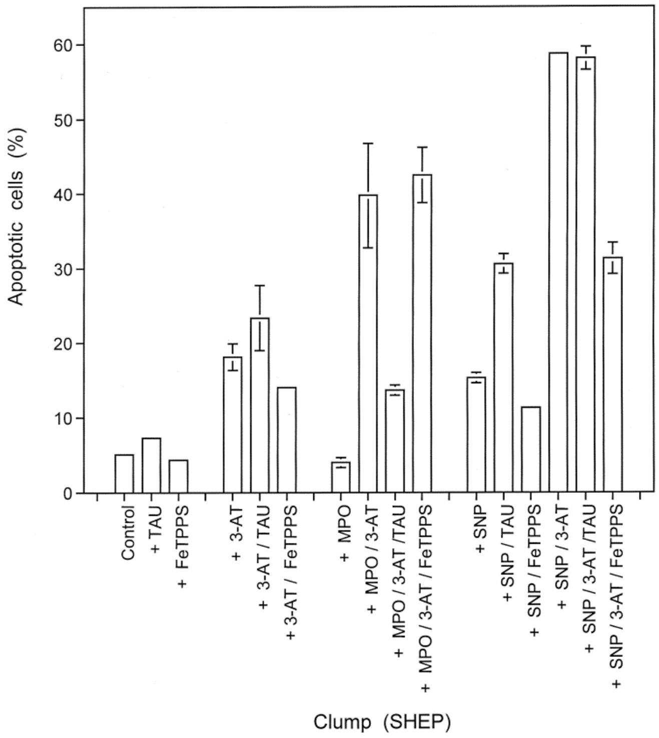


Supplementary Figure S19. MPO or SNP mimick the effector function of SISO or SHEP cells on SHEP target cells. Clumps of 1500 SHEP cells were seeded and remained either isolated or received either 200 mU/ml MPO or 0.5 mM of the NO donor SNP, in the absence or presence of the indicated inhibitors/scavengers (3-AT: 100 mM; taurine (TAU): 50 mM; FeTPPS (25 µM). After 15 hrs, the percentages of apoptotic cells were determined. Clumps in the absence of effector cells showed background apoptosis induction. The presence of either MPO or SNP caused remarked apoptosis induction, provided the catalase inhibitor 3-AT had been present. As seen from the differential inhibitory action of the HOCl scavenger taurine and the peroxynitrite decomposition catalyst FeTPPS, the presence of MPO caused apoptosis in SHEP cells through HOCl signaling, whereas SNP induced apoptosis through NO/peroxynitrite signaling.
*Statistical analysis: Apoptosis induction mediated by MPO in the presence of 3-AT, and its inhibition by taurine, as well as mediated by SNP in the presence of 3-AT and its inhibition by FeTPPS were highly significant (p<0.001).*

apoptosis inducing signaling. The reaction of SHEP and SISO target cells to either MPO or SNP was equally efficient, provided their catalase was inhibited. This finding confirms the central role of catalase for protection towards intercellular ROS/RNS signaling. It also shows that the target function establishes NO/peroxynitrite or HOCl signaling equally well, dependent on the effector molecules available.

Supplementary Figure S20 demonstrates that clumps of SISO cells showed apoptosis induction only i) when they were surrounded by either SISO or SHEP

Supplementary Figure S20

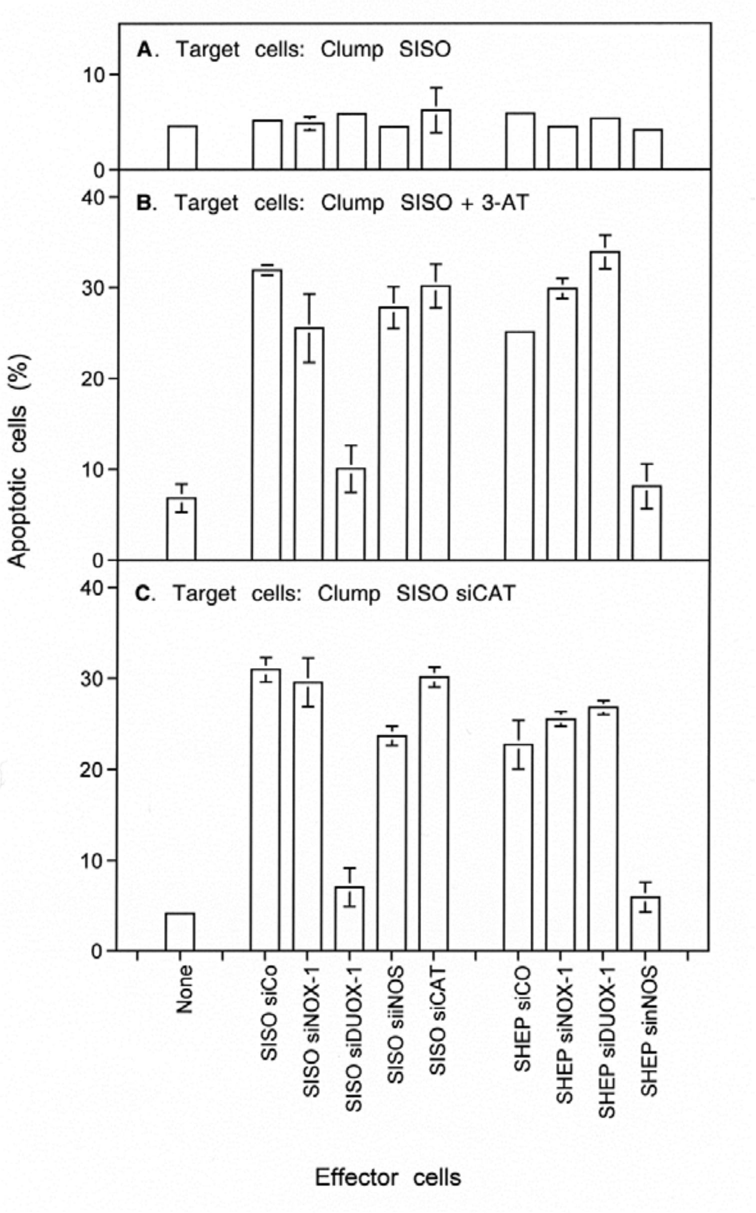


Supplementary Figure S20. The significance of catalase inactivation in target cells and DUOX expression in effector cells. SISO and SHEP cells were transfected with control siRNA (siCO), or siRNAs directed towards the expression of NOX-1 (siNOX-1), DUOX-1 (siDUOX-1), iNOS (siiNOS), nNOS (sinNOS), catalase (siCAT) and

cultivated for 24 hrs. Then, clumps of SISO cells transfected with control siRNA (A, B) or siRNA towards catalase (C) were surrounded by the indicated transfected cells. Assays described under B received 100 mM 3-AT. The percentages of apoptotic cells were determined after 19 hrs. SISO cells were resistant to apoptosis induction despite the presence of effector cells (A). However, when either catalase was inhibited by 3-AT (B) or catalase had been knocked down in the target cells (C), apoptosis induction was efficiently induced in target cells by surrounding SISO or SHEP effector cells. Effector cell function of SISO cells was mediated by DUOX1, whereas the effector function of SHEP cells was due to the action nNOS.
*Statistical analysis: The stimulating effects of effector cells, as well as the interfering effects of siRNA-mediated knockdown of DUOX (SISO cells) and iNOS (SHEP cells) were highly significant ($p<0.001$).*

effector cells and ii) when either their protective catalase was inhibited by 3-AT or had been knocked-down by specific siRNA. Knockdown of DUOX in the effector cells prevented the effector function of SISO cells, but not of SHEP cells. In contrast, the knockdown of NOS prevented the effector function of SHEP cells, but not that of SISO cells. These findings confirm the role of catalase for the protection of target cells and also confirm the role of DUOX-coded peroxidase for the effector function of SISO cells, and of NOS-derived NO for the effector function of SHEP cells. Further controls ensured that the effector function of SISO cells caused indeed HOCl signaling by SISO target cells, whereas the effector function of SHEP caused NO/peroxynitrite signaling (Supplementary Figure S21).

Supplementary Figure S21

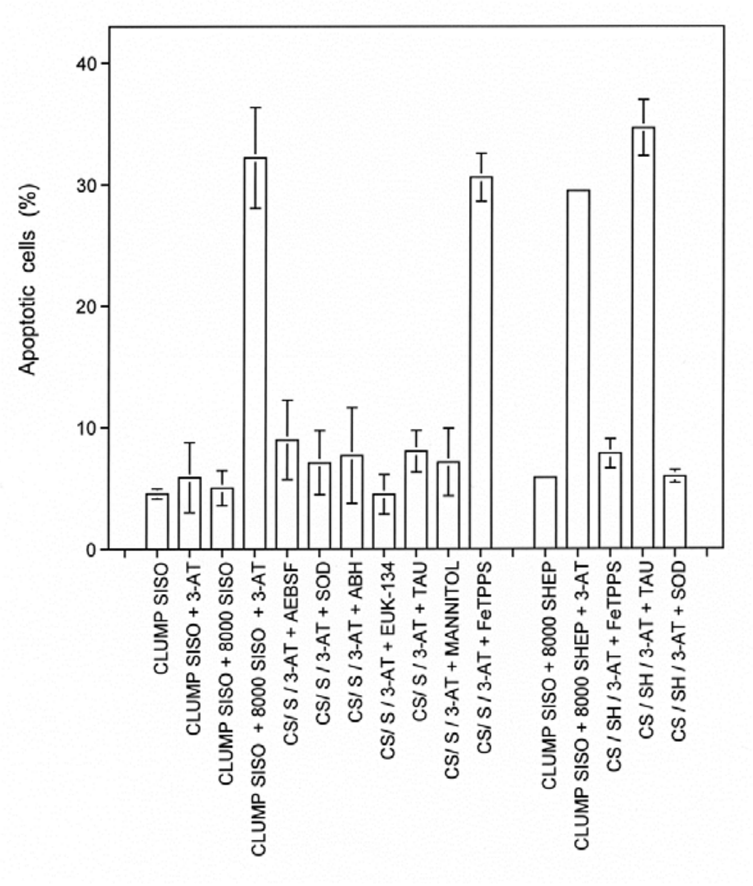


Supplementary Figure S21. Signaling pathways induced by SISO or SHEP effector cells on SISO target cells. Clumps of SISO cells remained without effector cells or were surrounded by either 8000 SISO or SHEP effector cells. Where indicated, the following additions were made: 3-AT (100 mM), AEBSF 100 µM, SOD (100 U/ml), ABG (150 µM), EUK-134 (20 µM), taurine (TAU) (50 mM), mannitol (20 mM), FeTPPS (25 µM). The percentages of apoptotic cells were determined after 19 hrs. Clumps not surrounded by effector cells did not show apoptosis above background, independent of the presence of the catalase inhibitor 3-AT. Target cells surrounded by either SISO or SHEP effector cells showed apoptosis in the presence of 3-AT. Apoptosis induction in SISO target cells plus SISO effector cells, in the presence of 3-AT required superoxide anions (as it was inhibited by AEBSF and SOD), peroxidase (as it was inhibited by ABH), $H_2O_2$ (as it was inhibited by EUK-134), HOCl (as it was inhibited by taurine) and hydroxyl radicals (as it was inhibited by mannitol). This pattern of inhibition is indicative of HOCl signaling. The lack of inhibition by the peroxynitrite decomposition catalyst FeTPPS proofs that NO/peroxynitrite signaling is not contributing to apoptosis induction. When apoptosis induction was induced by SHEP effector cells, signaling was based solely on NO/peroxynitrite signaling, as seen by the strong inhibition by FeTPPS and the lack of inhibition by taurine. The inhibitory effect of SOD points to the generation of peroxynitrite through the interaction between NO and extracellular superoxide anions. *Statistical analysis: The stimulating effect of 3-AT, as well as the inhibitory effects of AEBSF, SOD, ABH,*

*EUK-134, taurine and mannitol in the case of SISO effector cells, and the inhibitory effect of FeTPPS and SOD in the case of SHEP effector cells were highly significant ($p<0.001$).*

The knockdown of various redox-related enzymes in SISO target cells allowed the conclusion that i) the target cell function was determined by active NOX1, but not by NOS or DUOX and ii) that efficient target cell activity required either the presence of a catalase inhibitor or genetic knockdown of protective catalase (Supplementary Figure S22). These basic findings were confirmed for SISO target cells in combination with either SISO or SHEP effector cells (Supplementary Figure S 22) or with the effector molecules MPO or NO donor-derived NO (Supplementary Figure S23). These central findings were confirmed for SHEP target cells, which required active NOX1 for their target function and that responded to the effector function of DUOX released from SISO cells and NO released by SHEP effector cells (Supplementary Figure S24).

Supplementary Figure S22

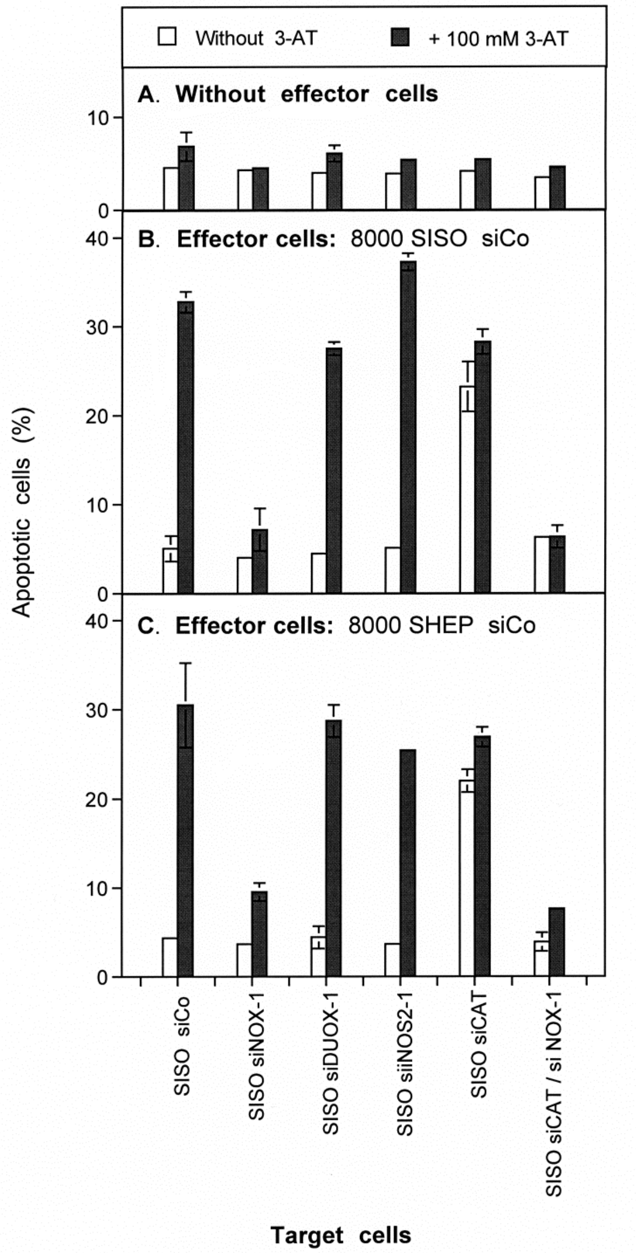


Supplementary Figure S22. Target function of SISO tumor cells is driven by NOX-1 and counteracted by catalase. Clumps of SISO target cells without knockdown (siCo), or siRNA-mediated knockdown of NOX-1, DUOX-1, iNOS2, catalase or catalase plus NOX-1 remained either without effector cells (A) or received 8000 SISO or SHEP effector cells. The effector cells had been transfected with control siRNA. Assays remained free of 3-AT or received 100 mM 3-AT. The percentages of apoptotic cells were determined after 19 hrs. There was no apoptosis induction in the absence of effector cells. SISO and SHEP effector cells mediated a similar degree of apoptosis induction in SISO (siCo) target cells, provided the catalase inhibitor 3-AT was present or catalase had been knocked down in the target cells. Apoptosis induction in the target cells depended on NOX1 and inhibition or knockdown of catalase. *Statistical analysis: The inhibitory effect of siRNA-mediated knockdown of NOX-1 was highly significant (p<0.001).*

Supplementary Figure S23

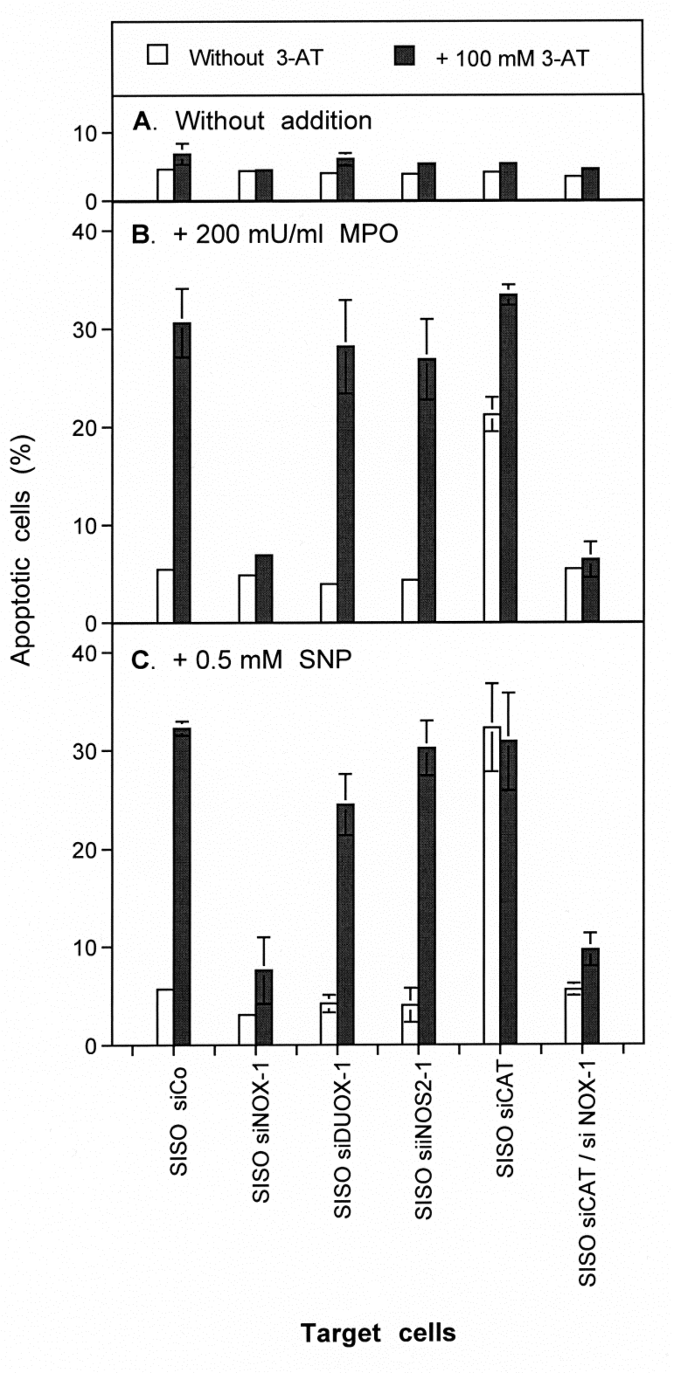


Supplementary Figure S23. SISO target cells respond to either exogenous MPO or SNP, dependent on NOX-1 activitiy and catalase inactivity. SISO target cells analogous to those presented in Supplementary Figure S22 received either 200 mU/ml MPO (B) or 0.5 mM SNP (C), in the absence or presence of 100 mM 3-AT.

Clumps under A only received 3-AT where indicated. The percentages of apoptotic cells were determined after 19 hrs. MPO or SNP completely mimick the effector functions of SISO or SHEP cells, provided the SISO target cells have intact NOX-1 and inactive or knocked-down catalase. *Statistical analysis: The inhibitory effect of siRNA-mediated knockdown of NOX-1 was highly significant (p<0.001).*

Supplementary Figure S24

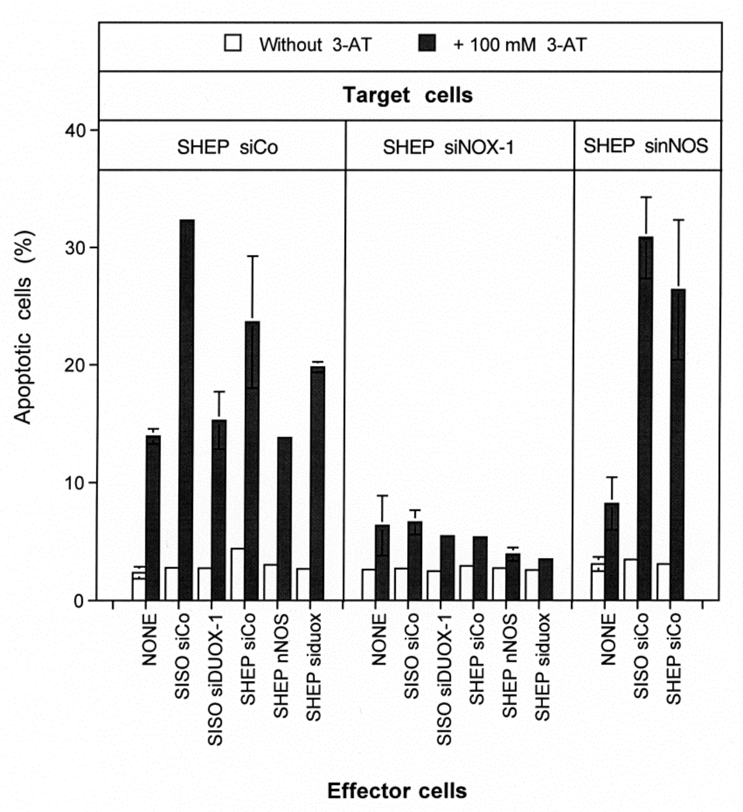


Supplementary Figure S24. The target cell function of SHEP tumor cells is dependent on active NOX-1 and inactive catalase. SHEP and SISO cells were transfected with the indicated siRNAS and incubated for 24 hrs for knockdown of the respective enzymes, Clumps of SHEP control cells (SHEP siCo) or SHEP with knockdown of NOX-1 (SHEP siNOX-19 or nNOS (SHEP sinNOS) remained either without effector cells or were surrounded by SISO or SHEP control cells (siCo) or with the indicated knockdown. Where indicated, 3-AT (100 mM) was added. The percentages of apoptotic cells were determined after 19 hrs.

The data show that the target function of SHEP cells requires active NOX-1 and inactive catalase. The effector function of SISO cells is essentially mediated by DUOX, the effector function of SHEP cells is mediated by nNOS.

*Statistical analysis: The effects of siRNA-mediated knockdown of DUOX-1 on the effector function of SISO cells, the effect of knockdown of nNOS on the effector function of SHEP cells, as well as the inhibitory effect of knockdown of NOX-1 on the target cell function of SHEP cells were highly significant (p<0.001).*

*10. The effect of exogenous NO on the quality of intercellular ROS/RNS signaling of tumor cells with different signaling qualities.*

In line with the antagonistic actions of NO and $H_2O_2$ as characterized in this manuscript, apoptosis induction in tumor cells with preferential HOCl signaling was initially inhibited by exogenous NO, before NO/peroxynitrite signaling was overt at higher concentrations of the NO donor (Supplementary Figures S25, S26A). In contrast, apoptosis induction in NO/peroxynitrite-preferring tumor cells was enhanced by exogenous NO (Supplementary Figure S 26 B).

Supplementary Figure S25

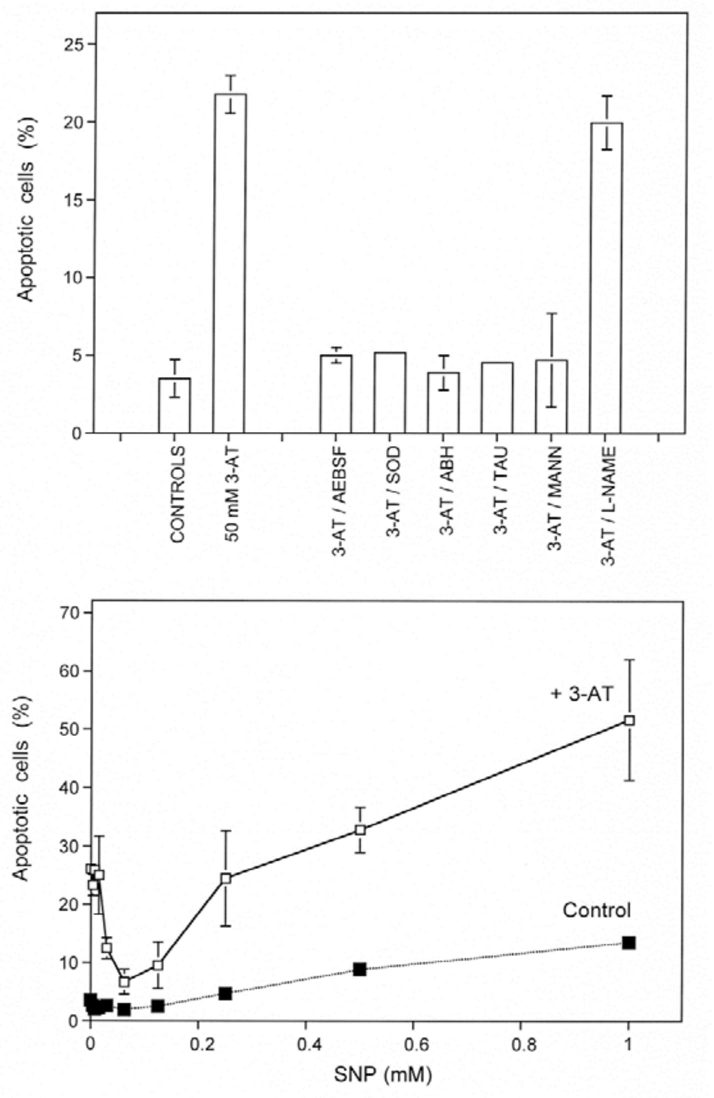

Legend to Supplementary Figure S25. The effect of exogenous NO on the signaling chemistry of PATU 8902 (human pancreatic carcinoma cells). Upper figure: PATU 8902 cells (12 500 cells per assay in 96 well plates, 100 µl medium) received the indicated additions (3-AT: 50 mM; AEBSF: 100 µM; SOD: 100 U/ml, ABH_ 150 µM; taurine (TAU): 50 mM, mannitol: 20 mM, L-NAME. 2.4 mM) and were cultivated for 48 hrs, before the percentages of apoptotic cells were determined. Lower figure: PATU 8902 cells in the absence (control) or presence of 50 mM 3-AT were cultured for 48 hrs in the presence of increasing concentrations of the NO donor SNP.

The data show that 50 mM 3-AT allow apoptosis induction selectively by the HOCl pathway (upper part). Exogenous NO from the NO donor first causes inhibition of 3-AT-mediated apoptosis induction and then resumption of apoptosis at higher concentrations of the NO donor, presumably due to NO/peroxynitrite signaling.

*Statistical analysis: The enhancing effect of 3-AT, as well as the inhibitory effects of AEBSF, SOD, ABH, taurine and mannitol (upper graph), an the effects of SNP (lower graph) were highly significant.*

Supplementary Figure S26

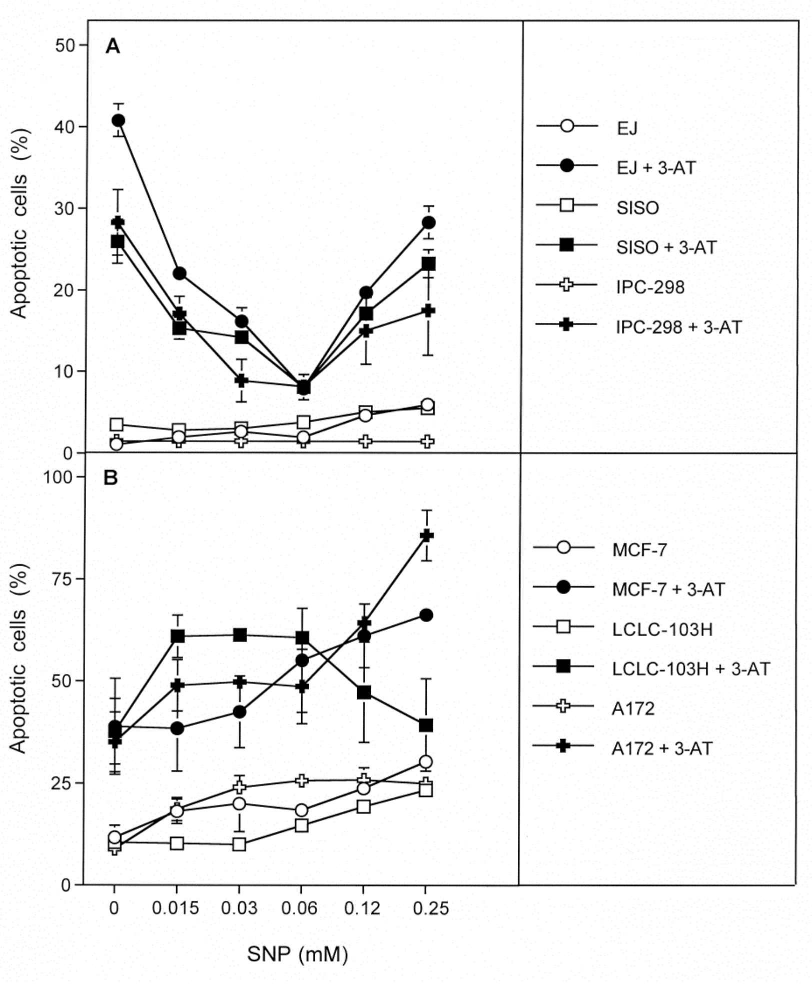

Supplementary Figure S26: Differential effects of exogenous NO on tumor cells with preference for HOCl or NO/peroxynitrite signaling. Human tumor lines were seede in 96 well plates, 12 500 cells per assay, 20ng/ml TGF-beta and 50 mM 3-AT, except for A 172 where 75 mM 3-AT were applied. The assays received the indicated concentrations of the NO donor SNP. Apoptosis induction was determined after 55 hrs, except for SISO cells where measurements were taken at 30 hrs.
Cell lines that showed preferential HOCl signaling in pretesting (A) showed inhibition of 3-AT-mediated apoptosis induction by SNP, followed by resumption of apoptosis induction. Cell lines that preferentially used NO/peroxynitrite signaling according to pretesting showed an increase in apoptosis induction after addition of the NO donor.
*Statistical analysis: the inhibitory effects of SNP under A and the stimulatory effects of SNP under B were highly significant (P<0.001).*

### Supplementary Materials and Methods

*Cell lines*

The non-malignant rat ovary epithelial cell line ROSE (rat ovary surface epithelial cell line) and its H-RAS transformed descendant ROSE A2/5 were a generous gift by Drs C. Sers and R. Schäfer, Berlin, Germany. The human neuroblastoma cell line SHEP was obtained from Dr. J. Roessler, Department of Pediatrics and Adolescent Medicine, University Medical Center, Freiburg.

The human tumor cell lines ICR-37, ICR-39; CAL-51, MCF-7, Fu-OV-1, HCC-1937, SISO, PC-3, LCL-103H, PATU-8902, IPC-298 were purchased from the from DSMZ, Braunschweig, Germany. The human tumor cell lines BG-1, EJ and A 172 were obtained from the American Type Culture Collection. All cell lines were cultivated according to the instructions of the supplyers.

The quantitation of apoptotic cells was performed as described under Material and Methods in the main manuscript.

*Irradiation of cells*

Cells (200,000 cells/ml in suspension) were irradiated using a Compagnie Oris (Gif-sur-Yvette, France) IBL 437 Cs137 gamma source (40 mGy/s). Immediately after irradiation, the cells were centrifuged, resuspended in fresh medium and incubated in suspension at 200000 cells/ml for 1 h to allow maximal enhancement of superoxide anion production [13]. The cells were then centrifuged and seeded at the density described in the respective figure legends.

*Knockdown by treatment with specific small interfering ribonucleic acids (siRNAs).*

All siRNAs were obtained from Qiagen (Hilden, Germany).

*The following siRNAs were used:*

Control siRNA which does not affect any known target in human and murine cells (siCo):

sense: r(UUCUCCGAACGUGUCACGU)dTdT,

antisense: CGUGACACGUUCGGAGAA)dTdT;

**SiRNAs directed towards human targets**:

custom-made siRNA directed towards NADPH oxidase-1 variant a (**siNOX1-a)**: target sequence: CCG ACA AAT ACT ACT ACA CAA

sense: r(GAC AAA UAC UAC UAC ACA A)dTdT,

antisense: r(UUG UGU AGU AGU AUU UGU C)dGdG;

custom made siRNA directed towards dual oxidase-1 variant a (**siDUOX1-a)**: target sequence: CCA GTC TAA CAC CAC AAC TAA,

sense: r(AGU CUA ACA CCA CAA CUA A)dTdT,

antisense: r(UUA GUU GUG GUG UUA GAC U)dGdG;

custom-made siRNA directed towards inducible NO synthase-2 (**siiNOS2)**: target sequence: CTG GGC CGT GCA AAC CTT CAA

sense r(GGG CCG UGC AAA CCU UCA A)dTdT;

antisense: r(UUG AAG GUU UGC ACG GCC C)dAd;

custom-made siRNA directed towards neuronal NO synthase (**sinNOS)**; target sequence: CAC AAG TGT GTC GAT CTT AGA, sense: r(CAA GUG UGU CGA UCU UAG A)dTdT; antisense: r(UCU AAG AUC GAC ACA CUU G)dTdG;

siRNA directed towards catalase **(siCAT)** (Hs_CAT_4_HP siRNA); target sequence: CCG GAT CTC ACT TGG CGG CAA; sense: r(GGAUCUCACUUGGCGGCAA)dTdT; antisense: r(UUGCCGCCAAGUGAGAUCC)dGdG;

SiRNAs were dissolved in suspension buffer supplied by Qiagen at a concentration of 20 µM. Suspensions were heated at 90°C for 1 minute, followed by incubation at 37°C for 60 minutes. Aliquots were stored at −20°C.

Before transfection, 88 µl of medium without serum and without antibiotics were mixed with 12 µl Hyperfect solution (Qiagen) and the required volume of specific siRNA or control siRNA to reach the desired concentration of siRNA during transfection (the standard concentration of siRNA was 24 nM for MKN-45 cells and 10 nM for 208Fsrc3 cells). The mixture was treated by a Vortex mixer for a few seconds and then allowed to sit for 10 minutes. It was then gently and slowly added to 300,000 MKN-45 cells in 1 ml RPMI-1640 medium containing 10% FBS and antibiotics (12-well plates) or to 200,000 208Fsrc3 cells/well in 2.3 ml medium supplemented with 5% FBS and antibiotics (6-well plates). The cells were incubated at 37°C in 5% $CO_2$ for 24 hours. Transfected cells were centrifuged and resuspended in fresh medium at the required density before use.

*Determination of the efficiency of siRNA-mediated knockdown.* The siRNA transfection system as described above had been optimized to allow a reproducible transfection efficiency of more than 95% of the cells and to avoid toxic effects (Bauer, unpublished data).

The efficiency of knockdown by siNOX1, DUOX1and catalase was based on functional quantitative assays and was more than 90 % [16, 4]. The efficiency of knockdown of iNOS, nNOS, was proven through complete block of apoptosis after knockdown, which was abrogated by supplementation with an NO donor [15].

**Supplementary References**

1 Scheit K and Bauer G. Direct and indirect inactivation of tumor cell protective catalase by salicylic acid and anthocyanidins reactivates intercellular ROS signaling and allows for synergistic effects. Carcinogenesis 36 (2015) 400-411.

2 Bauer G and Zarkovic N. Revealing mechanisms of selective, concentration-dependent potentials of 4-hydroxy-2-nonenal to induce apoptosis in cancer cells through inactivation of membrane-associated catalase. Free Radic. Biol. Med. 81: 128-144, 2015.

3 Bauer G. Autoamplificatory singlet oxygen generation sensitizes tumor cells for intercellular apoptosis-inducing signaling. Mechanisms of Ageing and Development 172: 59-77, 2018; doi:org/10.1016/j.mad.2017.11.005.

4 Di Mascio P, Bechara EJH, Medeiros MHG, Briviba K and Sies H: Singlet molecular oxygen production in the reaction of peroxynitrite with hydrogen peroxide. FEBS Lett *355*: 287-289, 1994.

5 Bauer G. The synergistic effect between hydrogen peroxide and nitrite, two long-lived molecular species from cold atmospheric plasma, triggers tumor cells to induce their own cell death. Redox Biol, 26, 101291 ( 2019). https://doi.org/10.1016/j.redox.2019.101291

6 Böhm B, Heinzelmann S, Motz M, Bauer G. Extracellular localization of catalase is associated with the transformed state of malignant cells. Biol. Chem. 396 (2015) 1339-1356.

7 Riethmüller M, Burger N, Bauer G: Singlet oxygen treatment of tumor cells triggers extracellular singlet oxygen generation, catalase inactivation and reactivation of intercellular apoptosis-inducing signaling. Redox Biol, 6 (2015) 157-168.

8 Escobar JA, Rubio A, Lissi EA. SOD and catalase inactivation by singlet oxygen and peroxyl radicals. Free Radic. Biol. Med. 1996;20: 285-290.

9 Kim YK, Kwon OJ, Park J-W. Inactivation of catalase and superoxide dismutase by singlet oxygen derived from photoactivated dye. Biochimie 2001;83: 437-444.

10 Bechtel W and Bauer G: Catalase protects tumor cells against apoptosis induction by intercellular ROS signaling. Anticancer Res *29*: 4541-4557, 2009

11 Bechtel W and Bauer G: Modulation of intercellular ROS signaling of human tumor cells. Anticancer Res *29*: 4559-4570, 2009

12 Bauer G. Low dose irradiation enhances specific signaling components of intercellular reactive oxygen-mediated apoptosis induction. J. Phys. Conf. ser. 261 012001, 2011

13 Temme J and Bauer G: Low-dose gamma irradiation enhances superoxide anion production by nonirradiated cells through TGF-β1-dependent bystander signaling. Rad. Res. 179: 422-432, 2013.

14 Pottgiesser S, Heinzelmann S and Bauer G. Intercellular HOCl-mediated apoptosis induction in malignant cells: interplay between NOX1-dependent superoxide anion generation and DUOX-related HOCl-generating peroxidase activity. Anticancer Res. 35: 5927-5943, 2015.

15 Bauer G. Central signaling elements of intercellular reactive oxygen/nitrogen species-dependent induction of apoptosis in malignant cells. Anticancer Res 37 (2017 a) 499-514.
